# GLP-1 analogs restore inflammatory, mitochondrial and intercellular signaling networks in the *Snca^G51D/G51D^* knock-in mouse model of Parkinson’s disease

**DOI:** 10.64898/2026.05.18.726024

**Authors:** Bhupesh Vaidya, Yan Li, YoungDoo Kim, Cameron Osterman, Jean-Pierre Revelli, Huda Y. Zoghbi

## Abstract

Parkinson’s disease (PD) is a neurodegenerative disorder characterized by a prolonged prodromal stage that culminates in motor deficits. Current PD therapies primarily alleviate symptoms, underscoring the need for disease-modifying strategies. Glucagon-like peptide-1 (GLP-1) analogs showed early promise as candidate disease modifiers, but recent clinical results have been inconsistent, and their mechanism of action remains poorly defined. Here, we employed our *Snca^G51D/G51D^* knock-in mouse model to investigate the effects of subcutaneously administered GLP-1 analogs, semaglutide and lixisenatide. Both analogs reversed motor and non-motor deficits and reduced gliosis and detergent-insoluble α-synuclein. Bulk and single-nuclei transcriptomics together with CellChat-based intercellular communication analysis revealed that GLP-1 analogs normalize early striatal mitochondrial and inflammatory dysregulation and restore neuregulin (NRG) and neurexin (NRXN) signaling networks to wild-type levels. Treatment was effective when initiated either before or shortly after symptom onset, defining an early therapeutic window for GLP-1 analog therapy in PD.

## Introduction

Parkinson’s disease (PD) is the second most prevalent neurodegenerative disorder in the world, with the number of patients expected to cross 25 million by 2050 (*1*). PD is characterized by prodromal deficits in the form of constipation and loss of olfaction that appear years before the onset of motor deficits and overt neurodegeneration in the substantia nigra and striatum (*2, 3*). Despite recent advances in symptomatic management of PD, current therapies are largely aimed at the restoration of dopamine levels to reduce motor deficits without any effect on the underlying pathology to halt or reverse disease progression (*4*). Consequently, there is a need to identify new disease-modifying therapeutics that target early pathogenic mechanisms to stop the overall course of PD or at least delay its progression.

Amongst the emerging candidate molecules, Glucagon-like peptide-1 (GLP-1) analogs have generated considerable interest as potential disease-modifying strategies (*5*). These drugs were originally developed as anti-obesity and anti-diabetic drugs (*6*), but later showed some improvements in toxin models such as 1-methyl-4-phenyl-1,2,3,6-tetrahydropyridine (MPTP) and 6-OHDA of PD (*7, 8*). These studies paved the way for the clinical investigation, where a Phase II clinical trial for the GLP-1 analog, lixisenatide, decreased progression of motor disability relative to placebo, even after 12 months (*5*). However, other GLP-1 analogs have yielded inconsistent results, with exenatide, for example, recently failing to demonstrate sufficient clinical benefits in PD patients (*9*). In parallel, trials with other drugs of this class, including semaglutide, are currently ongoing, reflecting a continued interest in GLP-1 analogs as a potential therapy and a need to clearly define their mechanism of action (*10*).

Although interest in GLP-1 analogs is growing, considerable uncertainty remains regarding their beneficial effects and mechanisms of action in PD. The main reason is that prior studies have relied heavily on toxin-based models, which do not recapitulate the progressive, α-synuclein-driven pathology observed in most PD patients (*11*). As a result, it is also unclear whether GLP-1 analogs act on the core disease mechanisms and drivers or merely mitigate downstream consequences of overt neurodegeneration. Importantly, disruption of intercellular communication has emerged as an unresolved question in PD pathogenesis, with recent evidence suggesting these alterations are a key event in disease progression rather than a secondary consequence. Therefore, it is important to determine whether GLP-1 analogs can normalize these cell-cell communication pathways and restore network homeostasis in PD, a question that remains unanswered.

To address these questions and characterize the mechanistic effects of GLP-1 analogs, we leveraged our recently developed *Snca^G51D/G51D^*PD mouse model (*12*), which recapitulates the prodromal symptoms and clinical progression of PD. It further provided an opportunity to test whether GLP-1 analogs can intervene at early stages of PD to reverse the disease trajectory rather than simply alleviate symptoms. Given their ongoing clinical investigation (*5, 10*), we tested two GLP-1 receptor analogs, *viz*., semaglutide and lixisenatide, to assess class-wide and compound-specific effects in PD.

We performed a comprehensive multi-level analysis integrating behavioral phenotyping, molecular characterization, transcriptomic analysis, and ligand-receptor interaction modeling to study the effect of these GLP-1 analogs in PD. We find that GLP-1 analog treatment rescued both the prodromal and motor symptoms of PD. Moreover, it reduced gliosis and insoluble alpha-synuclein levels, rescued early mitochondrial deficits, and normalized intercellular signaling networks, including the neuregulin (NRG) and neurexin (NRXN) pathways, in *Snca^G51D/G51D^* mice. Overall, we establish the mechanism of action of GLP-1 analogs in PD, with translational implications for patients, especially when initiated early or shortly after symptom onset.

## Results

### GLP-1 analogs rescue early olfaction and constipation phenotypes in *Snca^G51D/G51D^* mice

Prodromal symptoms, such as constipation and loss of olfaction, precede the onset of motor deficits in PD patients (*13*). In contrast to most available PD models, the *Snca^G51D/G51D^* mouse model exhibits early prodromal symptoms and subsequently develops motor deficits (*12*). Therefore, it captures a clinically relevant window for early intervention that is otherwise inaccessible using conventional toxin-based or overexpression models and is important for disease-modifying drug discovery. Using this mouse model, we tested whether treatment with GLP-1 analogs could ameliorate these early clinical manifestations. We dosed the mice subcutaneously with lixisenatide or semaglutide, starting at 4 months of age before the onset of prodromal symptoms, and assessed their effects at 6 months (Fig. 1A). Doses were determined in accordance with the prior literature and validated using a pilot experiment (Supplementary Fig. 1). The selected doses were well tolerated, with none of the GLP-1-treated WT mice experiencing greater than 10% reduction in body weight.

**Fig. 1:**
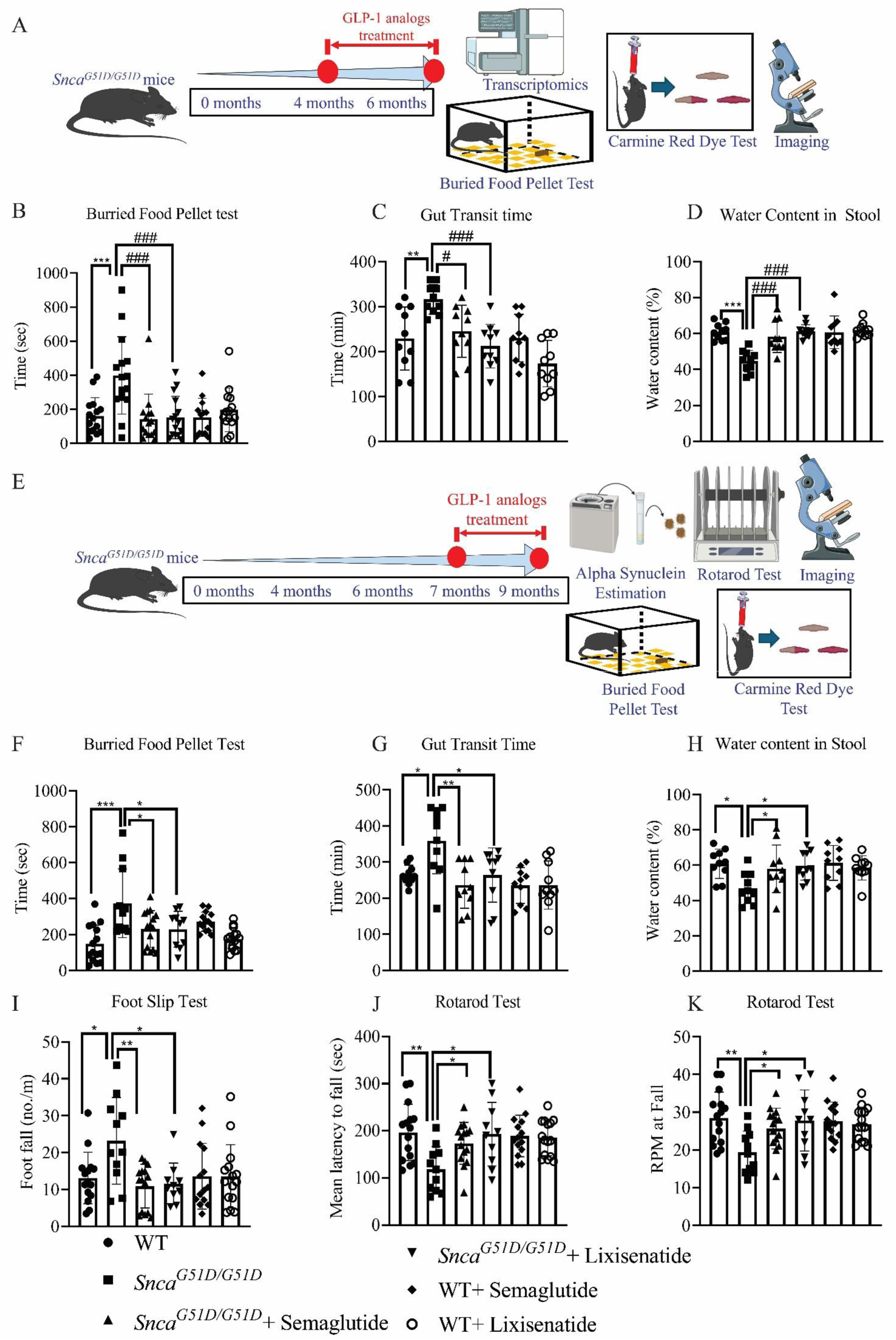
GLP-1 analogs treatment rescues prodromal symptoms of PD in *Snca^G51D/G51D^* mice at 6 and 9 months. (A) Schematic of GLP-1 analog treatment to *Snca^G51D/G51D^* mice at 6 months, followed by behavioral assays, transcriptomics, and confocal microscopy. (B) Buried Food Pellet Test (C) Gut Transit Time following oral gavage of carmine red dye and (D) Water content in stool of mice compared across groups at 6 months. (E) Schematic of GLP-1 analog treatment to *Snca^G51D/G51D^* mice at 9 months, followed by behavioral assays, alpha synuclein estimations, and confocal microscopy. (F) Buried Food Pellet Test (G) Gut Transit Time following oral gavage of carmine red dye (H) Water content in stool of mice (I) Parallel Rod Floor Test (J) Time Spent on Day 3 of Rotarod Test and (K) RPM at which mice fell from the accelerating rotarod on Day 3 at 9 months. Individual data points represent biological replicates. Results are expressed as mean ± SD. ***p<0.001, **p<0.01, *p<0.05.

At 6 months of age, *Snca^G51D/G51D^* mice exhibited a profound olfactory deficit compared to WT mice, taking a significantly longer time to locate a buried food pellet. However, *Snca^G51D/G51D^* mice that received GLP-1 analogs, *viz.*, semaglutide and lixisenatide, showed reduced latencies to locate the buried pellet and restored performance to levels comparable to those of WT animals. Moreover, GLP-1 analogs did not have any deleterious effects on olfactory performance in WT animals (Fig. 1B).

Constipation is another well-recognized prodromal symptom of PD, often manifesting years before the onset of motor deficits in patients (*14*), and has been previously reported in *Snca^G51D/G51D^* mice (*12*). Consistent with these findings, *Snca^G51D/G51D^* mice in our cohort also exhibited delayed expulsion of carmine red dye following oral gavage and reduced stool water content compared to WT mice (Fig. 1C, D). In contrast, treatment with GLP-1 analogs shortened gut transit time and increased stool water content in *Snca^G51D/G51D^* mice, normalizing the gastrointestinal function toward WT-like behavior. (Fig. 1C-D). In contrast with clinical studies involving GLP-1 analogs, we did not identify any constipation-related side effects of lixisenatide or semaglutide (10 and 30 nmol/kg, respectively) in WT mice (*15, 16*).

### Late-stage GLP-1 analog treatment reverses non-motor phenotypes and prevents motor deficits in *Snca^G51D/G51D^* mice

We next asked whether the GLP-1 analogs remain efficacious even after the initial appearance of disease symptoms. To test that, we employed a late-intervention paradigm in which a separate cohort of mice was dosed after the initiation of symptoms for two months, starting at 7 months of age, with motor and non-motor outcomes assessed at 9 months (Fig. 1E). Given that non-motor symptoms often worsen with PD progression (*16, 17*), both motor and non-motor parameters were evaluated at this time point.

We previously reported olfactory and gastrointestinal deficits in *Snca^G51D/G51D^* mice at 6 months (*12*). Here, we confirm that these deficits persist at 9 months of age. However, to our surprise, late treatment with GLP-1 analogs significantly reduced latency to locate the buried food pellet and normalized olfactory function, even at 9 months (Fig. 1F-H). Interestingly, GLP-1 analog administration also brought the gut transit time and stool water content to WT levels, demonstrating a complete reversal of non-motor behavioral phenotypes.

Finally, we assessed the effects of GLP-1 analogs on motor function across treatment groups by performing the parallel rod floor test and rotarod assays. The parallel rod floor test is used to detect subtle changes in motor coordination and serves as a more sensitive readout of skilled locomotor behavior (*18*). Accelerated rotarod helps detect global motor coordination and balance, and is commonly used to detect overt motor dysfunctions (*19, 20*). Treatment with GLP-1 analogs in *Snca^G51D/G51D^* mice prevented the onset of motor deficits and normalized outcomes in both assays. Moreover, WT mice treated with semaglutide or lixisenatide did not exhibit worsening performance in any behavioral assay, arguing against deleterious effects on basal functions in this paradigm (Fig. 1I-K). These results demonstrate that GLP-1 analogs may be effective in treating PD even when administered after symptom onset.

We also examined common metabolic parameters like cholesterol, insulin, triglycerides, and glucose in the plasma of 9-month-old GLP-1-treated mice (Supplementary Fig. 2). However, aside from reductions in total cholesterol, most of these parameters remained consistent across groups at the doses selected in our study, suggesting that therapeutic benefits are largely mediated by central nervous system effects of these drugs.

### GLP-1 analog treatment reverses early and late-stage region-specific gliosis in *Snca^G51D/G51D^* mice

Neuroinflammation is a prominent feature of PD and is increasingly viewed as an active contributor to disease progression rather than a passive byproduct (*21*). *Snca^G51D/G51D^* mice exhibit gliosis at 12 months (*12*); however, these changes have not been examined at an earlier time point. Therefore, we phenotypically characterized neuroinflammatory changes across groups at 6- and 9-month time points and assessed the effect of GLP-1 analog treatment using immunofluorescence for GFAP (astrogliosis) and Iba1 (microgliosis). This region-focused approach was based on evidence from previous studies, which have revealed that PD pathology first appears in the olfactory bulb and associated cortical regions, such as the piriform cortex, before affecting the substantia nigra and striatum (*22, 23*).

Relative to controls, *Snca^G51D/G51D^* mice exhibited the largest increase in GFAP and Iba1 signal intensity (markers of astrogliosis and microgliosis, respectively) in the olfactory bulb (Fig. 2A), while the piriform cortex and substantia nigra only exhibited elevated GFAP expression at 6 months. Following GLP-1 analog treatment, there was a complete reversal of increased GFAP expression in the olfactory bulb, piriform cortex, and substantia nigra, as well as decreased microgliosis in the olfactory bulb.

**Fig. 2:**
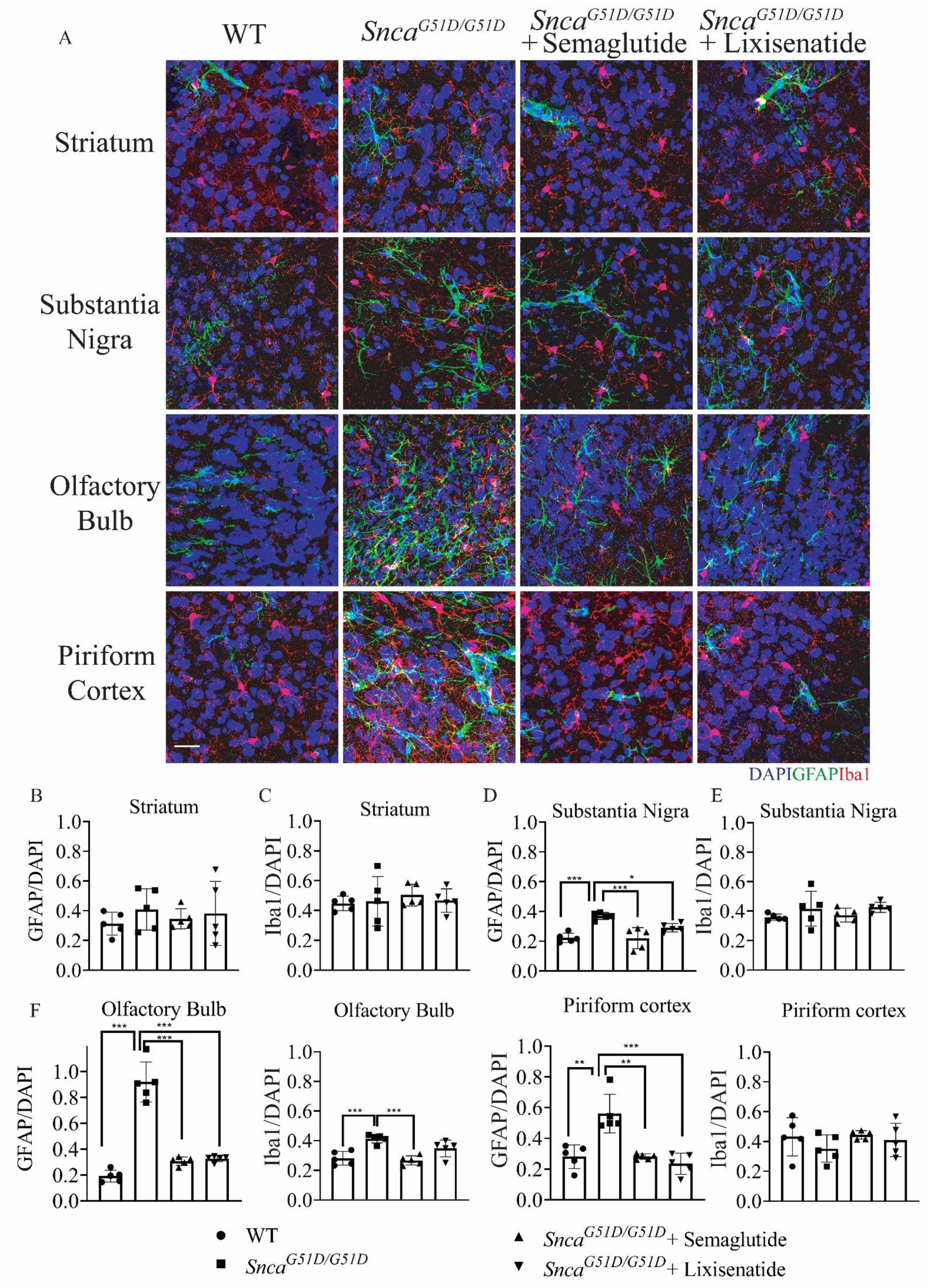
GLP-1 analogs treatment rescues gliosis across different brain regions at 6 months. (A) Representative images of astrocyte (GFAP) and microglia (Iba1) staining in the striatum, substantia nigra, olfactory bulb and piriform cortex of different groups. (B-I) Quantification graphs of Iba1 and GFAP signal intensity in the striatum, substantia nigra, olfactory bulb and piriform cortex of animals belonging to different groups. Individual data points represent biological replicates. Results are expressed as mean ± SD. ***p<0.001, **p<0.01, *p<0.05.

On the other hand, GFAP and Iba1 expression levels in the striatum were not significantly different from those of the WT mice at 6 months, indicating relative sparing at this age (Fig. 2B-I). This is consistent with previous observations in PD patients, which demonstrate that the striatum is resistant to early phenotypic changes, exhibiting symptoms only after 80% of dopaminergic terminals in the striatum are affected (*24*).

Strikingly, two months of GLP-1 analog administration after symptom onset also reduced GFAP levels across all brain regions, and Iba1 levels in the olfactory bulb and piriform cortex of 9-month-old *Snca^G51D/G51D^* mice (Fig. 3A-I). Importantly, administration of lixisenatide and semaglutide to WT mice did not exert any deleterious effects on basal GFAP or Iba1 levels at either time point, demonstrating disease-associated normalizations to WT levels without any nonspecific effects on glial function (Supplementary Fig. 3A, B).

**Fig. 3:**
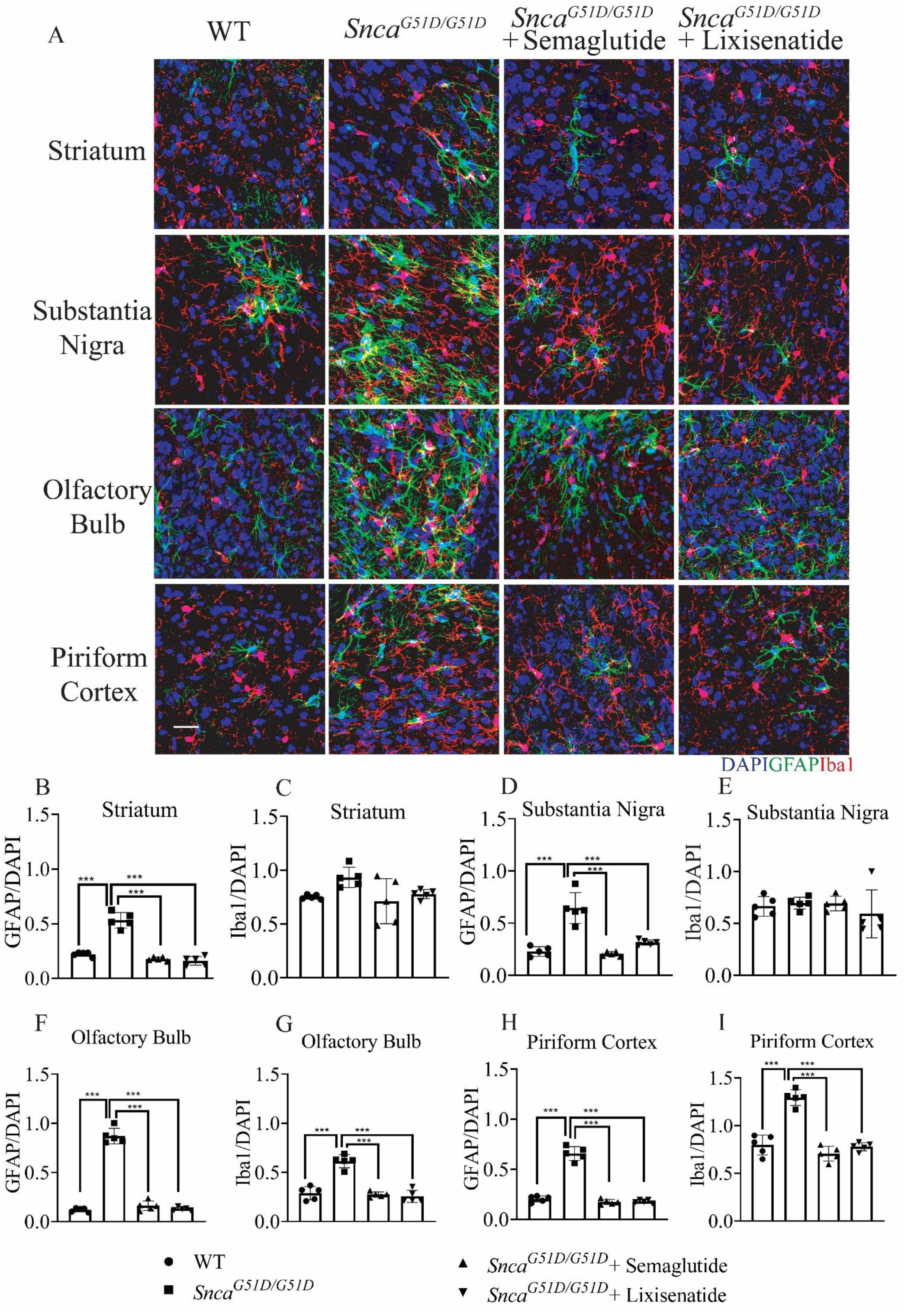
GLP-1 analogs treatment rescues gliosis across different brain regions at 9 months. (A) Representative images of astrocyte (GFAP) and microglia (Iba1) staining in the striatum, substantia nigra, olfactory bulb and piriform cortex of diffe rent groups. (B-I) Quantification graphs of Iba1 and GFAP signal intensity in the striatum, substantia nigra, olfactory bulb and piriform cortex of animals belonging to different groups. Individual data points represent biological replicates. Results are expressed as mean ± SD. ***p<0.001

### GLP-1 analogs reduce insoluble alpha synuclein levels across brain regions in *Snca^G51D/G51D^* mice

Alpha-synuclein accumulation is a major pathological event in PD (*25*). Previous studies in patients with G51D mutations, as well as in *Snca^G51D/G51D^* mice, have highlighted the formation of toxic insoluble alpha-synuclein aggregates, resulting in PD progression (*12, 26*).

Therefore, to assess whether the GLP-1 analog treatment affects α-synuclein aggregation levels, we quantified soluble and insoluble α-synuclein fractions in the midbrain, olfactory bulb, striatum, and piriform cortex of 9-month-old mice from different groups. We previously showed that *Snca^G51D/G51D^* mice exhibit increased accumulation of insoluble α-synuclein in multiple brain regions (*12*). Two months of GLP-1 analog treatment significantly reduced insoluble alpha-synuclein levels across different brain regions. This reduction brought levels back to nearly WT levels and was accompanied by a modest shift toward the soluble fraction, suggesting a decreased aggregation burden rather than altered total protein levels. Notably, the olfactory bulb, one of the earliest sites of pathology, also showed reduced insoluble α-synuclein, which aligns with the rescue of non-motor phenotypes at 9 months of age (Fig. 4A-I).

**Fig. 4:**
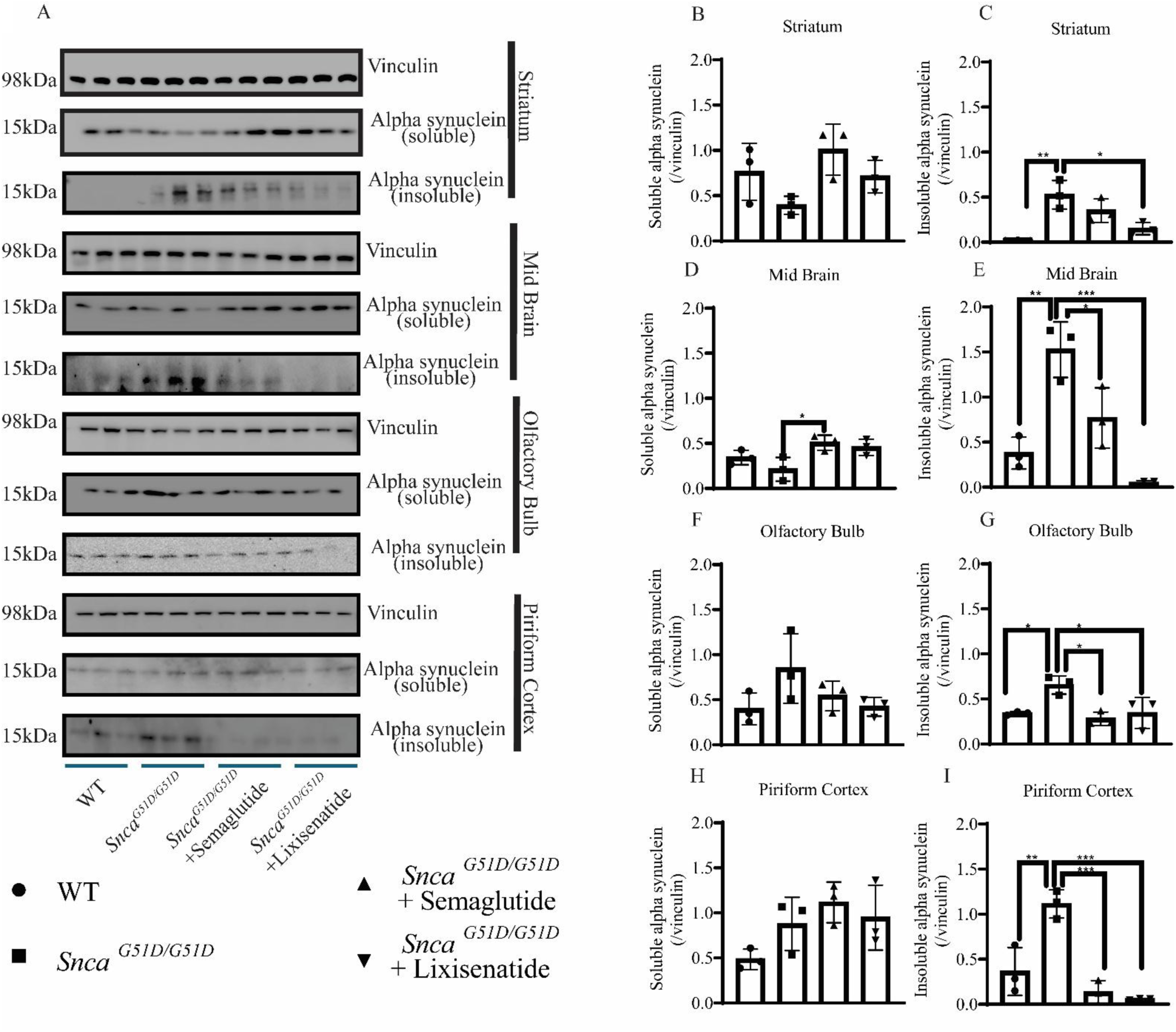
GLP-1 analog treatment reduces insoluble alpha-synuclein aggregates across brain regions at 9 months. (A) Representative immunoblots showing expression of soluble and insoluble alpha-synuclein fractions across brain regions. (B-I) Quantification for the expression of alpha-synuclein levels in the striatum, midbrain, olfactory bulb, and piriform cortex across different treatment groups. Individual data points represent biological replicates. Results are expressed as mean ± SD. ***p<0.001, **p<0.01, *p<0.05.

### GLP-1 analogs reverse early mitochondrial transcriptional changes in the prodromal striatum of *Snca^G51D/G51D^* mice

Our immunofluorescence data indicated that the striatum remained histologically intact at 6 months, despite the onset of prodromal symptoms in *Snca^G51D/G51D^* mice, with astrogliosis becoming evident only at 9 months. This histologically preserved window provides a critical opportunity to capture the earliest molecular changes preceding neuroinflammation and motor dysfunction. Identifying transcriptional dysregulation at this presymptomatic stage is particularly important, as these early molecular alterations may underlie disease initiation and represent targets for intervention before irreversible pathological changes occur.

This window also allows testing whether GLP-1 analog treatment could reverse these early, disease-driving transcriptomic alterations rather than affect downstream consequences. We therefore performed bulk RNA sequencing in the striatum of WT, *Snca^G51D/G51D^*, and GLP-1 analog-treated cohorts (semaglutide or lixisenatide) (Fig. 5A).

**Fig. 5:**
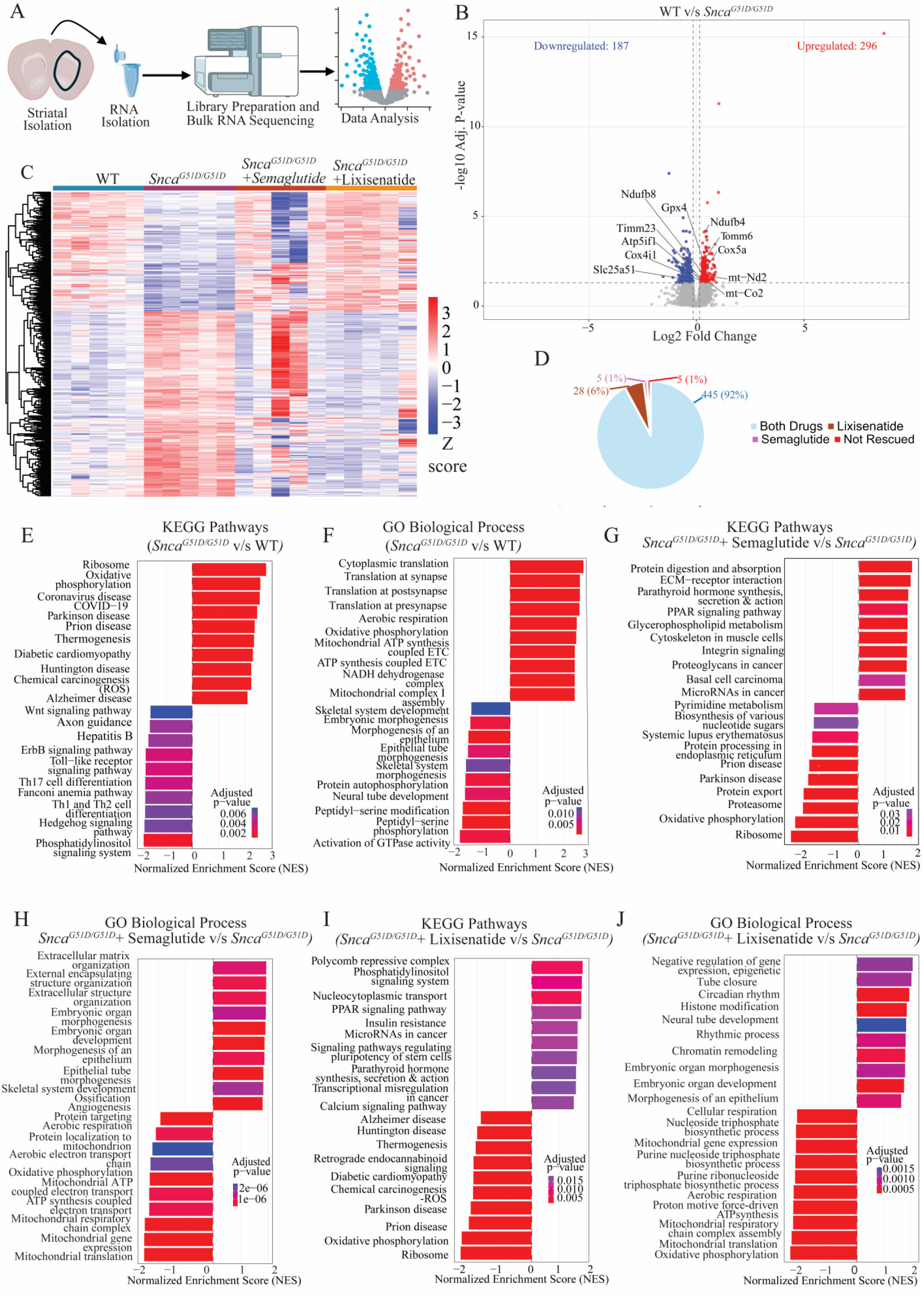
GLP-1 analogs treatment reverses transcriptomic dysregulation in the striatum of *Snca^G51D/G51D^* mice at 6 months. (A) Schematic of the experimental design involving striatal RNA isolation, library preparation, sequencing and data analysis (B) Volcano plots of DEGs between WT and *Snca^G51D/G51D^*mice. (C) Heatmap depicting DEGs in *Snca^G51D/G51D^* striatum, which are rescued in the GLP-1-treated groups. (D) Pie chart indicating rescue in the corrected genes by GLP-1 analogs. (E) KEGG pathway analysis of DEGs in *Snca^G51D/G51D^* mice compared to WT mice (F) Gene Ontology analysis of DEGs in *Snca^G51D/G51D^* mice compared to WT mice (G) KEGG pathway analysis of DEGs in *Snca^G51D/G51D^* + Semaglutide mice compared to *Snca^G51D/G51D^* mice. (H) Gene Ontology analysis of DEGs in *Snca^G51D/G51D^* + Semaglutide mice compared to *Snca^G51D/G51D^*mice (I) KEGG pathway analysis of DEGs in *Snca^G51D/G51D^* + Lixisenatide mice compared to *Snca^G51D/G51D^* mice. (J) Gene Ontology analysis of DEGs in *Snca^G51D/G51D^* + Lixisenatide mice compared to *Snca^G51D/G51D^* mice

Principal component analysis revealed separation driven primarily by genotypes and drug treatments, with GLP-1-treated *Snca^G51D/G51D^*mouse samples showing partial convergence towards WT sample profiles, suggesting treatment-dependent normalization of transcriptional states (Supplementary Fig. 4A). In-depth analysis further revealed 483 differentially expressed genes in the *Snca^G51D/G51D^* group, with 296 upregulated and 187 downregulated compared with WT mice (Fig. 5B). Notably, several of the prominently altered transcripts encoded the components of the mitochondrial electron transport chain or associated biological processes, including subunits of complex I-V, oxidative stress, mitochondrial import, and ribosomal proteins (Fig. 5B and Supplementary Fig. 4B).

Hierarchical clustering of DEGs in a heatmap revealed that the disease-associated transcriptional changes were broadly shifted toward WT following treatment with GLP-1 analogs (Fig. 5C). Quantification of this rescue pattern revealed a striking normalization of the disease signatures by GLP-1 analog treatment. Specifically, the altered expression of most genes in *Snca^G51D/G51D^* mice was rescued by both GLP-1 analogs (445 genes, 92%), with an additional, smaller subset of 28 genes (6%) selectively rescued by lixisenatide and an additional 5 genes (1%) rescued by semaglutide (Fig. 5D).

Gene Set Enrichment Analysis further underscored the centrality of mitochondrial dysfunction in this model before the onset of astrogliosis in the striatum. In WT versus *Snca^G51D/G51D^* striatum, KEGG pathway and gene ontology analyses identified oxidative phosphorylation, electron transport chain activity, and related PD-linked pathways among the most significantly altered in *Snca^G51D/G51D^* mice (Fig. 5E-F). In semaglutide-treated mice, pathways related to oxidative phosphorylation, mitochondrial translation, and mitochondrial ATP synthesis coupled electron transport were among the most strongly normalized (Fig. 5G-H), while lixisenatide also normalized PD disease signatures and pathways associated with mitochondrial functions, including oxidative phosphorylation, mitochondrial translation, and mitochondrial respiratory chain complex assembly (Fig. 5I-J). Although there was no overt gliosis in the striatum at 6 months, GLP-1 analogs also normalized the expression of inflammatory and immune-related genes, with rescue observed across cytokine, chemokine, immune, and glial-associated pathways (Supplementary Fig. 4C-D). These findings suggest that GLP-1 receptor activation coordinately restores both metabolic and inflammatory programs during early PD.

### Single-nuclei RNA sequencing reveals cell-type–specific vulnerability in *Snca^G51D/G51D^* mice, which is rescued by GLP-1 analog treatment

To resolve the cellular basis of early transcriptional alterations and delve deeper into the alteration in signaling responsible for changes seen in bulk RNA sequencing, we next performed single-nuclei RNA sequencing (snRNA-seq) in the striatum of 6-month-old mice (Fig. 6A). We targeted approximately 10,000 nuclei per sample (n=4/group, 16 samples total) and 188,303 nuclei passed the quality control criteria for further analysis (Supplementary Table 1). Overall, we did not observe any significant differences in total RNA count across different treatment groups (Supplementary Fig. 5A).

**Fig. 6:**
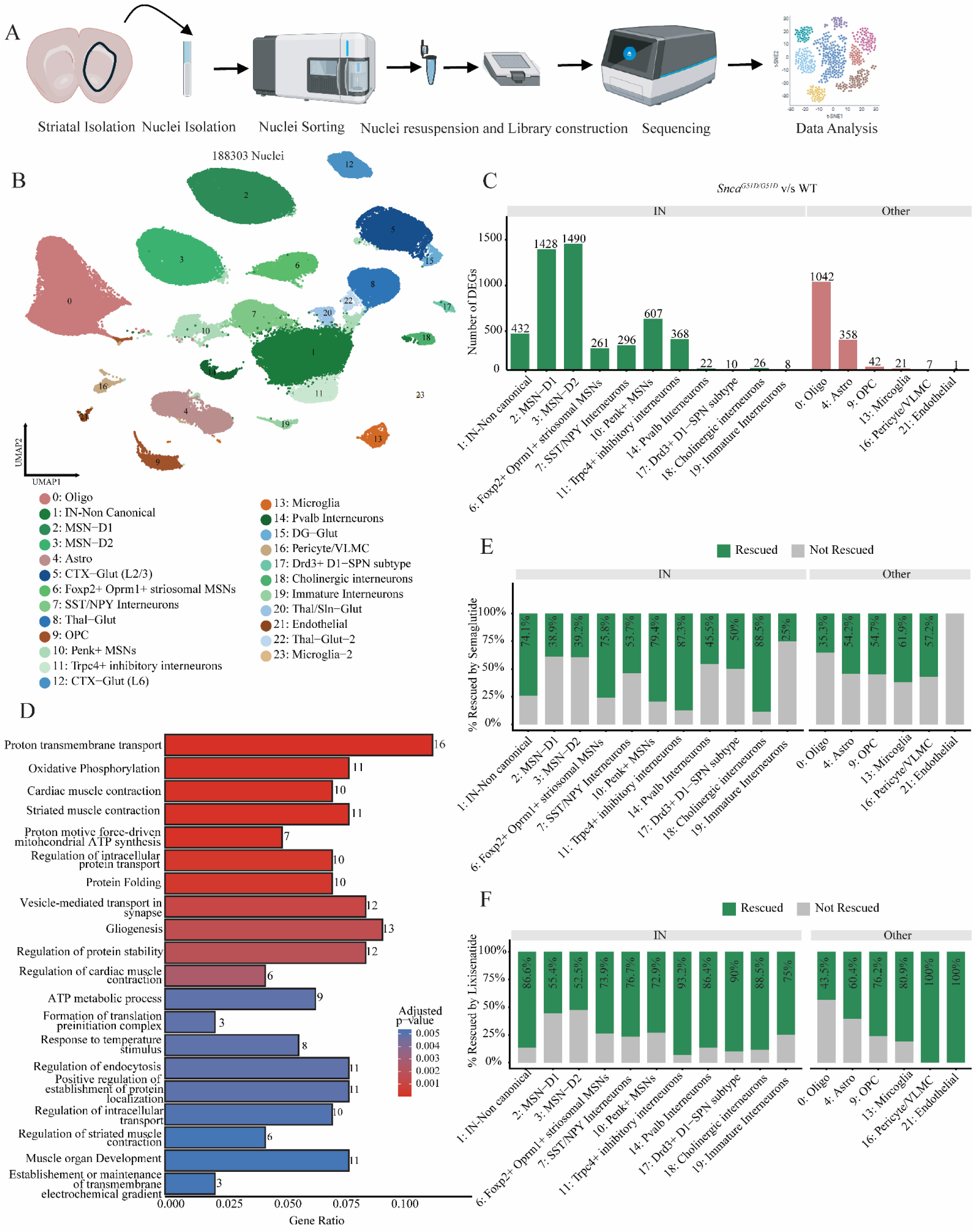
GLP-1 analogs treatment reverses transcriptomic dysregulation across different cell clusters in the striatum of *Snca^G51D/G51D^* mice at 6 months. (A) Schematic of the experimental design involving striatal nuclei isolation, sorting, library preparation, sequencing, and data analysis (B) UMAP of all striatal nuclei, which passed the quality control across different groups and annotated by cell type. (C) Bar graphs showing the number of differentially expressed genes (|log2FC|>0.15 and adjusted p-value <0.05) either upregulated or downregulated within each cluster (D) Gene Ontology analysis of shared DEGs from top 3 transcriptionally altered clusters in *Snca^G51D/G51D^* mice compared to WT mice. (E) Rescue analysis of differentially expressed genes across different cell clusters following Semaglutide treatment. (F) Rescue analysis of differentially expressed genes across different cell clusters following Lixisenatide treatment.

We identified distinct cell clusters (Fig. 6B) by comparing their top unique marker genes with the known marker genes of striatal cell types from prior investigations (Supplementary Fig. 5B). Cell-type composition was additionally examined at the sample and cluster level across groups to contextualize downstream differential expression and communication analyses (Supplementary Table 2). Detailed mapping identified some cell clusters derived from the cortex or thalamus, which were removed from further analysis, as in other studies (*27*). We next focused on the observed transcriptional changes across clusters to reflect on the overall functional alterations previously identified from bulk RNA sequencing data.

Differential expression analysis across cell types revealed that medium spiny neurons (MSN-D1 and MSN-D2) exhibited the most prominent transcriptional alterations within the neuronal populations, while astrocytes and oligodendrocytes showed the most differentially expressed genes amongst the identified non-neuronal clusters in *Snca^G51D/G51D^* mice (Fig. 6C). Next, we overlapped DEGs across the top 3 clusters, MSN-D1, MSN-D2 and oligodendrocytes with the highest transcriptional burden (Supplementary Table 1). GO analysis of these shared genes revealed mitochondrial processes, such as proton membrane transport and oxidative phosphorylation, as the most prominently affected pathways in *Snca^G51D/G51D^* mice, consistent with our previously observed bulk RNA sequencing findings (Fig. 6D).

These findings suggest that early disease-associated changes in the striatum are driven by mitochondrial dysfunction in neuronal and non-neuronal populations prior to the onset of overt neuroinflammation. Importantly, treatment with GLP-1 analogs led to a broad normalization of these transcriptional alterations across multiple cell types, further indicating coordinated effects across multiple cell clusters on rescuing mitochondrial dysfunction within the striatum (Fig. 6E-F).

### GLP-1 analogs normalize intercellular neuregulin (NRG) and neurexin (NRXN) signaling networks in the *Snca^G51D/G51D^* mouse

We next performed the CellChat analysis on our single-nucleus RNA sequencing data to infer ligand-receptor-mediated signaling networks across groups (*28, 29*). This analysis was motivated by our prior observations of coordinated mitochondrial transcriptional dysregulation in the striatum of *Snca^G51D/G51D^* mice at 6 months, suggesting that these cell-intrinsic perturbations may result from changes in circuit-level intercellular signaling networks.

We first assessed the global patterns of intercellular communication, which revealed widespread yet structured alterations in *Snca^G51D/G51D^* mice relative to WT, indicating early remodeling of signaling networks across neuronal and non-neuronal populations (Fig. 7A). A similar increase in the strength of intercellular signaling has also been reported in previous analyses of existing PD datasets by other authors (*30*). Notably, treatment with both GLP-1 analogs, lixisenatide and semaglutide, selectively normalized pathways altered in *Snca^G51D/G51D^* mice rather than globally dampening intercellular signaling.

**Fig. 7:**
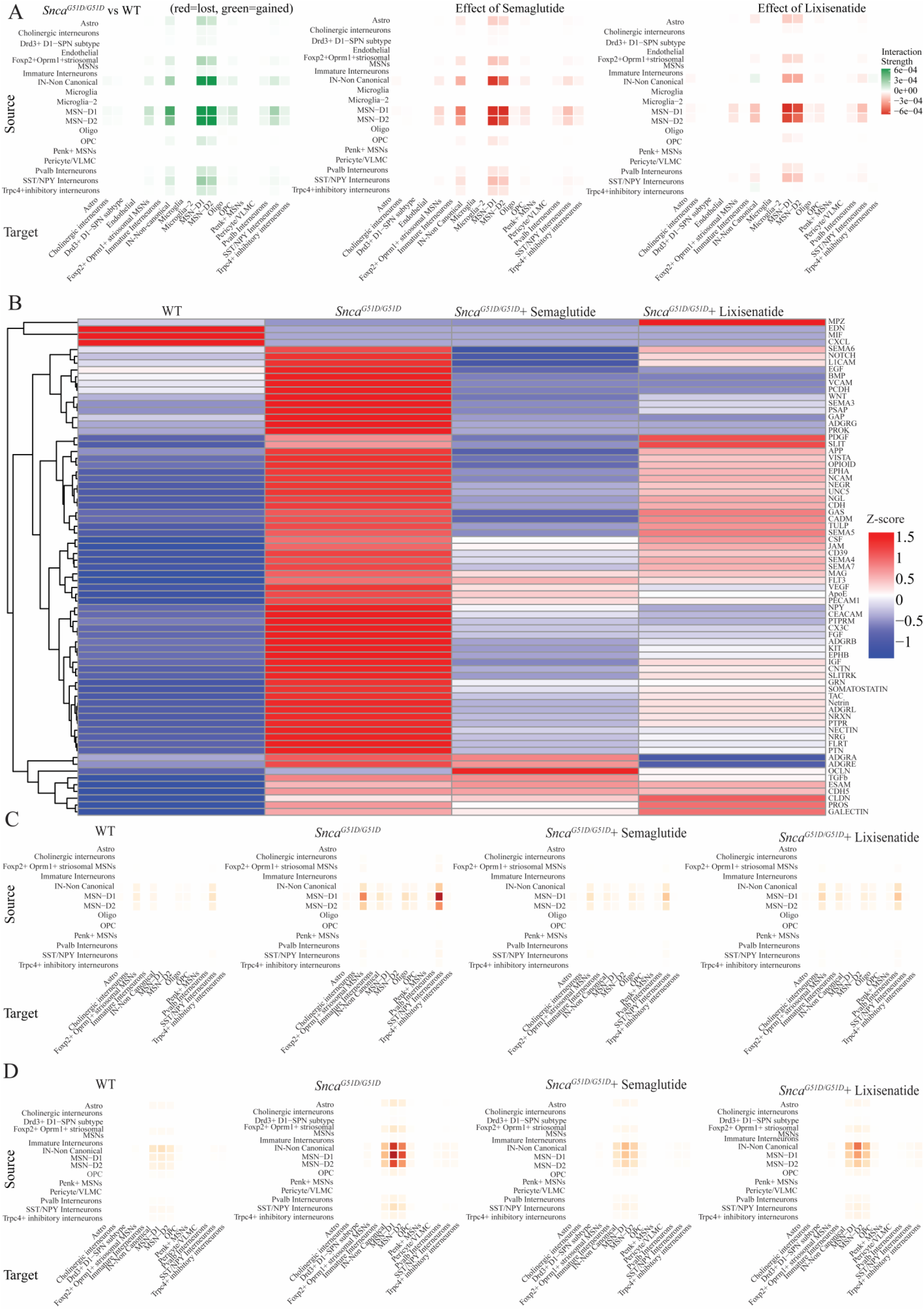
CellChat analysis reveals expansion of intercellular signaling networks in *Snca^G51D/G51D^* that are restored by GLP-1 analog treatment. (A) Heatmaps showing global differential interaction strength (WT vs *Snca^G51D/G51D^*) and the effects of GLP-1 analogs on cell–cell communication across annotated striatal cell types (B) Heatmap depicting major altered signaling pathways across WT, *Snca^G51D/G51D^*, and GLP-1-treated groups (C) Altered strength of NRG signaling across different treatment groups and different source and target striatal cell clusters (D) Altered strength of neurexin signaling across different treatment groups and different source and target striatal cell clusters

To identify distinct signaling pathways that might be responsible for the perturbation of mitochondrial genes, we next examined pathway-level alterations across groups. Multiple signaling programs were altered in *Snca^G51D/G51D^* mice compared to WT (Fig. 7B). However, the extent and consistency of these alterations varied across different cell types. Among the pathways analyzed, neuregulin (NRG) and neurexin (NRXN) signaling emerged as the most prominently altered across cell clusters (Supplementary text 1). Both pathways showed altered signaling activity in *Snca^G51D/G51D^* mice across diverse source-target cell pairs, which were robustly and uniformly normalized following GLP-1 analog treatment (Fig. 7C-D). Although both signaling pathways were centered on MSNs as primary sources, with MSNs and selected interneuron populations as the principal target clusters, *Snca^G51D/G51D^* mice exhibited additional source-target interactions, indicating a disease-linked expansion of intercellular communication networks. GLP-1 analog treatment normalized these interactions, thereby restoring a more constrained, WT-like signaling architecture. In contrast, other signaling pathways, although altered in disease, did not exhibit the same degree of consistent rescue across different clusters (Supplementary text 1). These findings indicate that NRG and NRXN represent key nodes of disease-associated signaling dysregulation that are particularly responsive to GLP-1 receptor activation.

## Discussion

In this study, we demonstrated the beneficial effects of the GLP-1 analogs, lixisenatide and semaglutide, in a genetically and clinically accurate mouse model of PD. We employed behavioral, molecular, transcriptomic, and intercellular communication analyses to show that GLP-1 analogs exert coordinated effects across multiple levels of PD pathology, including early prodromal phenotypes, motor deficits, inflammation, gliosis, mitochondrial alterations, and circuit-level signaling networks. These findings further provide a mechanistic framework for the effects of GLP-1 analogs in restoring early disease-associated changes in PD.

Prior studies evaluating GLP-1 receptor agonists have largely relied on toxin-induced late-stage PD models, such as MPTP and 6-OHDA, which cause acute dopaminergic injury rather than the stepwise disease progression observed in human PD (*31, 32*). One study reported the neuroprotective effect of GLP-1 analog NLY01 in α-synuclein preformed fibrils (α-syn PFF) and a human A53T α-synuclein (hA53T) transgenic mouse model of α-synucleinopathy-induced neurodegeneration. That study suggested that NLY01 treatment could rescue dopaminergic neuronal death, microglia-mediated activation of astrocytes and alpha-synuclein pathology in these PD models (*33*). However, overall detailed mechanistic insights and studies involving clinically accurate alpha-synuclein pathology driver model systems remain sparse, even though alpha-synuclein pathology is a defining feature in approximately 85% of PD patients (*34*). In the present study, we used *Snca^G51D/G51D^*mice (*12*), which replicate PD phenotypes at the behavioral, pathological, and transcriptomic levels. GLP-1 analogs were effective when administered both before and immediately after symptom onset in *Snca^G51D/G51D^* mice, resulting in a complete rescue of olfaction, constipation, and motor performance. This is particularly important considering PD’s long prodromal phase, where non-motor symptoms precede clinical diagnosis by several years (*2, 35*). Additionally, the precise dose selection ensured that mice receiving GLP-1 analogs did not show any constipation-related phenotypes, which are otherwise common side effects of clinically used GLP-1 analogs (*36*).

Previous studies have linked mitochondrial changes to PD pathogenesis (*37*), including complex I deficiency in patient brains (*38*), and established links between PINK1, PARKIN, and DJ-1 to mitochondrial quality control (*39, 40*) in PD. Moreover, a recent study also showed that transplantation of encapsulated mitochondria alleviates pathology in the MPTP PD model (*41*). Here, we identified mitochondrial dysregulation as the earliest driver of PD pathology in *Snca^G51D/G51D^* mice, occurring even before symptom onset. Our findings further show that mitochondrial alterations are responsive to GLP-1 analog treatment, ultimately resulting in a strong normalization of differentially expressed genes (∼92%) in our PD model. Although GLP-1 analogs have been shown to exert protective effects on mitochondrial function and oxidative stress–dependent pathways in diabetes and cardiac tissues (*42–44*), analogous effects have not previously been reported in PD or other neurodegenerative disorders. Our results reveal mitochondrial alterations as an important, early, and central driver of PD pathology that are completely normalized with the timely initiation of GLP-1 therapy.

When we specifically examined the inflammation-related DEGs, almost complete normalization of the inflammation-linked disease signatures was observed following GLP-1 treatment. However, we observed that these changes arose before the onset of astrogliosis and microgliosis reported in our mice at later ages (*12*). Attenuation of these changes in the GLP-1-treated groups prevented the onset of this apparent inflammatory response, which is another critical contributor to PD progression and alpha-synuclein toxicity (*45, 46*).

In addition to these molecular perturbations, we observed a marked reduction in detergent-insoluble α-synuclein across multiple brain regions following GLP-1 analog treatment, including the midbrain, olfactory bulb, striatum, and piriform cortex. As the accumulation of insoluble α-synuclein aggregates is a key driver of neurotoxicity in PD (*47*), these findings further highlight that GLP-1 analogs may exert disease-modifying effects by reducing aggregation-prone α-synuclein species, potentially through restoration of mitochondrial pathways and decreasing inflammation upstream of these changes.

Importantly, the absence of striatal gliosis at 6 months, despite transcriptional alterations, suggests that molecular dysfunction precedes overt neuroinflammation and neurodegeneration and could be targeted to reverse PD pathogenesis. Interestingly, mitochondrial dysfunction was the transcriptionally shared signature among the most transcriptionally altered cell clusters, particularly MSNs and oligodendrocytes, in single-nuclei transcriptomic analysis. These findings are consistent with the high bioenergetic demands and selective vulnerability of MSNs in PD (*48*). GLP-1 analog treatment significantly rescued the transcriptomic program across multiple clusters, attenuating gene expression changes associated with mitochondrial dysfunction. Overall, this suggests that GLP-1 analogs exert system-wide effects that improve mitochondrial function rather than targeting a single cell type.

These transcriptional changes disrupted intercellular communication networks, which may be an important feature of early PD. Among the signaling pathways altered in the PD model, NRG and NRXN signaling emerged as the most prominently altered and consistently rescued following GLP-1 analog treatment. Both pathways play important roles in synaptic organization and the maintenance of circuit integrity (*49, 50*). Emerging evidence links PD to disruption of these signaling pathways and to mutations in the genes encoding their component proteins (*48, 51, 52*). However, it is possible that these correlations are not merely a reflection of altered intercellular communication and synaptic dysregulation, but may also contribute directly to bioenergetic failure and mitochondrial dysfunctions in vulnerable PD circuits. In particular, neuregulin signaling has been mechanistically linked to mitochondrial homeostasis, directly regulating cellular respiration, mitochondrial content, and oxidative stress (*53–55*). Data from *Drosophila* studies also implicate neurexins in cellular metabolism. Loss of Neurexin 1 has been shown to affect NAD⁺ levels, resulting in mitochondrial structural abnormalities, including disrupted cristae architecture, consistent with impaired bioenergetic capacity (*56*). Moreover, NRG and NRXN signaling also modulate neuroinflammation by attenuating NLRP3 inflammasome, NF-κB pathways, and pro-inflammatory cytokines (e.g., TNF-α, IL-1β) in models of cerebral ischemia-reperfusion injury and inflammatory pain (*57, 58*). Additionally, alpha-synuclein has been shown to bind to various neurexins, including neurexin 1β, neurexin 3β, neurexin 1α, and neurexin 2β, which could have deleterious implications for PD (*59*). Taken together, these observations indicate that a coordinated dysregulation of NRG and NRXN signaling may converge on mitochondrial dysfunction as a key event in early PD, with potential consequences for alpha-synuclein pathology. Therefore, normalization of these pathways by GLP-1 analogs, as revealed by CellChat analysis, likely reflects a broader restoration of circuit integrity in which restored strength of intercellular signaling supports mitochondrial function, reduces neuroinflammation, and re-establishes network-level homeostasis. Such restoration may prevent inflammatory amplification of α-synuclein pathology, creating a multi-pronged therapeutic mechanism. These region-specific effects, particularly in the striatum, highlight the potential of GLP-1 analogs to preserve circuit integrity across PD-vulnerable networks.

These findings provide important context for the mixed results seen in GLP-1 clinical trials for PD. While initial trials for drugs like exenatide demonstrated some benefits (*60*), subsequent trials have yielded variable outcomes (*9*), and recent data for drugs like lixisenatide indicate a modest decrease in disease progression (*5*). Furthermore, epidemiological studies have indicated reduced PD risk among diabetic patients receiving GLP-1 analogs (*61*). Therefore, the overall variability may be attributed to the exact disease stage at treatment initiation and to the underlying genetic variability among patients. Our data support the hypothesis that GLP-1 analogs might be effective when initiated during relatively early stages of disease. Although GLP-1 analogs appear efficacious even after symptom onset in *Snca^G51D/G51D^* mice, the disease is not yet significantly advanced; therefore, considering a trial in those with early PD might be beneficial. Most importantly, because PD cases are highly heterogeneous, GLP-1 analogs might be effective in a particular subtype of PD patients. As *Snca^G51D/G51D^* mice recapitulate the clinical progression of PD driven primarily by synuclein pathology, the findings of this study could be extrapolated to approximately 85% of PD patients in whom alpha-synuclein drives pathology (*11*). Hence, reanalysis of existing clinical trial data and further patient stratification, potentially using alpha-synuclein skin biopsies or seed amplification assays, could pave the way for future clinical trials with GLP-1 analogs.

Collectively, our study provides detailed mechanistic insights into the beneficial effects of GLP-1 analogs in PD and identifies the coordinated restoration of mitochondrial, inflammatory, and intercellular signaling pathways as a central mechanism underlying their therapeutic benefits. We envision that our understanding of GLP-1-based therapies will be important for targeting early disease processes in PD and beyond.

## Materials and Methods

### Animals

Baylor College of Medicine Institutional Animal Care & Use Committee (Protocol number: AN-1013) approved all the experiments. Mice used in this study were housed in a level three American Association for Laboratory Animal Science-certified animal facility maintained on a 14/10h light/dark cycle and provided with the standard chow diet and water *ad libitum*.

### Subcutaneous Dosing of GLP-1 analogs

WT and *Snca^G51D/G51D^* were divided in two cohorts, each comprising six groups: WT, *Snca^G51D/G51D^, Snca^G51D/G51D^* +Lixisenatide (10 nmol/kg), *Snca^G51D/G51D^* +Semaglutide (30 nmol/kg), WT +Lixisenatide (10 nmol/kg), and WT + Semaglutide (30 nmol/kg). The cohorts were dosed subcutaneously daily for 2 months, from four to six months, and from seven to nine months, respectively.

### Behavioral Assays

Mice for behavioral assays were acclimatized for 30 min before the start of experiments. Behavioral tests were conducted in male and female mice at 6 and 9 months of age.

### Parallel rod floor test

Motor coordination of mice was tested using the parallel rod floor test. Briefly, mice were placed in the center of an open field fitted with an underlying wire grid and allowed to explore for 10 min. The number of foot slips was quantified automatically using the ANY-maze behavior and video-tracking system (Stoelting). To account for locomotor activity, foot slips were normalized to the distance traveled by each mouse during the trial.

### Rotarod

Mice were trained on the accelerating rotarod paradigm, which consisted of a cylinder accelerating from 5 to 40 rpm over 5 min (Type 7650, Ugo Basile). Mice underwent 4 trials per day, with a 10-min cutoff and a minimum intertrial interval of 30 min. A trial was completed when either the mouse fell off the rotating rod or rode the cylinder for two consecutive revolutions without falling. Data collected on day 3 was compared across groups.

### Buried food pellet Test

The buried food pellet test was performed using a rectangular box with an inch-thick layer of bedding. Each mouse underwent a 15-min habituation trial in the box each day for 3 days. At the end of the third day, mice were fasted for 24 h, followed by a probe trial the next day. During the probe trial, a food pellet was placed approximately 3 mm below the bedding surface and the time the mouse took to find it was recorded.

### Whole gut transit time

Mice were given carmine red dye (150 µl of 6% solution) via oral gavage and then housed individually in a sterile cage. Mice were checked every 10 min, and the time to the first appearance of dye-stained fecal pellets was recorded as the measure of whole gut transit time.

### Water content in the stool

Mice were housed individually in a sterile, clean cage for 60 min. Freshly expelled fecal pellets by each mouse were collected immediately in a pre-weighed 1.5 ml microcentrifuge tube. The tubes were weighed again after collection to note the wet weight. Thereafter, the pellets were dried overnight at 55 °C, and the dry weight was noted. Finally, the % water content of the stool was calculated by subtracting the dry weight from the total weight of the fecal pellets, then dividing by the total weight.

### Triton X-100 Soluble/Insoluble Fractionation and immunoblotting

Triton X-100 Soluble/Insoluble Fractionation was carried out as described previously (*62*). Briefly, 9-month-old wild-type and *Snca^G51D/G51D^* mouse brain tissues were lysed in Phosphate-buffered saline (PBS) with 1% triton X-100. For the preparation of the detergent-soluble fraction, the tissue lysates were mixed with an equal volume of 2× concentrated RIPA buffer without SDS, and the mixture was centrifuged at 4 °C for 30 min at 100,000g. The supernatants were collected and kept separately, while the pellet was dissolved and incubated with 1× RIPA containing 1% sarkosyl (without SDS) for 60 min on an orbital shaker at room temperature. The supernatant was then collected by centrifugation at 200,000g for 30 min at 4 °C. Each separated supernatant fraction was mixed with the sample buffer and heated to 95 °C for 5 min.

Equal amounts of protein were loaded onto 17-well NuPAGE 4 to 12% Bis-Tris gels (Thermo Fisher Scientific). Gels were run in 1× NuPAGE™ MOPS SDS running buffer and transferred onto nitrocellulose membranes in Tris-Glycine transfer buffer at 110V for 2 h. Thereafter, the membranes were blocked with 5% bovine serum albumin in Tris-buffered saline with 0.1% Tween 20 (TBST) for 1 h, and incubated overnight with mouse anti-vinculin (1:10,000) and mouse anti-α-Syn (1:3,000). Thereafter, the membranes were washed with TBST and probed for 1 h with HRP-conjugated anti-mouse secondary antibodies. Finally, the membranes were washed, and chemiluminescence was detected with ECL (GE Healthcare, RPN2236) and imaged with an Amersham imager 680 (GE Healthcare) (*63*).

### Immunofluorescence (IF)

Tissue preparation and immunofluorescence were performed as previously described (*64*). Briefly, mice were transcranially perfused with PBS followed by 4% paraformaldehyde (PFA). Thereafter, the brains were removed and fixed for 48h in 4% PFA at 4 °C. The brains were then dehydrated for 24 h in 15% w/v sucrose solution in PBS, followed by a 2-day incubation in 30% w/v sucrose solution in PBS at 4 °C. The brains were then frozen in cryomolds in Optimal Cut Temperature (OCT) compound. Subsequently, 30 µm floating sections (olfactory bulb, striatum, piriform cortex, and substantia nigra) were cut using a cryostat (Leica CM 3050S) and stored in 1× PBS containing 0.01% NaN3. For immunofluorescence, floating sections were washed with 1× PBS and blocked with 5% donkey serum in 0.3% Triton X-100 at room temperature for an hour. Afterward, sections were incubated overnight with primary antibodies, goat anti-GFAP (1:1000) and rabbit anti-Iba1 (1:1000). The next day, sections were washed and incubated with secondary antibodies (Alexa Fluor 488 or 555) at a 1:1000 dilution for 3 h. Finally, the sections were washed, mounted with VECTASHIELD® PLUS Antifade Mounting Medium with DAPI, and imaged on a confocal microscope (Leica STED TCS SP8X) using LAS X software (Leica) at 63× magnification.

### Estimation of peripheral metabolic parameters

Intracardiac blood was drawn at the end of the study. Plasma was isolated as previously described (*65*). Peripheral metabolic parameters were measured as described in the kit instructions for glucose, cholesterol species, triglycerides, and insulin.

### Bulk RNA sequencing

Striatal tissues were dissected from the brains of mice in different groups at 6 months of age and flash-frozen in liquid nitrogen. Tissues were homogenized using the TissueLyser II (Qiagen, #85300), and RNA was extracted using the RNeasy Mini Kit (Cat#75106, Qiagen) in accordance with the manufacturer’s protocol. Thereafter, RNA levels were quantified using a NanoDrop and samples were sent to GeneWiz for library preparation and sequencing on the Illumina HiSeq platform. Approximately 80-100 million 150 bp paired-end reads were generated for each sample. Raw reads were trimmed and mapped using default parameters using Trimmomatic v0.39.71. Trimmed reads were aligned to GRCm39/mm39, GENCODE vM36 assembly using all default parameters. The count matrix of all samples was imported into R for differential expression analysis using DESeq2. Genes with fewer than 10 total counts summed across all samples were removed prior to analysis.

#### Disease signature

To characterize the baseline PD transcriptional phenotype with maximal statistical clarity, WT and *Snca^G51D/G51D^* samples were extracted from the full dataset comprising four experimental groups (WT, *Snca^G51D/G51D^, Snca^G51D/G51D^* + Semaglutide, *Snca^G51D/G51D^* + Lixisenatide, n=4 biological replicates each) and analyzed as an independent two-group DESeq2 contrast (∼ group, reference: WT). This subset design isolates the disease contrast from the variance contributed by drug-treated samples. Size factor normalization and dispersion estimation were performed using DESeq2 defaults.

#### Drug treatment comparisons

All remaining comparisons: drug rescue (*Snca^G51D/G51D^* + Semaglutide vs. WT; *Snca^G51D/G51D^* + Lixisenatide vs. WT) and drug mechanism (*Snca^G51D/G51D^* + Semaglutide vs. *Snca^G51D/G51D^*; *Snca^G51D/G51D^* + Lixisenatide vs. *Snca^G51D/G51D^*) were performed using a single combined DESeq2 object constructed from all four groups simultaneously. This combined approach provides more stable dispersion estimates by pooling variance information across all samples and avoids inflated significance that can arise from pairwise-only analyses.

Pairwise contrasts were extracted using the results function. Genes with Benjamini-Hochberg adjusted p-value < 0.05 and |log2 fold change| > 0.15 were considered significantly differentially expressed. For GSEA, all detected genes were retained and ranked by the DESeq2 Wald statistic without pre-filtering. Sample quality was assessed by principal component analysis (PCA) of variance-stabilizing transformed (VST) counts.

#### Drug Rescue Quantification

To assess transcriptional normalization toward wild-type, disease-dysregulated genes (*Snca^G51D/G51D^* vs. WT, padj < 0.05, |log2FC| > 0.15) were cross-referenced against drug rescue results from the combined analysis (*Snca^G51D/G51D^* + Semaglutide vs. WT and *Snca^G51D/G51D^* + Lixisenatide vs. WT). Genes absent from the drug comparison results as well as the ones which were still significant but with attenuated fold change in the same direction as disease, |log2FC| > 0.15) were classified as rescued.

#### Gene Set Enrichment Analysis (GSEA)

GSEA was performed using the clusterProfiler package (v4.14.6) in R. For each pairwise comparison (disease signature: *Snca^G51D/G51D^* vs. WT; drug effects: *Snca^G51D/G51D^*+ Semaglutide vs. *Snca^G51D/G51D^* and *Snca^G51D/G51D^*+ Lixisenatide vs. *Snca^G51D/G51D^*; drug rescue: *Snca^G51D/G51D^*+ Semaglutide vs. WT and *Snca^G51D/G51D^* + Lixisenatide vs. WT), all detected genes were included in ranked gene lists without pre-filtering for significance. Genes were ranked by the DESeq2 Wald statistic, which integrates both effect size and statistical confidence into a single metric. Genes with significant differential expression were defined as those with an adjusted p-value < 0.05 (Benjamini-Hochberg correction) and |log2 foldchange| > 0.15. This threshold was applied when reporting DEG lists, but was not used to filter the input to GSEA.

GSEA was performed against four pathway databases: Gene Ontology (GO) Biological Process (BP), Cellular Component (CC), and Molecular Function (MF), and the Kyoto Encyclopedia of Genes and Genomes (KEGG). GO enrichment was performed using gseGO with the mouse genome annotation database (org.Mm.eg.db, gene symbols as identifiers). For KEGG analysis, gene symbols were converted to Entrez IDs via bitr prior to running gseKEGG (organism code “mmu”). All analyses used a minimum gene set size of 10, a maximum of 500, and a Benjamini-Hochberg adjusted p-value cutoff of 0.05.

### Nuclei isolation

Nuclei isolation was performed as described previously and was adapted from the 10x Genomics protocol for adult brain tissue (*66*). Briefly, the striatum was lysed in an ice-cold lysis buffer (10 mM NaCl, 3 mM MgCl2, 10 mM Tris-HCl, pH 7.4, and 0.1% IGEPAL CA-630 from Sigma-Aldrich) and incubated on ice for 10 min. The lysates were then filtered through a MACS SmartStrainer (30-μm) to remove debris and were finally centrifuged at 500g for 300 s at 4°C. Subsequently, the supernatant was discarded, and the pellet was washed with a wash buffer (1% BSA in 1X Dulbecco’s phosphate-buffered saline). Following three washes, the concentration of nuclei in the final suspension in 3 mL of wash buffer was measured using a Thermo Fisher Countess II with Trypan blue. The concentrations were kept below 1 × 10^7 to prevent clogging of the SONY MA900 flow cytometer during nuclei sorting.

### Nuclei Sorting

DRAQ5 fluorescent probe (1:1000 dilution) was used to stain nuclei suspensions approximately 5 min before the beginning of nuclei sorting, as described earlier (*66*). The forward scatter (FSC) threshold was set at 1.50% with an FSC gain of 11, and the back scatter (BSC) gain was adjusted to 30.5%. The gating was determined by initially passing the prepared samples through the (BSC-A vs FSC-A) gate to filter out debris using control nuclei without DRAQ5 staining, then through two single-nucleus gates (FSC-H vs FSC-A and BSC-H vs BSC-W). Prior to sorting, the collection tubes were coated with 1% bovine serum albumin and filled with single-cell sorting buffer (50 μL of 1X DPBS with 1% BSA). Thereafter, the nuclei suspension was centrifuged at 1500 × g for 10 min at 4°C. Finally, the nuclei were resuspended to an approximate concentration of 1000 nuclei/μL. Finally, nuclei were prepared and loaded on the 10X GEM-X Chip according to the 10x Genomics user guide (CG000733) (*67*).

### Library construction and sequencing

The cDNA libraries were constructed in accordance with the manufacturer’s instructions for the 10x Genomics Chromium GEM-X 5’ Gene Expression v3 chemistry. Droplets containing nuclei, reverse transcription reagents, and barcoded gel beads were generated by Chromium X Controller. The first-strand cDNA was then amplified and checked on a Tape Station using HS D5000 Screen Tape. Subsequently, the cDNA libraries were fragmented, size-selected using SPRI beads, and ligated with sequencing adapters and sample indices. The libraries were sequenced at Azenta Life Science using Illumina NovaSeq X Plus with 3 25B lanes to achieve a target read depth of 50000 reads per nucleus.

### Data pre-processing

Raw sequencing reads were aligned to the mouse reference genome (GRCm39, 2024-A release) and quantified using Cell Ranger Count v9.0.1, with intronic reads included to maximize nuclear transcript capture (includeIntrons:true). BAM files were generated for quality verification. All other parameters were set to default. Filtered count matrices were imported into R using Seurat (v5). Prior to filtering, the dataset comprised 221,633 nuclei with an average of 52,608 reads per nucleus, 60.7% sequencing saturation, and 74.1% of reads mapping within cells, indicating adequate sequencing depth and library quality. Nuclei were subjected to initial quality filtering, retaining cells with 200–6,000 detected genes, 500–50,000 UMI counts, and ≤5% mitochondrial gene content. Doublets were identified and removed using scDblFinder, applying empirical doublet rates derived from 10x Genomics GEM-X 5’ v3 chemistry specifications. Ambient RNA contamination was estimated and corrected using decontX, which models contamination at the per-cell level; decontX-corrected count matrices were used for all downstream analyses. In total, 188,303 nuclei passed all quality control and were carried forward into downstream analyses. Following ambient RNA removal, corrected counts were log-normalized and the top 2,000 highly variable features were identified. Principal component analysis (PCA) was performed on scaled data, and the number of informative PCs was determined by elbow plot inspection (20 PCs retained). UMAP visualization and unsupervised clustering were performed directly on the PCA embedding. Sample mixing was quantitatively assessed by computing Shannon entropy of sample composition per cluster. The majority of clusters exhibited normalized entropy >0.95 (mean across all clusters: 0.93), indicating homogeneous distribution of nuclei from all 16 samples and no substantial technical batch effect requiring correction. Unsupervised clustering was performed at resolution 0.2 using the Louvain algorithm, yielding 24 distinct clusters (clusters 0–23).

### Cell type annotation

Cluster identities were assigned through a multi-step process: (1) supervised annotation using canonical striatal marker genes for major cell classes (MSNs, interneurons, astrocytes, oligodendrocytes, OPCs, microglia, and vascular cells); (2) automated annotation of a representative cell subset using MapMyCells against the Allen Brain Cell Atlas (2024); and (3) manual curation reconciling both approaches with literature-based marker validation. Nuclei mapping to non-striatal populations (cortical, thalamic, hippocampal) based on Allen Brain Atlas annotations were identified and excluded prior to specific downstream analysis.

### Identification of differentially expressed genes (DEGs) in snRNA-seq

Differential expression analysis was performed using MAST, with logfc.threshold set to 0 to capture subtle transcriptional changes. The disease signature was defined as genes differentially expressed between *Snca^G51D/G51D^* and WT (|log₂FC| > 0.15 and adjusted p-value < 0.05). Drug rescue was assessed by comparing treatment groups (*Snca^G51D/G51D^* + Semaglutide or *Snca^G51D/G51D^*+ Lixisenatide) against both WT and *Snca^G51D/G51D^*. Genes were classified as rescued or not rescued based on their fold-change trajectories across conditions, using criteria similar to those used for bulk RNA sequencing.

#### Gene ontology enrichment analysis

Gene ontology (GO) enrichment analysis was performed using clusterProfiler with the org.Mm.eg.db annotation database. Disease signature genes were mapped to Entrez IDs and tested for enrichment in Biological Process, Molecular Function, and Cellular Component ontologies. To avoid inflated enrichment statistics, the background gene set was defined as the full expressed transcriptome (∼23,592 Entrez-mapped genes) detected across all cells in the decontX assay.

### CellChat for pathway identification

Cell-cell communication analysis was performed using CellChat with the mouse CellChatDB, combining the Secreted Signaling and Cell-Cell Contact interaction categories. Analysis was restricted to interneuron and striatal cell populations. CellChat objects were constructed independently for each experimental condition using decontX-corrected count matrices. Pathway-level information flow was defined as the total communication probability summed across all cell type pairs for a given signaling pathway computed for each condition and compiled into a pathway * condition matrix. Pathways present in at least one condition were retained for comparison. To identify PD-associated communication changes, log₂ fold change in information flow between *Snca^G51D/G51D^* and WT was calculated; pathways exceeding |log₂FC| > 0.15 were considered altered in disease, consistent with the significance threshold applied in bulk RNA sequencing analyses. A pathway was classified as rescued by drug treatment if the drug effect (*Snca^G51D/G51D^* + Semaglutide or *Snca^G51D/G51D^* + Lixisenatide vs. *Snca^G51D/G51D^*) exceeded the same |log₂FC| > 0.15 threshold and was directionally opposite to the disease effect, indicating reversal of PD-associated gains or losses in intercellular communication. Relative pathway activity across conditions was visualized as a z-scored heatmap ordered by the magnitude of PD-associated change (WT-*Snca^G51D/G51D^* difference).

### Statistical Analyses

For all experiments involving more than two groups, data were analyzed using one-way ANOVA followed by Tukey’s post hoc test. All analyses were conducted using GraphPad Prism 8 software unless otherwise specified. Data for RNA sequencing was analyzed using R

## Acknowledgments

We thank the Baylor College of Medicine Genetically Engineered Mouse Core and the following cores (Microscopy Core, Neuropathology Core, and Animal Behavior Core) supported by the Jan and Dan Duncan Neurological Research Institute (NRI) and The Baylor College of Medicine Intellectual and Developmental Disabilities Research Center (BCM-IDDRC) from the Eunice Kennedy Shriver National Institute of Child Health and Development (NICHD-U54HD083092). We also thank Zhijian Yu, Hu Chen and the NRI Bioinformatics core for aligning and processing data for Bulk RNA Sequencing. This work was supported by the Howard Hughes Medical Institute, the Huffington Foundation, and the Freedom Together Foundation.

## Author contributions

B.V. and H.Y.Z. conceived the study and designed the experiments. B.V., Y.L. and Y.K. performed experiments, analyzed and interpreted data. C.O. performed experiments, J.P.R. sorted nuclei using flow cytometry. B.V. and Y.L. analyzed data for bulk RNA Sequencing and single-nuclei RNA sequencing and wrote the manuscript. H.Y.Z edited the manuscript.

## Data, code, and materials availability

All data associated with this study are present in the paper or the Supplementary Materials. All the reagents and chemicals used in the manuscript have been listed in Supplementary Table 3. All other materials used or generated in this study are commercially available or will be supplied upon reasonable request.

## Supplementary Figures

**Supplementary Fig 1:**
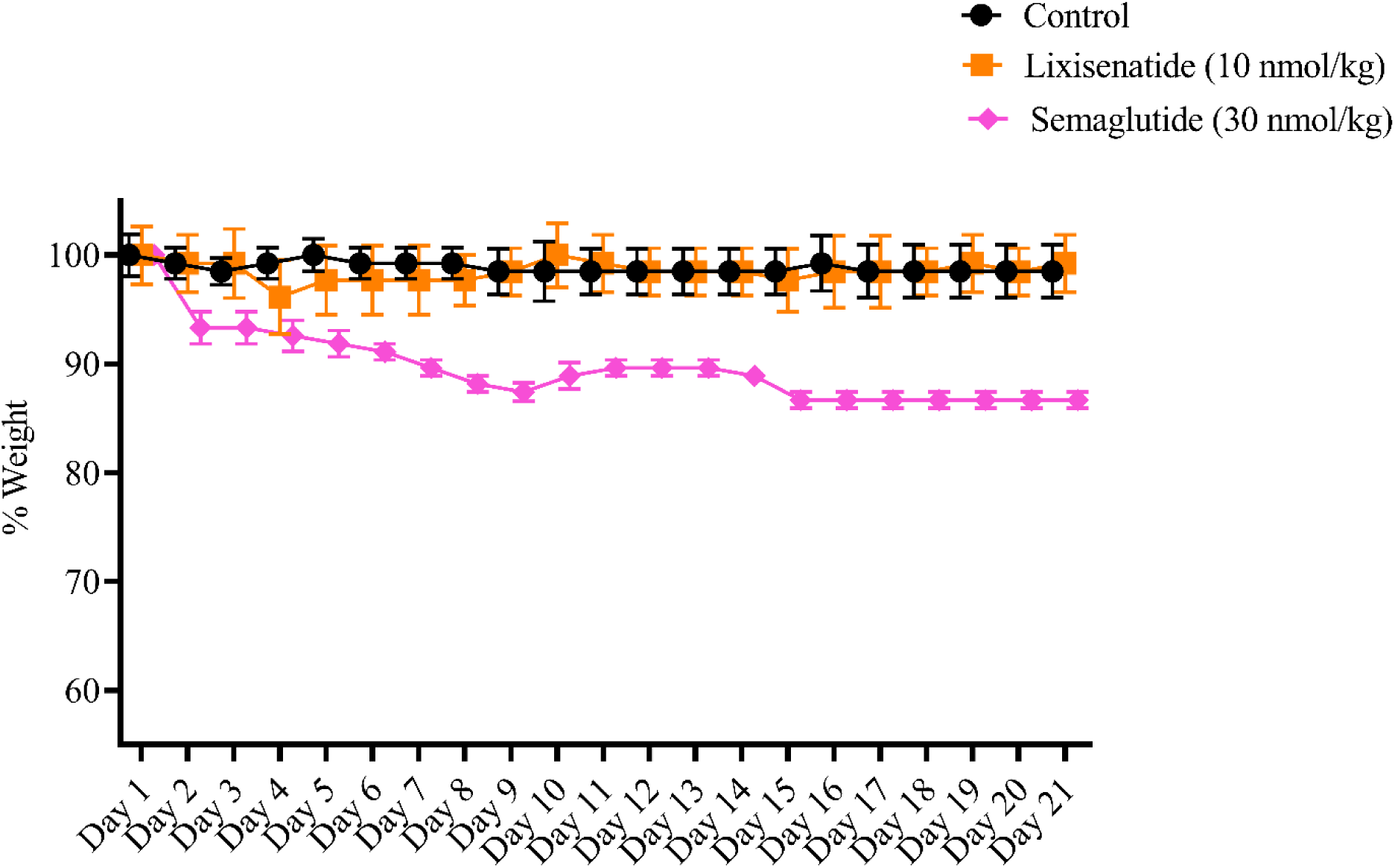
Dose Standardization for GLP-1 analogs. Lixisenatide (10 nmol/kg) and Semaglutide (30 nmol/kg) administration for 3 weeks in 6-month-old WT mice did not cause more than 10% weight loss and were therefore selected for further experiments.

**Supplementary Fig 2:**
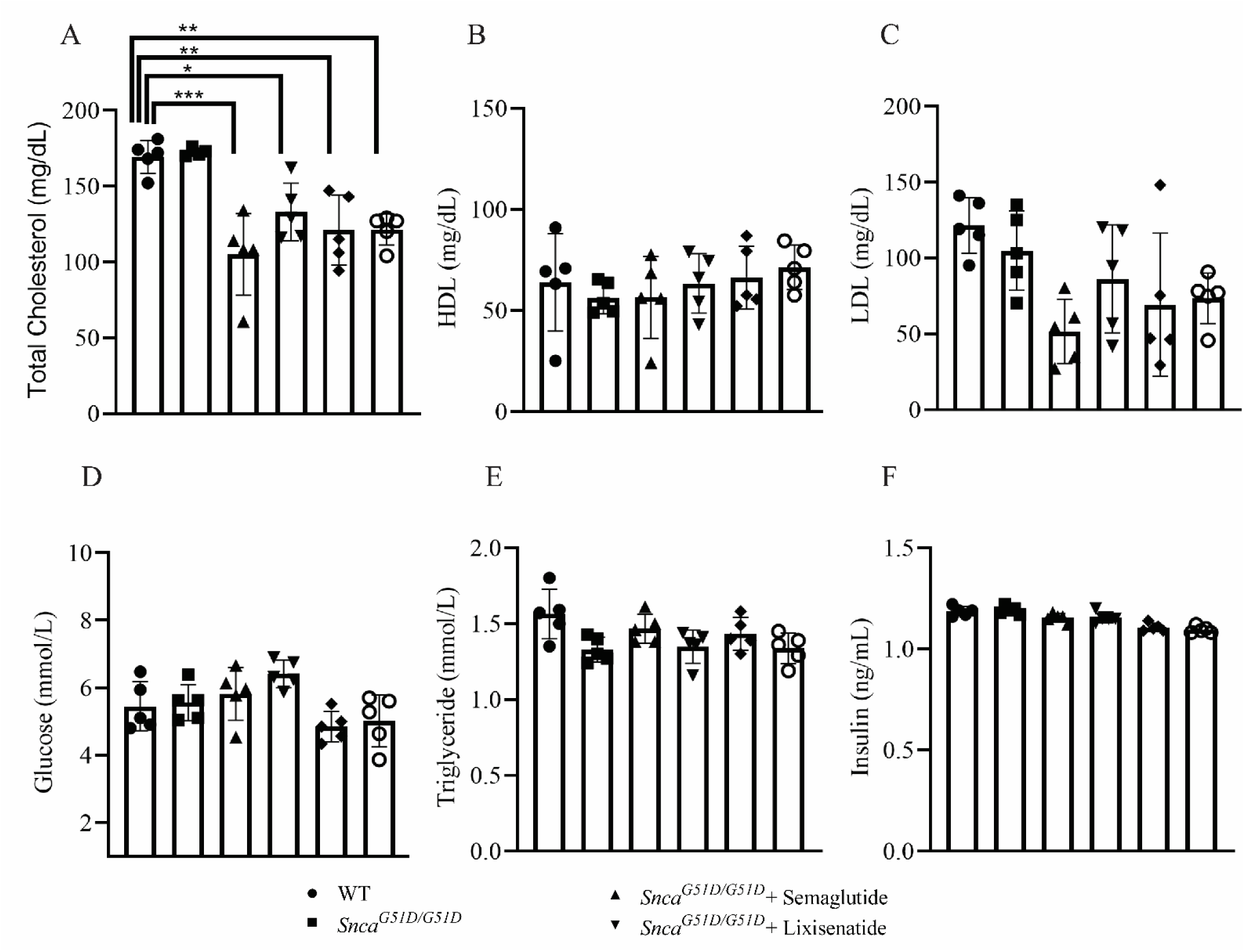
Effect of GLP-1 analogs on plasma metabolic parameters at 9 months (A) Total Cholesterol levels (B) HDL levels (C) LDL levels (D) Glucose levels (E) Triglycerides levels (F) Insulin levels. Individual data points represent biological replicates. Results are expressed as mean ± SD. ***p<0.001, **p<0.01, *0<0.05.

**Supplementary Fig 3:**
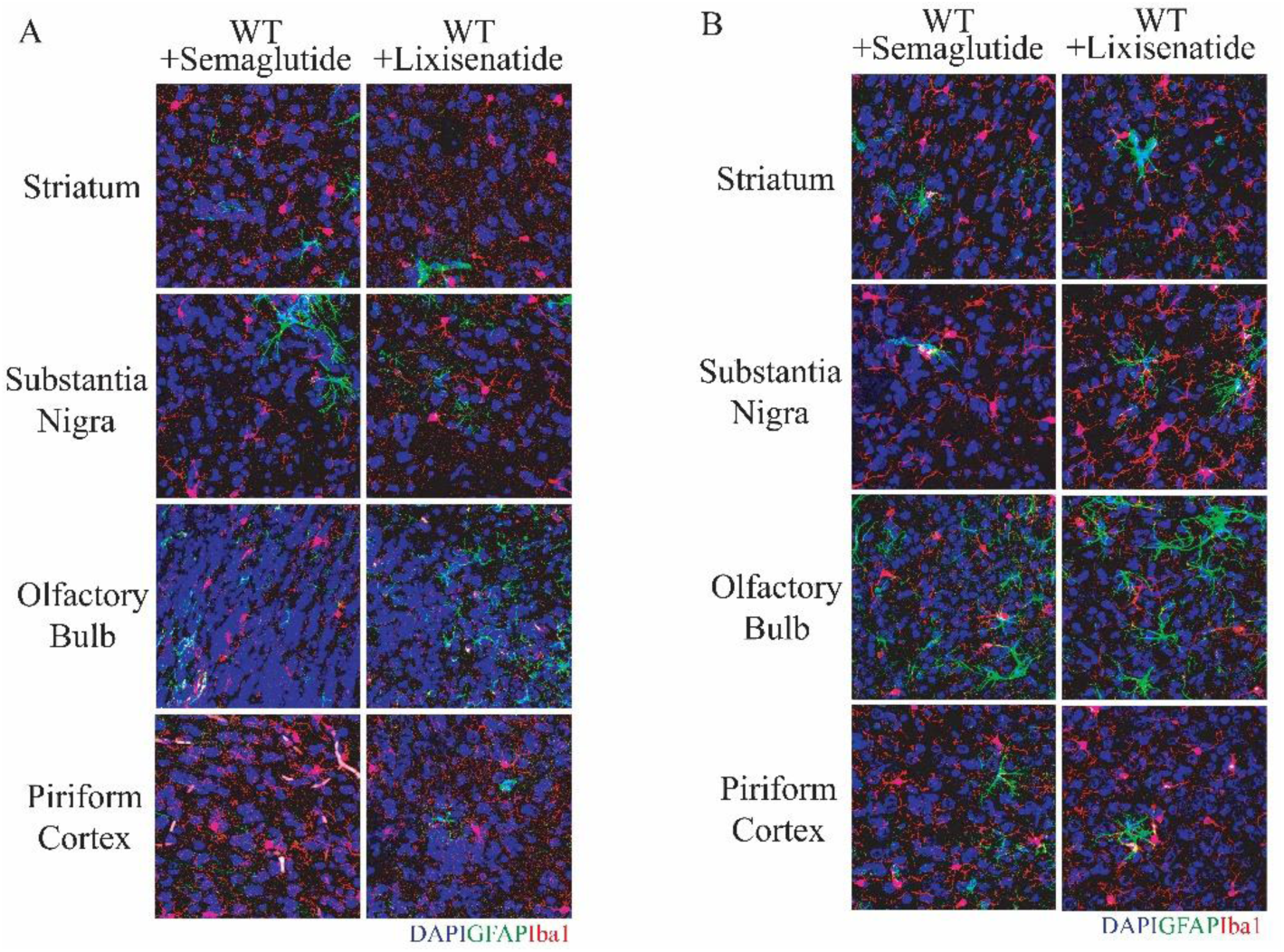
GLP-1 analogs did not exhibit gliosis in WT animals across different brain regions. Representative images of astrocyte (GFAP) and microglia (Iba1) staining in the striatum, substantia nigra, olfactory bulb and piriform cortex of WT animals treated with GLP-1 analogs at (A) 6 months and (B) 9 months.

**Supplementary Fig. 4:**
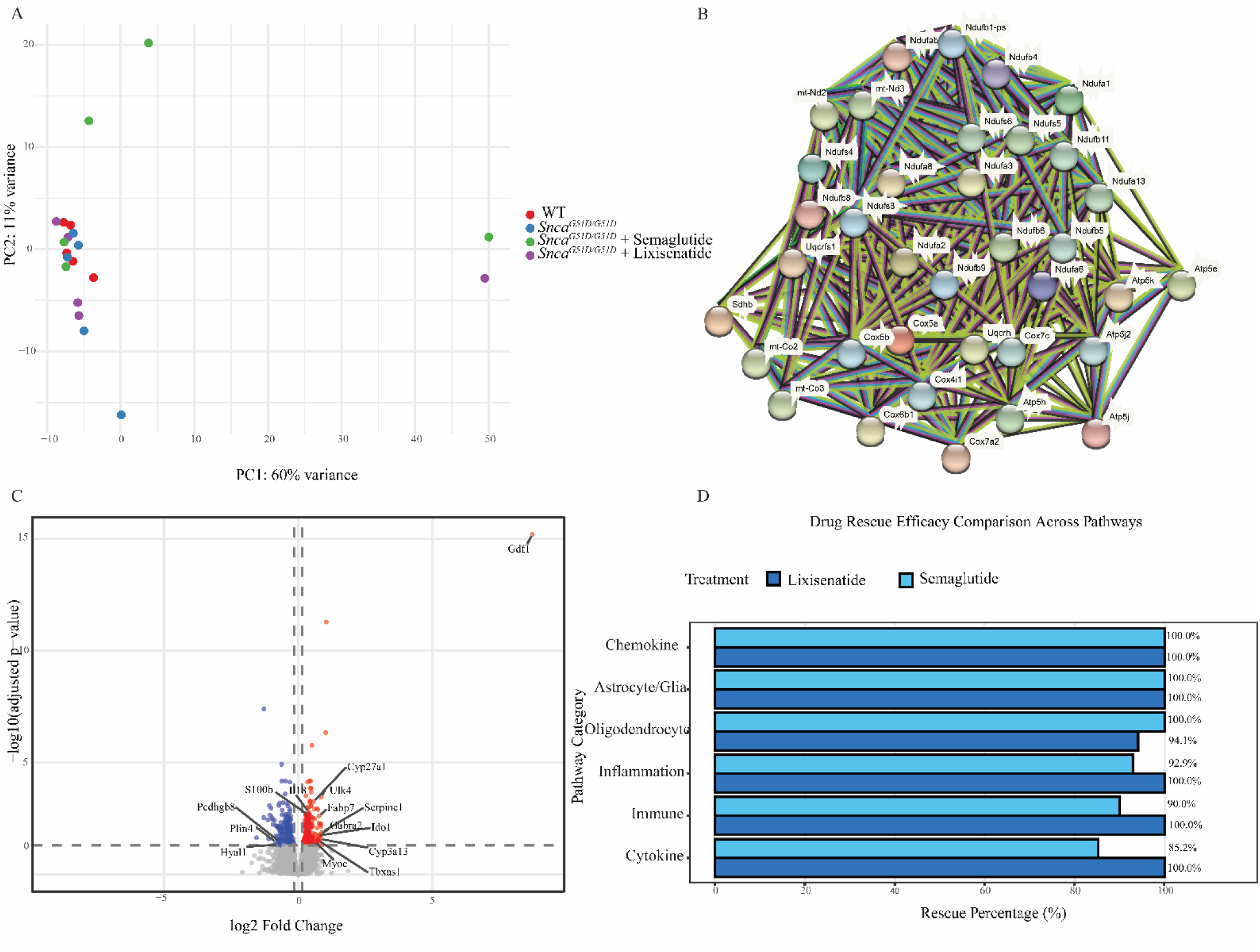
Bulk RNA Sequencing reveals changes across different treatment groups. (A) PCA plot for different treatment groups (B). String analysis reveals mitochondrial changes as the earliest hallmark of PD in *Snca^G51D/G51D^* mice (C). Volcano plot shows changes in inflammation-related DEGs (D). Rescue analysis of inflammation-related DEGs in Bulk RNA Sequencing data.

**Supplementary Fig. 5:**
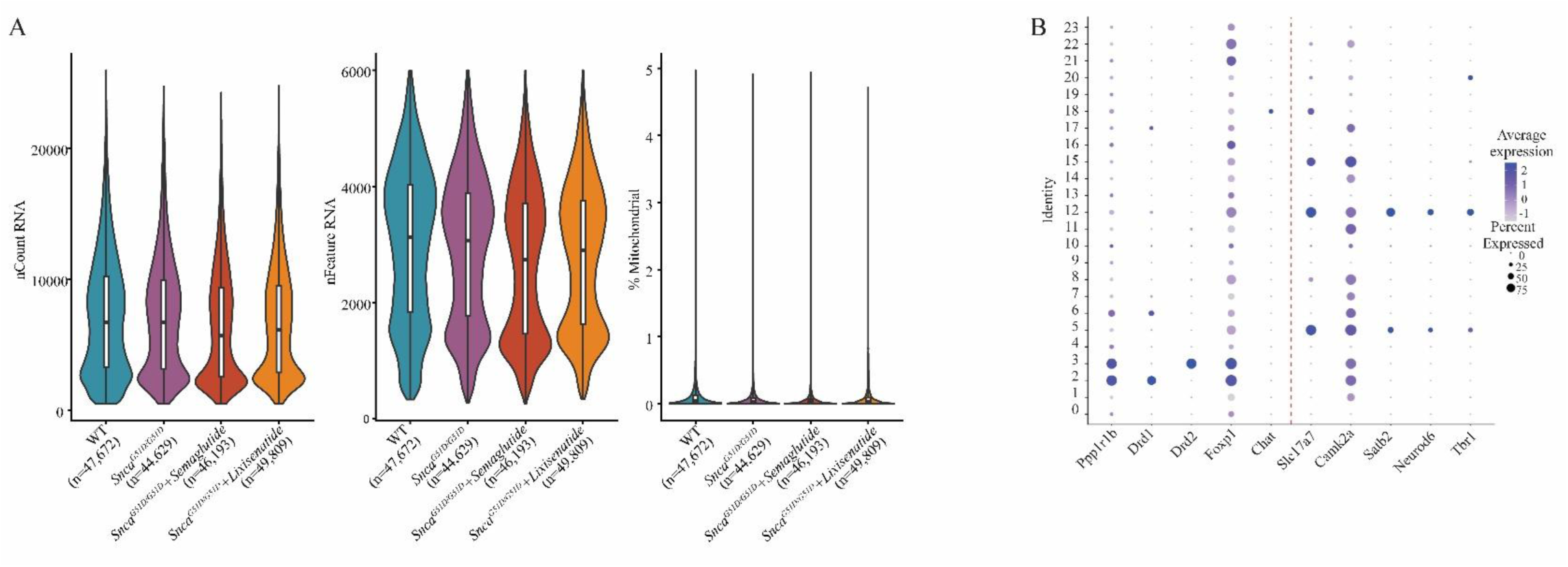
Quality control parameters for the single nuclei RNA Sequencing. (A) There were no significant differences in total RNA count across different treatment groups. (B) Top unique marker genes across different cell clusters in snRNA sequencing

**Supplementary Table 1:** Nuclei belonging to each sample which were included for analysis.

| Group | N_c<br>ells | nCount_<br>median | nCount<br>(meanÂ±SD<br>) | nFeature_<br>median | nFeature<br>(meanÂ±SD) | pct_mt_<br>median | %mt<br>(meanÂ±S<br>D) |
| --- | --- | --- | --- | --- | --- | --- | --- |
| WT1 | 112<br>54 | 4967 | 5254.2 Â±<br>3798.1 | 2517.5 | 2354.4 Â±<br>1235.5 | 0.01 | 0.05 Â±<br>0.1 |
| WT2 | 130<br>88 | 5501 | 6067.4 Â±<br>3598.7 | 2801 | 2774.2 Â±<br>1182.7 | 0.08 | 0.16 Â±<br>0.26 |
| WT3 | 110<br>19 | 8678 | 8739.5 Â±<br>4770 | 3698 | 3439.3 Â±<br>1286.7 | 0.03 | 0.08 Â±<br>0.18 |
| WT4 | 123<br>11 | 8791 | 8729.9 Â±<br>4935.8 | 3677 | 3394.1 Â±<br>1300.7 | 0.07 | 0.13 Â±<br>0.21 |
| *Snca*<sup>G51D/G51D</sup> 1 | 116<br>52 | 7077 | 7326.3 Â±<br>4545.1 | 3075 | 2911.1 Â±<br>1254.2 | 0.02 | 0.07 Â±<br>0.2 |
| *Snca*<sup>G51D/G51D</sup> 2 | 135<br>88 | 5852 | 6168.7 Â±<br>3748.9 | 2827 | 2709.1 Â±<br>1177.4 | 0.03 | 0.06 Â±<br>0.14 |
| *Snca*<sup>G51D/G51D</sup> 3 | 986<br>3 | 7684 | 7780.9 Â±<br>4559 | 3404 | 3189.4 Â±<br>1297.2 | 0.07 | 0.12 Â±<br>0.23 |
| *Snca*<sup>G51D/G51D</sup> 4 | 952<br>6 | 7031 | 7421.3 Â±<br>4773.8 | 3137.5 | 2974.7 Â±<br>1314 | 0.02 | 0.06 Â±<br>0.15 |
| *Snca*<sup>G51D/G51D</sup><br>+Semaglutide 1 | 106<br>33 | 5058 | 5663.7 Â±<br>3864.4 | 2522 | 2474.5 Â±<br>1218.1 | 0.02 | 0.07 Â±<br>0.2 |
| *Snca*<sup>G51D/G51D</sup><br>+Semaglutide 2 | 136<br>12 | 5588 | 6429.9 Â±<br>4240.5 | 2717 | 2728.6 Â±<br>1276.6 | 0.03 | 0.08 Â±<br>0.22 |
| *Snca*<sup>G51D/G51D</sup><br>+Semaglutide 3 | 107<br>88 | 7297 | 7503.7 Â±<br>4693.6 | 3177 | 2960 Â±<br>1281.7 | 0.03 | 0.08 Â±<br>0.2 |
| *Snca*<sup>G51D/G51D</sup><br>+Semaglutide 4 | 111<br>60 | 4575 | 5923.5 Â±<br>4092.5 | 2314 | 2508.5 Â±<br>1251.1 | 0.01 | 0.06 Â±<br>0.18 |
| *Snca*<sup>G51D/G51D</sup><br>+Lixisenatide 1 | 112<br>54 | 6989.5 | 7311.5 Â±<br>4514.5 | 3141 | 2971.8 Â±<br>1271 | 0.03 | 0.09 Â±<br>0.23 |
| *Snca*<sup>G51D/G51D</sup><br>+Lixisenatide 2 | 123<br>30 | 4927.5 | 5833.1 Â±<br>3882.4 | 2452 | 2518.2 Â±<br>1190.8 | 0.04 | 0.09 Â±<br>0.22 |
| *Snca*<sup>G51D/G51D</sup><br>+Lixisenatide 3 | 150<br>54 | 5964 | 6237.4 Â±<br>3791.1 | 2850 | 2684.4 Â±<br>1140.4 | 0.03 | 0.09 Â±<br>0.22 |
| *Snca*<sup>G51D/G51D</sup><br>+Lixisenatide 4 | 111<br>71 | 7258 | 7669.9 Â±<br>4786.8 | 3262 | 3046.5 Â±<br>1287.9 | 0.03 | 0.08 Â±<br>0.19 |

**Supplementary Table 2:**
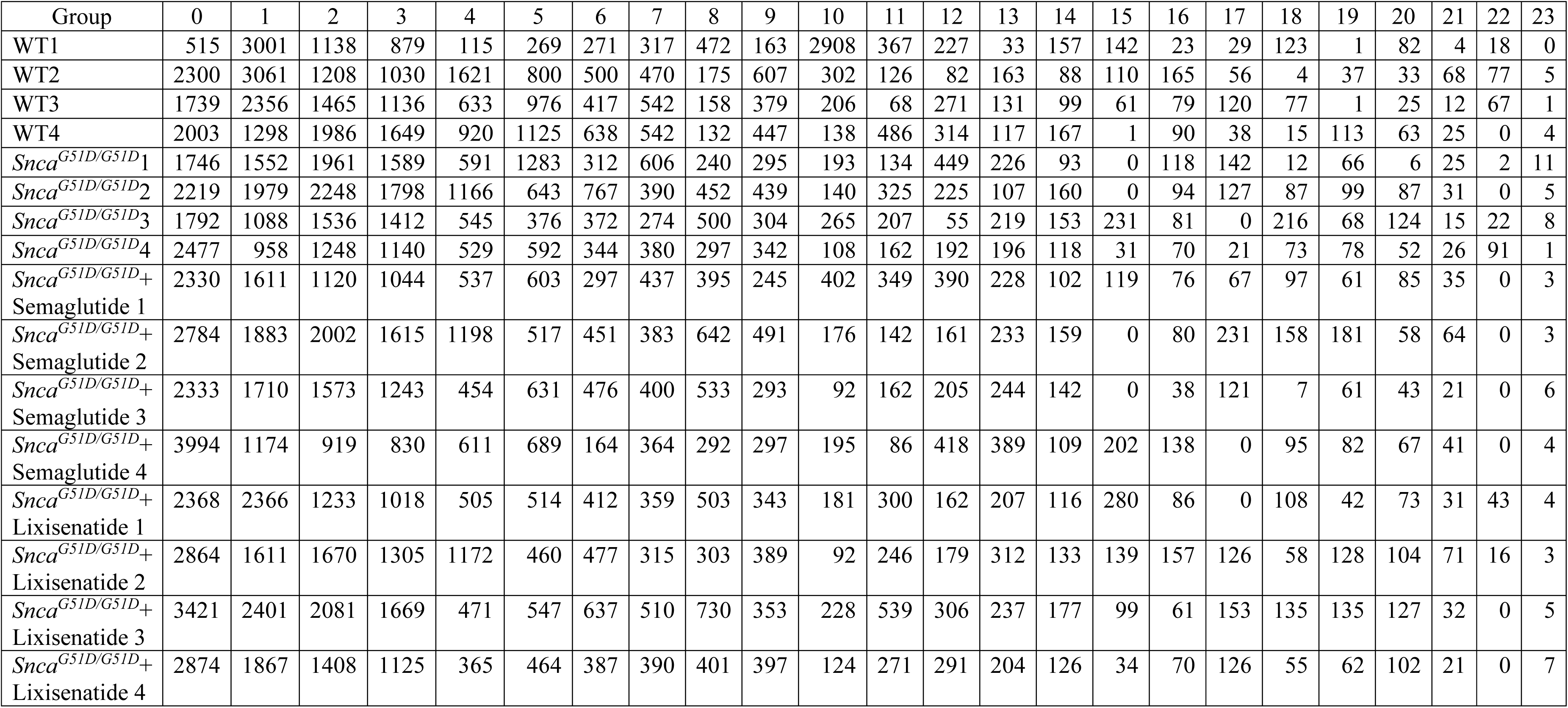
Cell-type composition examined at the sample and cluster level across groups.

| Group | 0 | 1 | 2 | 3 | 4 | 5 | 6 | 7 | 8 | 9 | 10 | 11 | 12 | 13 | 14 | 15 | 16 | 17 | 18 | 19 | 20 | 21 | 22 | 23 |
| --- | --- | --- | --- | --- | --- | --- | --- | --- | --- | --- | --- | --- | --- | --- | --- | --- | --- | --- | --- | --- | --- | --- | --- | --- |
| WT1 | 515 | 3001 | 1138 | 879 | 115 | 269 | 271 | 317 | 472 | 163 | 2908 | 367 | 227 | 33 | 157 | 142 | 23 | 29 | 123 | 1 | 82 | 4 | 18 | 0 |
| WT2 | 2300 | 3061 | 1208 | 1030 | 1621 | 800 | 500 | 470 | 175 | 607 | 302 | 126 | 82 | 163 | 88 | 110 | 165 | 56 | 4 | 37 | 33 | 68 | 77 | 5 |
| WT3 | 1739 | 2356 | 1465 | 1136 | 633 | 976 | 417 | 542 | 158 | 379 | 206 | 68 | 271 | 131 | 99 | 61 | 79 | 120 | 77 | 1 | 25 | 12 | 67 | 1 |
| WT4 | 2003 | 1298 | 1986 | 1649 | 920 | 1125 | 638 | 542 | 132 | 447 | 138 | 486 | 314 | 117 | 167 | 1 | 90 | 38 | 15 | 113 | 63 | 25 | 0 | 4 |
| *Snca*<sup>G51D/G51D</sup>1 | 1746 | 1552 | 1961 | 1589 | 591 | 1283 | 312 | 606 | 240 | 295 | 193 | 134 | 449 | 226 | 93 | 0 | 118 | 142 | 12 | 66 | 6 | 25 | 2 | 11 |
| *Snca*<sup>G51D/G51D</sup>2 | 2219 | 1979 | 2248 | 1798 | 1166 | 643 | 767 | 390 | 452 | 439 | 140 | 325 | 225 | 107 | 160 | 0 | 94 | 127 | 87 | 99 | 87 | 31 | 0 | 5 |
| *Snca*<sup>G51D/G51D</sup>3 | 1792 | 1088 | 1536 | 1412 | 545 | 376 | 372 | 274 | 500 | 304 | 265 | 207 | 55 | 219 | 153 | 231 | 81 | 0 | 216 | 68 | 124 | 15 | 22 | 8 |
| *Snca*<sup>G51D/G51D</sup>4 | 2477 | 958 | 1248 | 1140 | 529 | 592 | 344 | 380 | 297 | 342 | 108 | 162 | 192 | 196 | 118 | 31 | 70 | 21 | 73 | 78 | 52 | 26 | 91 | 1 |
| *Snca*<sup>G51D/G51D</sup>+<br>Semaglutide 1 | 2330 | 1611 | 1120 | 1044 | 537 | 603 | 297 | 437 | 395 | 245 | 402 | 349 | 390 | 228 | 102 | 119 | 76 | 67 | 97 | 61 | 85 | 35 | 0 | 3 |
| *Snca*<sup>G51D/G51D</sup>+<br>Semaglutide 2 | 2784 | 1883 | 2002 | 1615 | 1198 | 517 | 451 | 383 | 642 | 491 | 176 | 142 | 161 | 233 | 159 | 0 | 80 | 231 | 158 | 181 | 58 | 64 | 0 | 3 |
| *Snca*<sup>G51D/G51D</sup>+<br>Semaglutide 3 | 2333 | 1710 | 1573 | 1243 | 454 | 631 | 476 | 400 | 533 | 293 | 92 | 162 | 205 | 244 | 142 | 0 | 38 | 121 | 7 | 61 | 43 | 21 | 0 | 6 |
| *Snca*<sup>G51D/G51D</sup>+<br>Semaglutide 4 | 3994 | 1174 | 919 | 830 | 611 | 689 | 164 | 364 | 292 | 297 | 195 | 86 | 418 | 389 | 109 | 202 | 138 | 0 | 95 | 82 | 67 | 41 | 0 | 4 |
| *Snca*<sup>G51D/G51D</sup>+<br>Lixisenatide 1 | 2368 | 2366 | 1233 | 1018 | 505 | 514 | 412 | 359 | 503 | 343 | 181 | 300 | 162 | 207 | 116 | 280 | 86 | 0 | 108 | 42 | 73 | 31 | 43 | 4 |
| *Snca*<sup>G51D/G51D</sup>+<br>Lixisenatide 2 | 2864 | 1611 | 1670 | 1305 | 1172 | 460 | 477 | 315 | 303 | 389 | 92 | 246 | 179 | 312 | 133 | 139 | 157 | 126 | 58 | 128 | 104 | 71 | 16 | 3 |
| *Snca*<sup>G51D/G51D</sup>+<br>Lixisenatide 3 | 3421 | 2401 | 2081 | 1669 | 471 | 547 | 637 | 510 | 730 | 353 | 228 | 539 | 306 | 237 | 177 | 99 | 61 | 153 | 135 | 135 | 127 | 32 | 0 | 5 |
| *Snca*<sup>G51D/G51D</sup>+<br>Lixisenatide 4 | 2874 | 1867 | 1408 | 1125 | 365 | 464 | 387 | 390 | 401 | 397 | 124 | 271 | 291 | 204 | 126 | 34 | 70 | 126 | 55 | 62 | 102 | 21 | 0 | 7 |

**Supplement Table 3.** Reagents and resources used for this paper.

| Reagent or resource | Source | Identifier |
| --- | --- | --- |
| **Antibodies** |  |  |
| Anti-Alpha-Synuclein antibody (clone 42) | BD | RRID:<br>AB_398107 |
| Anti-Vinculin | Sigma-Aldrich | RRID:<br>AB_477629 |
| Anti-GFAP | Novus Biologicals | RRID:<br>AB_829022 |
| Anti-Iba1 | Wako Chemicals | RRID:<br>AB_839504 |
| Donkey anti-goat IgG (H+L) Highly Cross-Adsorbed Secondary Antibody, Alexa Fluor™ 488 | Thermo Fisher Scientific | RRID:<br>AB_2534102 |
| Donkey anti-Rabbit IgG (H+L) Highly Cross-Adsorbed Secondary Antibody, Alexa Fluor™ 555 | Thermo Fisher Scientific | RRID:<br>AB_162543 |
| **Other Reagents** |  |  |
| VECTASHIELD® PLUS Antifade Mounting Medium with DAPI | Vector Laboratories | H-2000-2 |
| ECL | GE Healthcare | RPN2236 |
| Optimal Cut Temperature Compound | VWR | 25608-930 |
| **Drugs** |  |  |
| Semaglutide | MedKoo | Cat# 581055 |
| Lixisenatide | MedChemExpress | HY-P0119 |
| **Kits** |  |  |
| Glucose Estimation Kit | Elabscience | E-BC-K234-S |
| Cholesterol species estimation kit | Sigma-Aldrich | MAK331 |
| Triglycerides estimation kit | Elabscience | E-BC-K238 |
| Insulin Measurement Kit | Crystal Chem | Cat#62100 |

## Supplementary Text 1

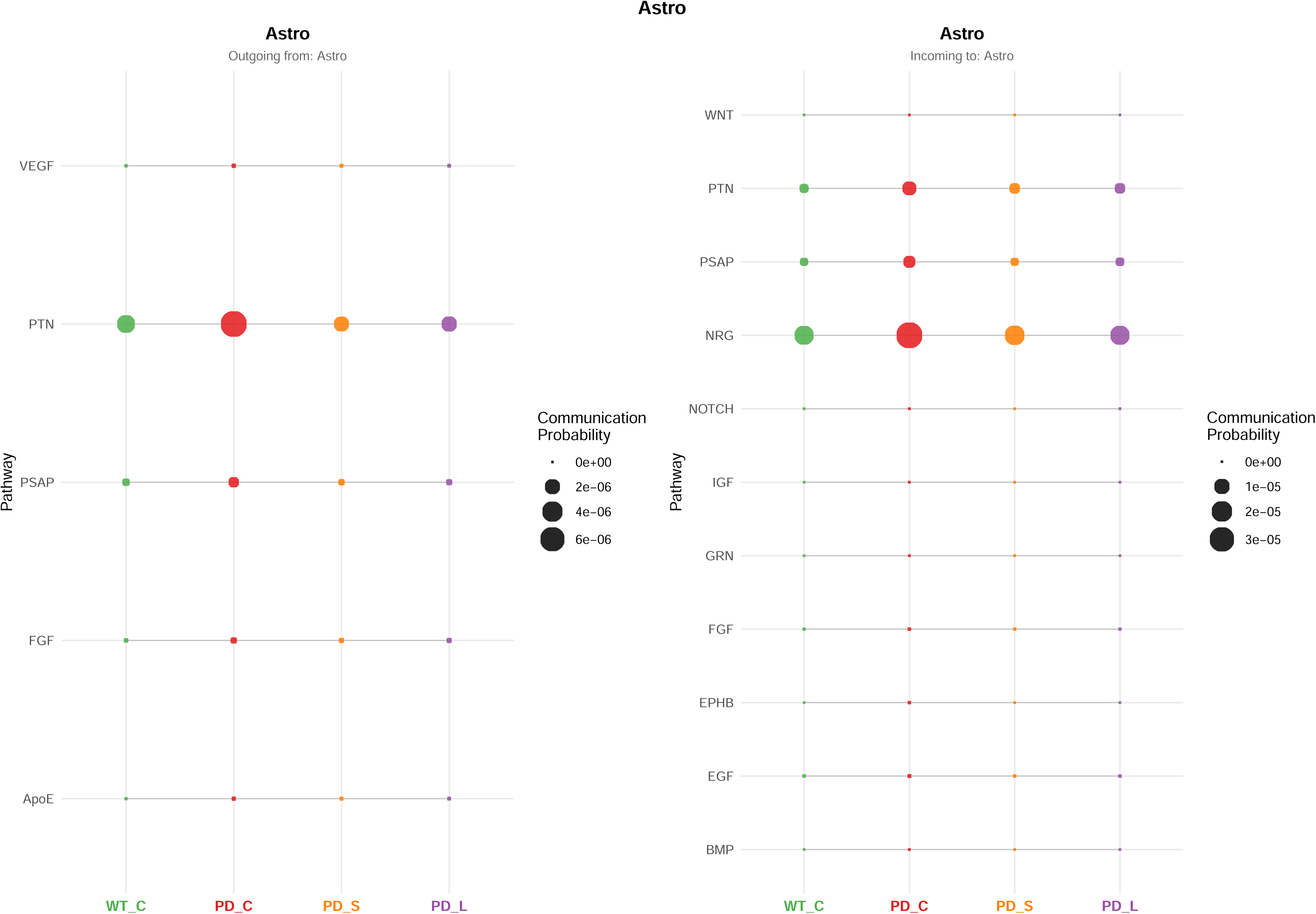

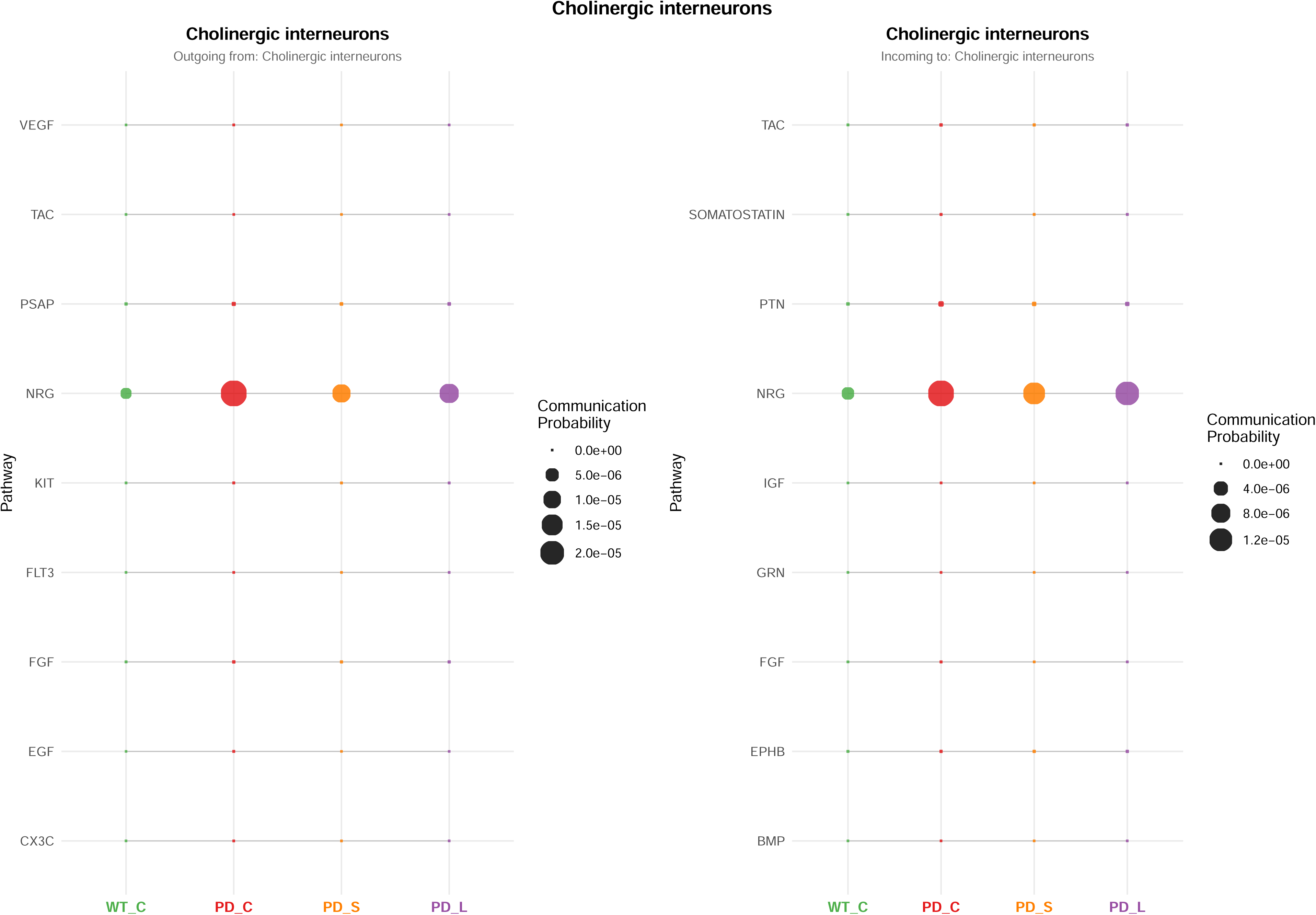

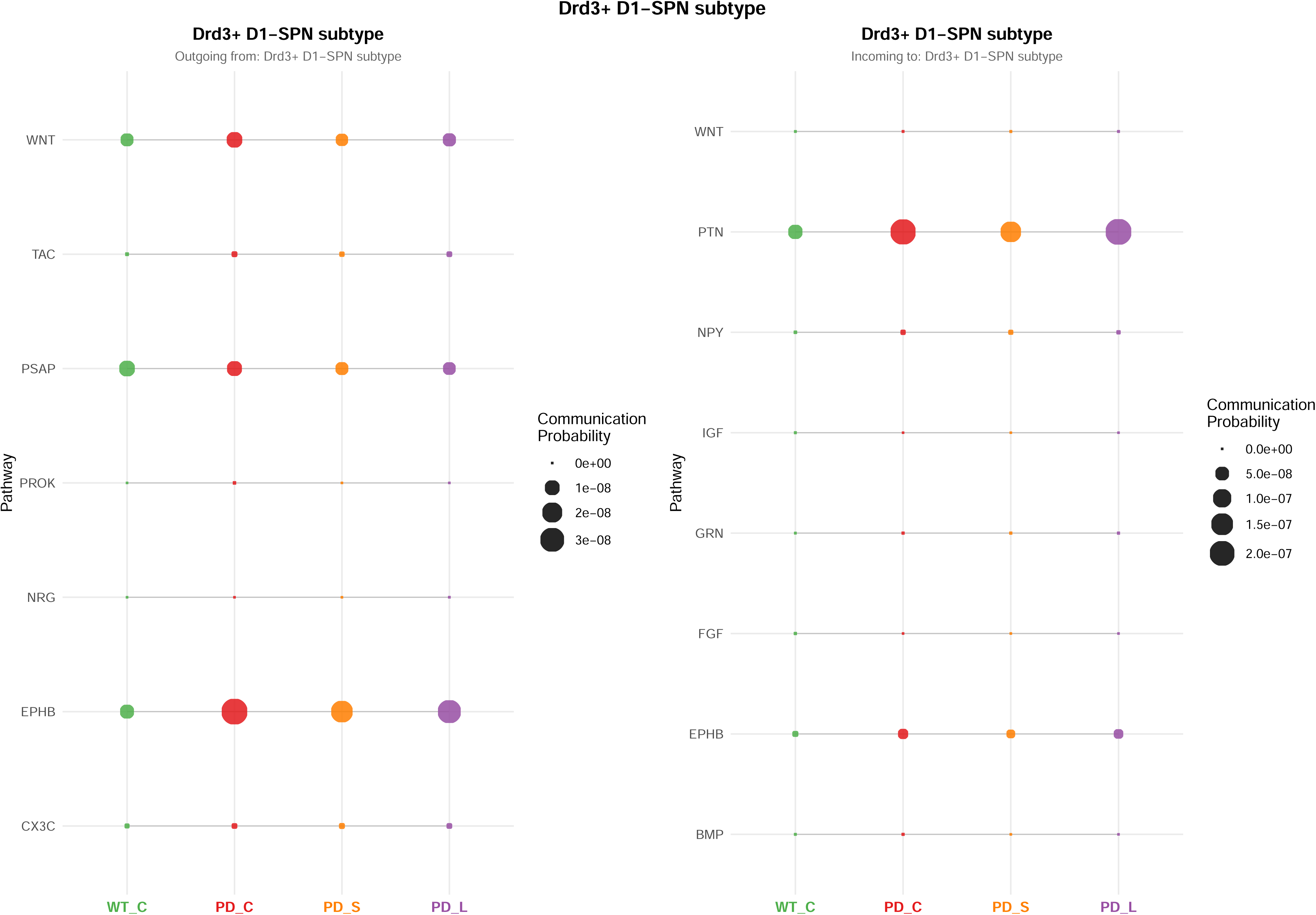

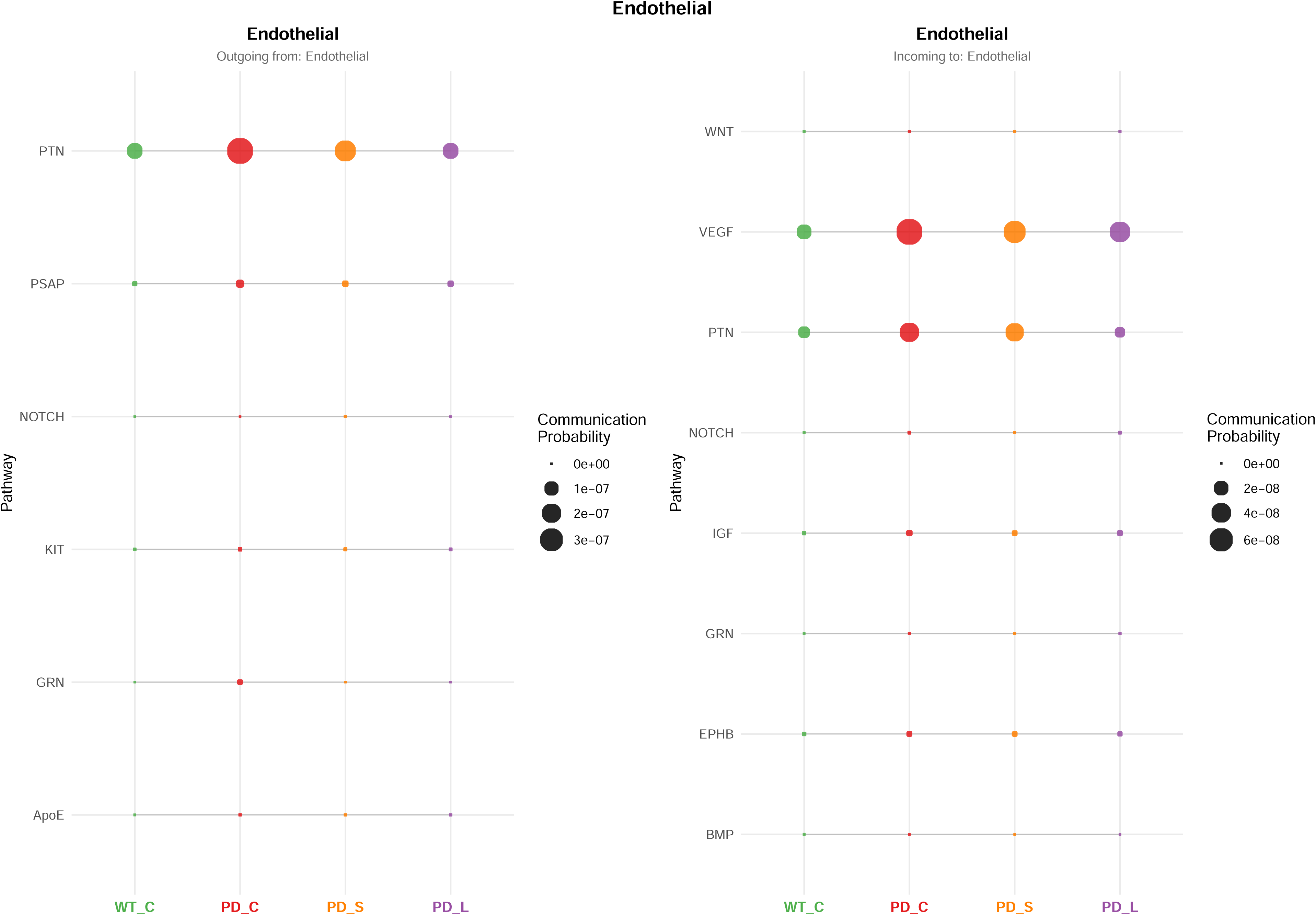

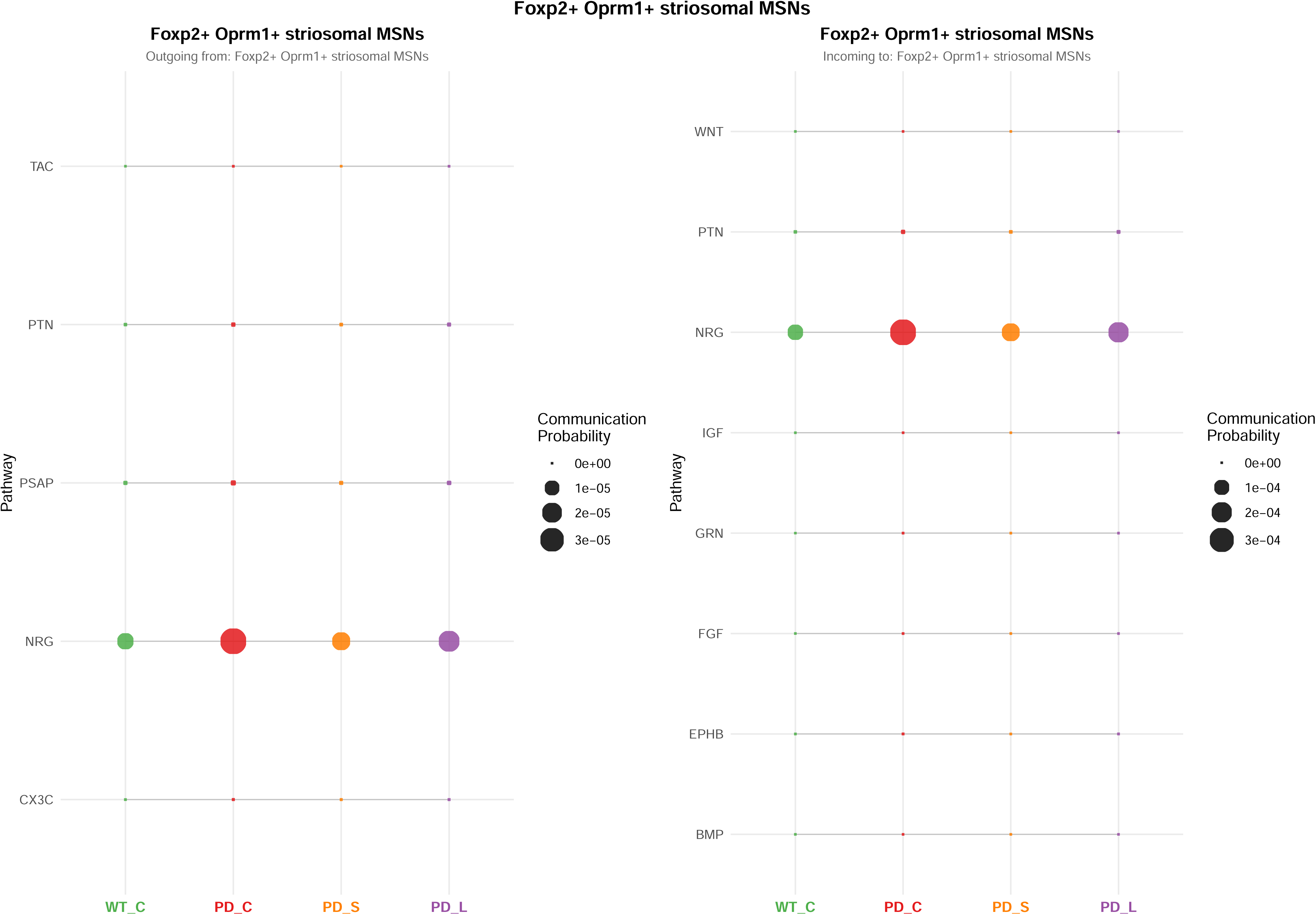

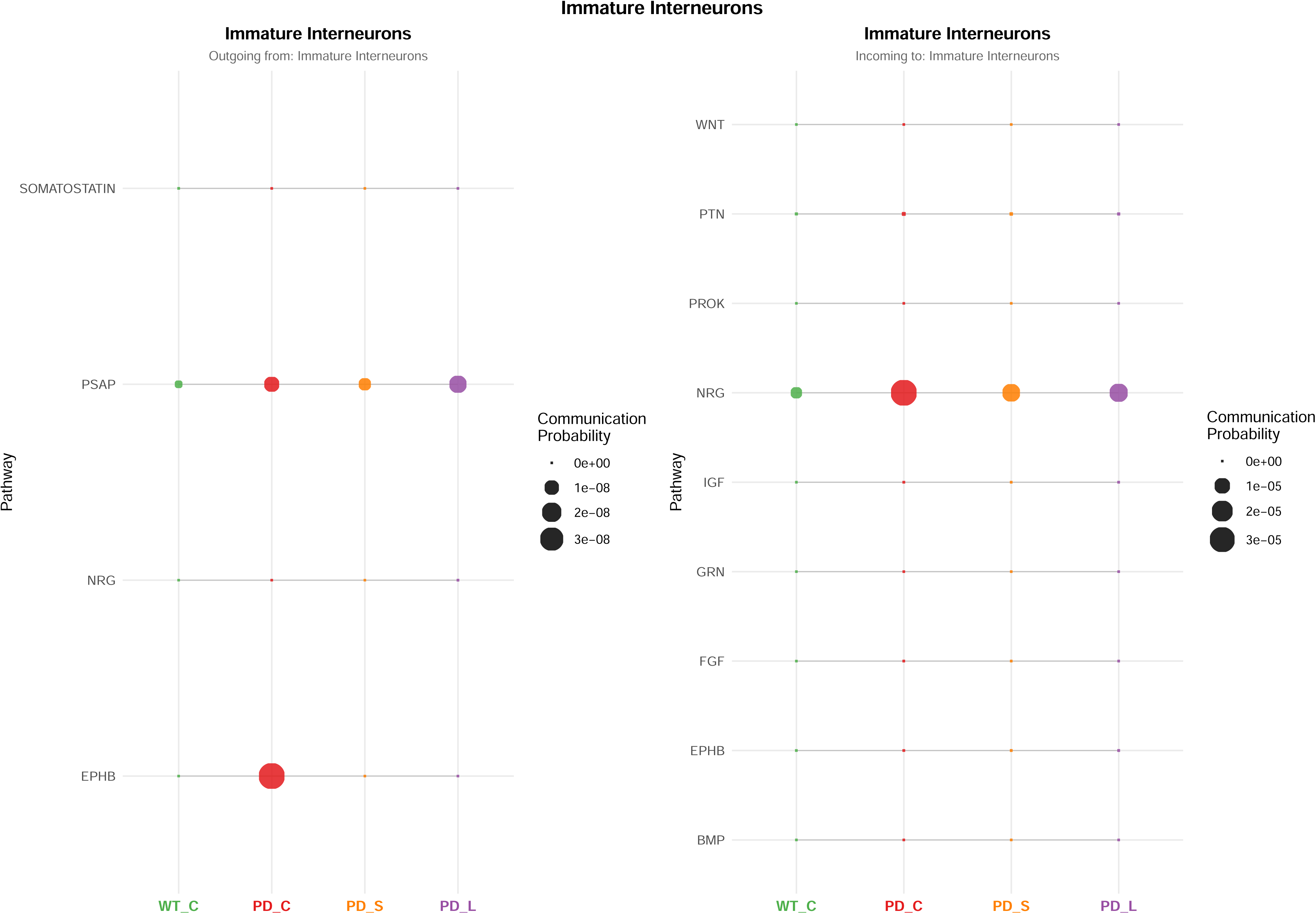

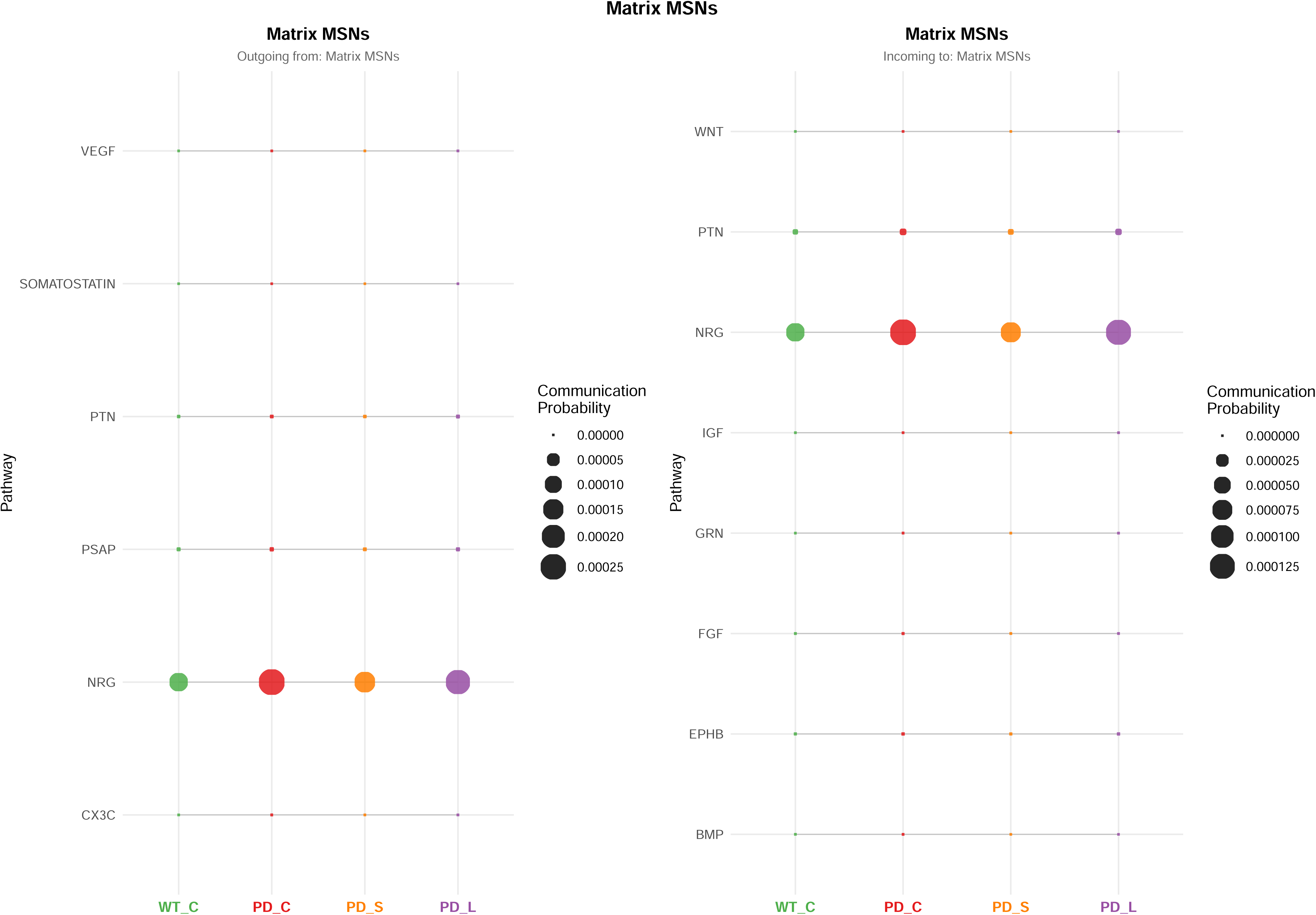

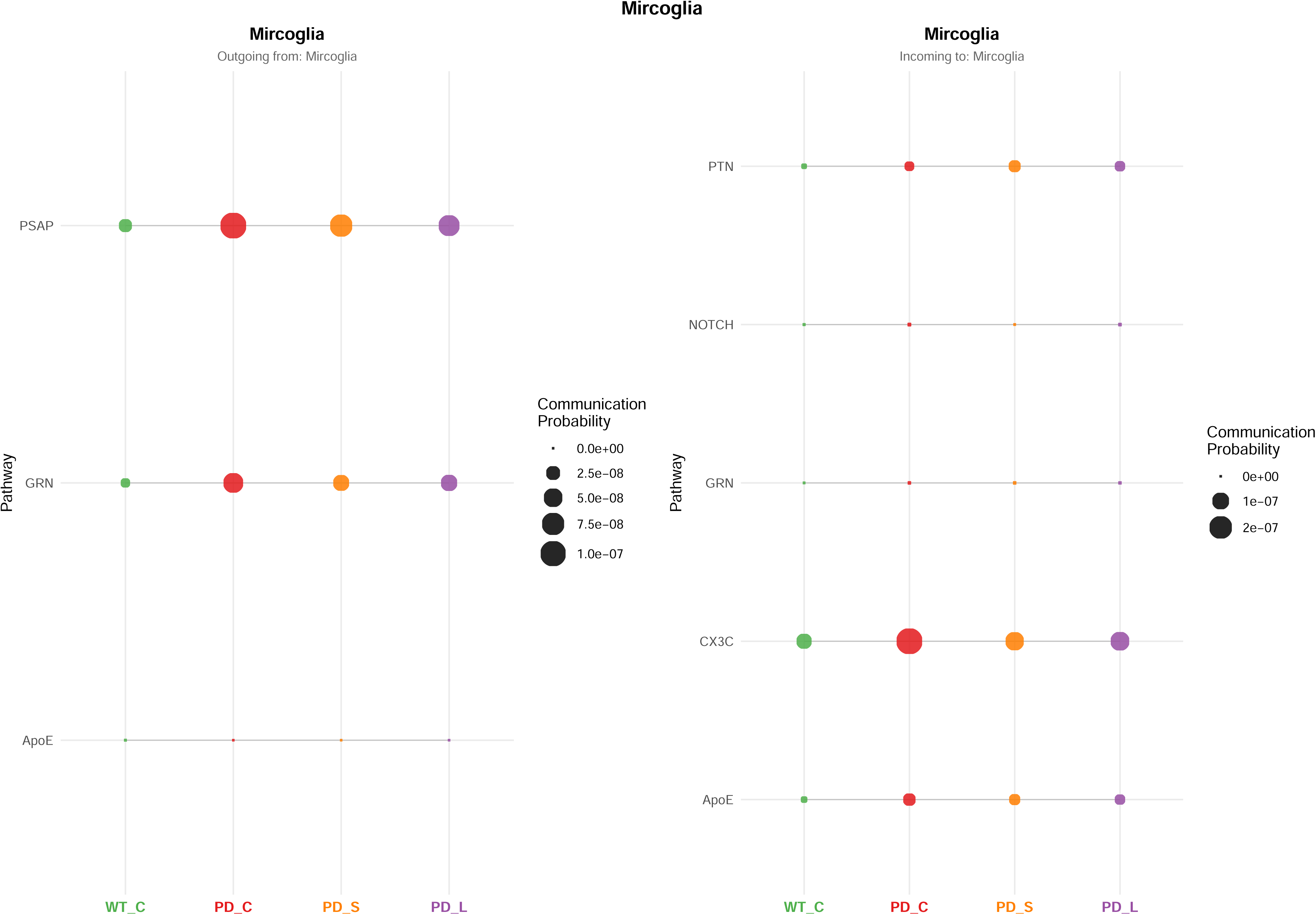

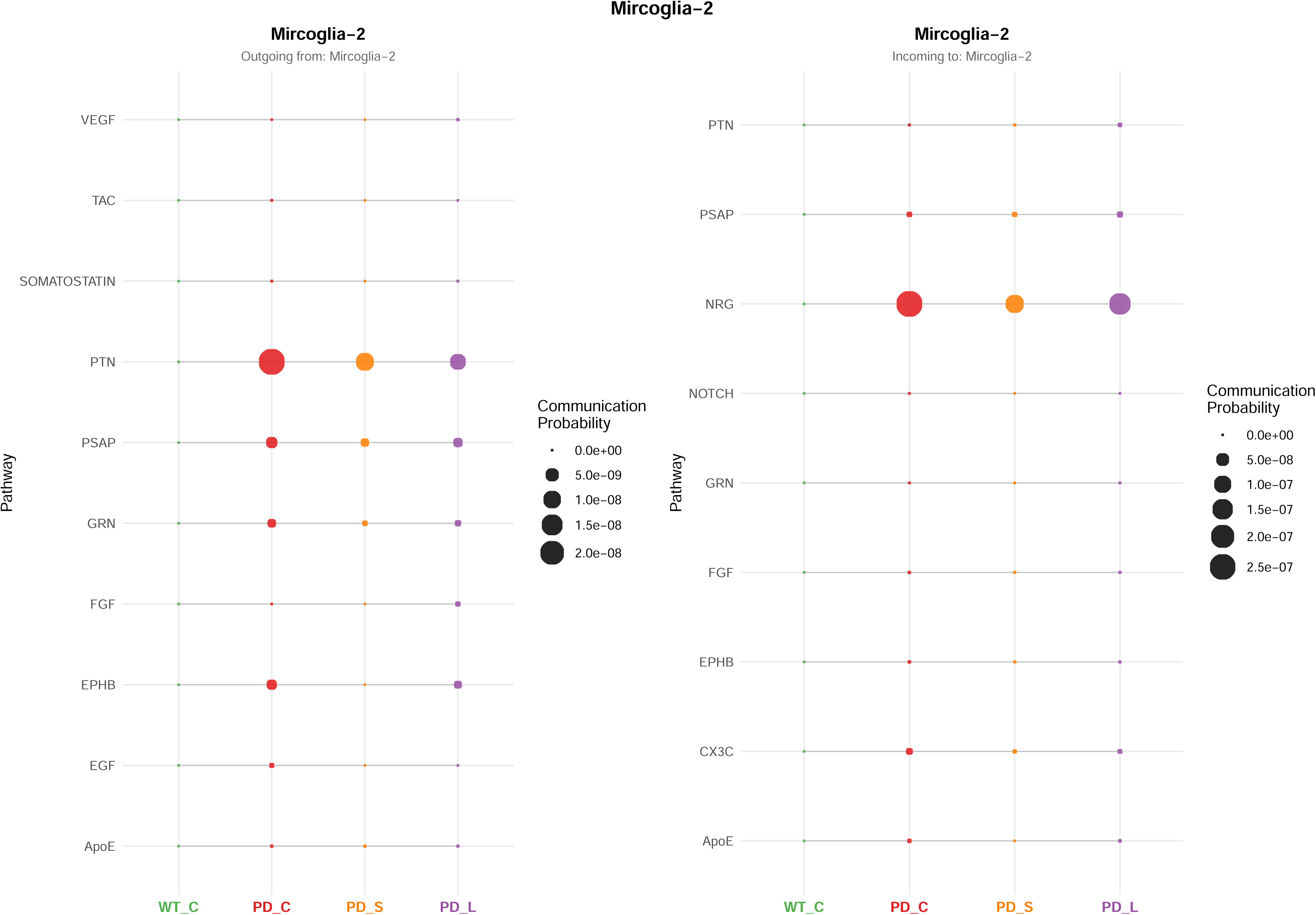

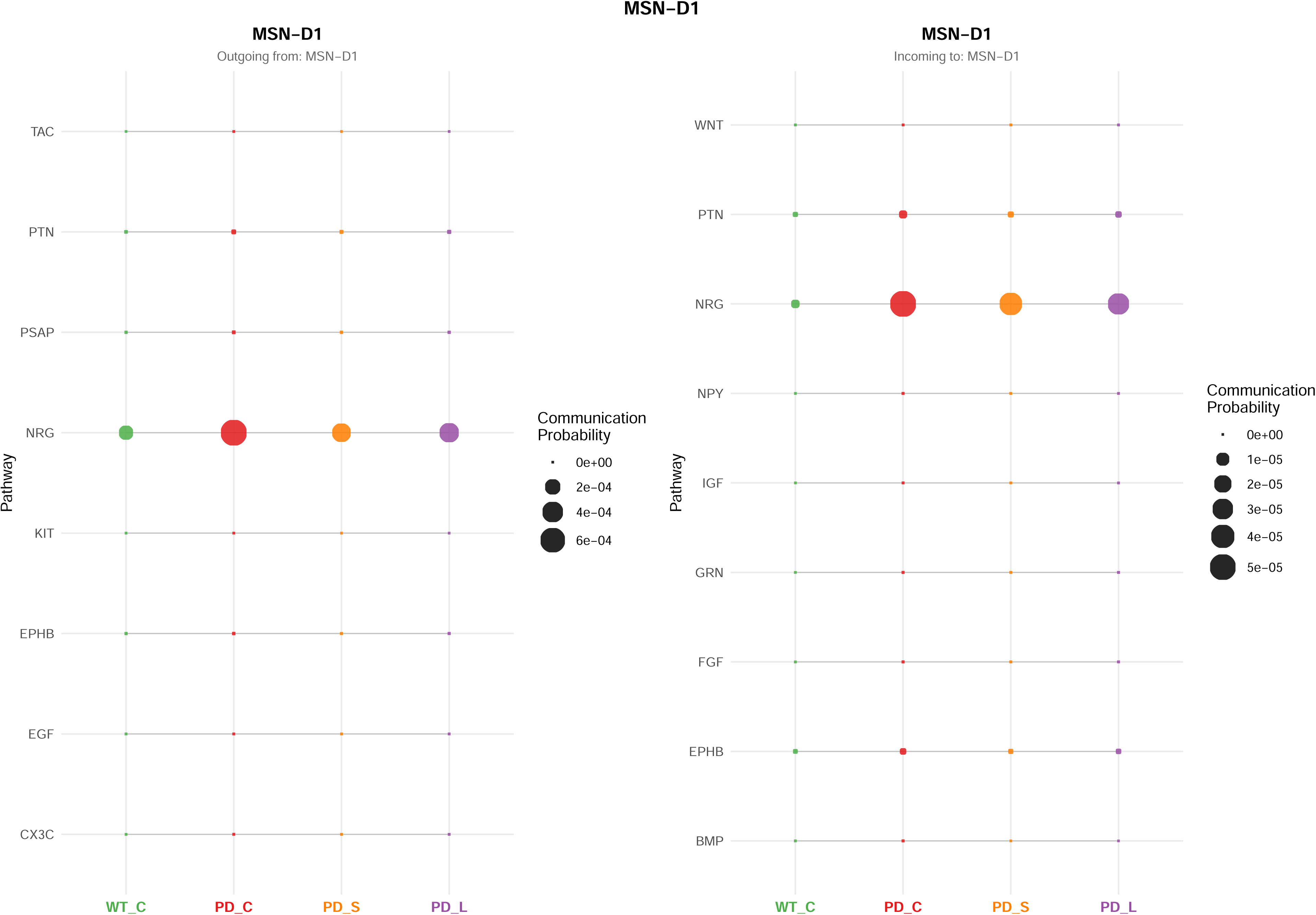

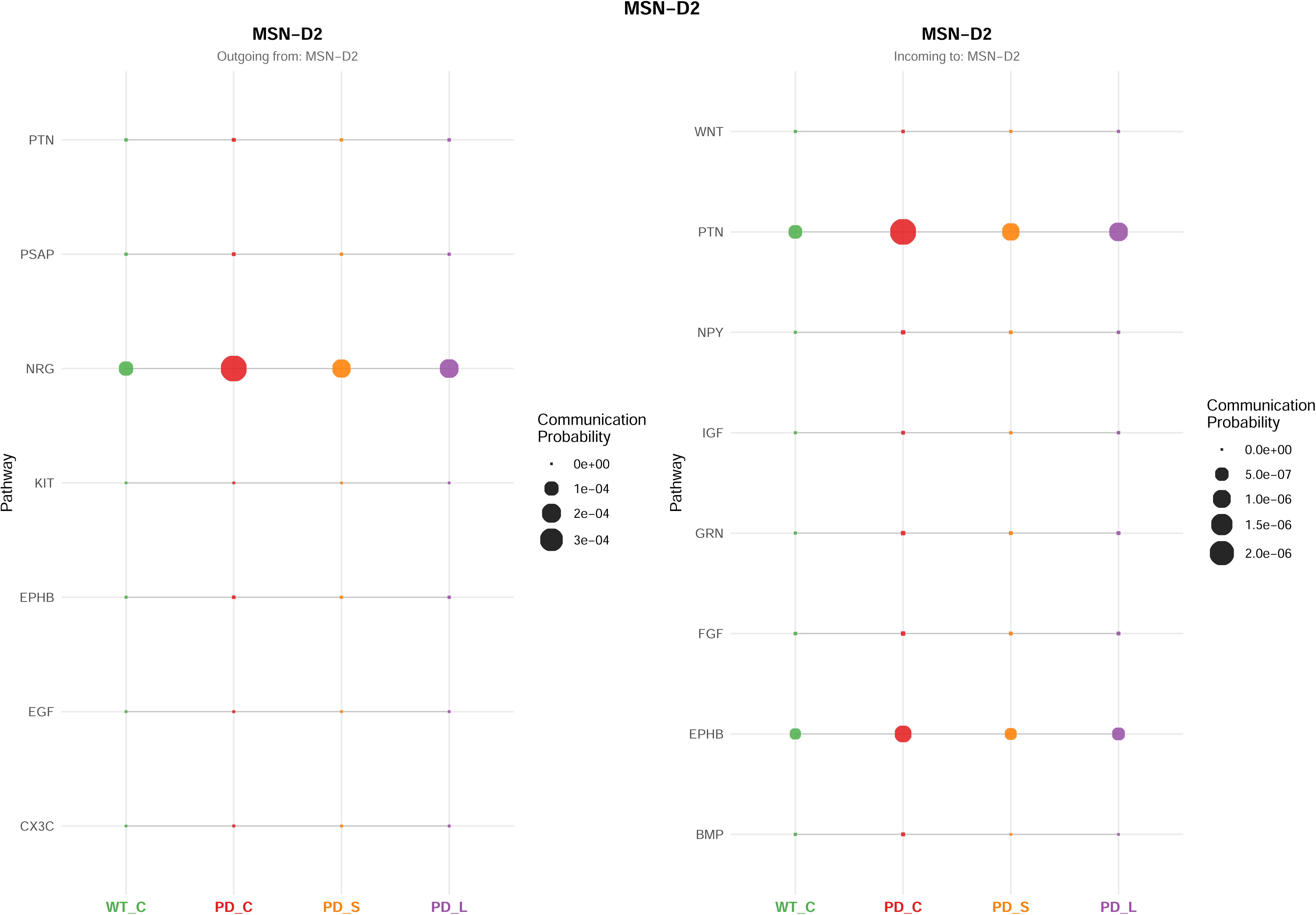

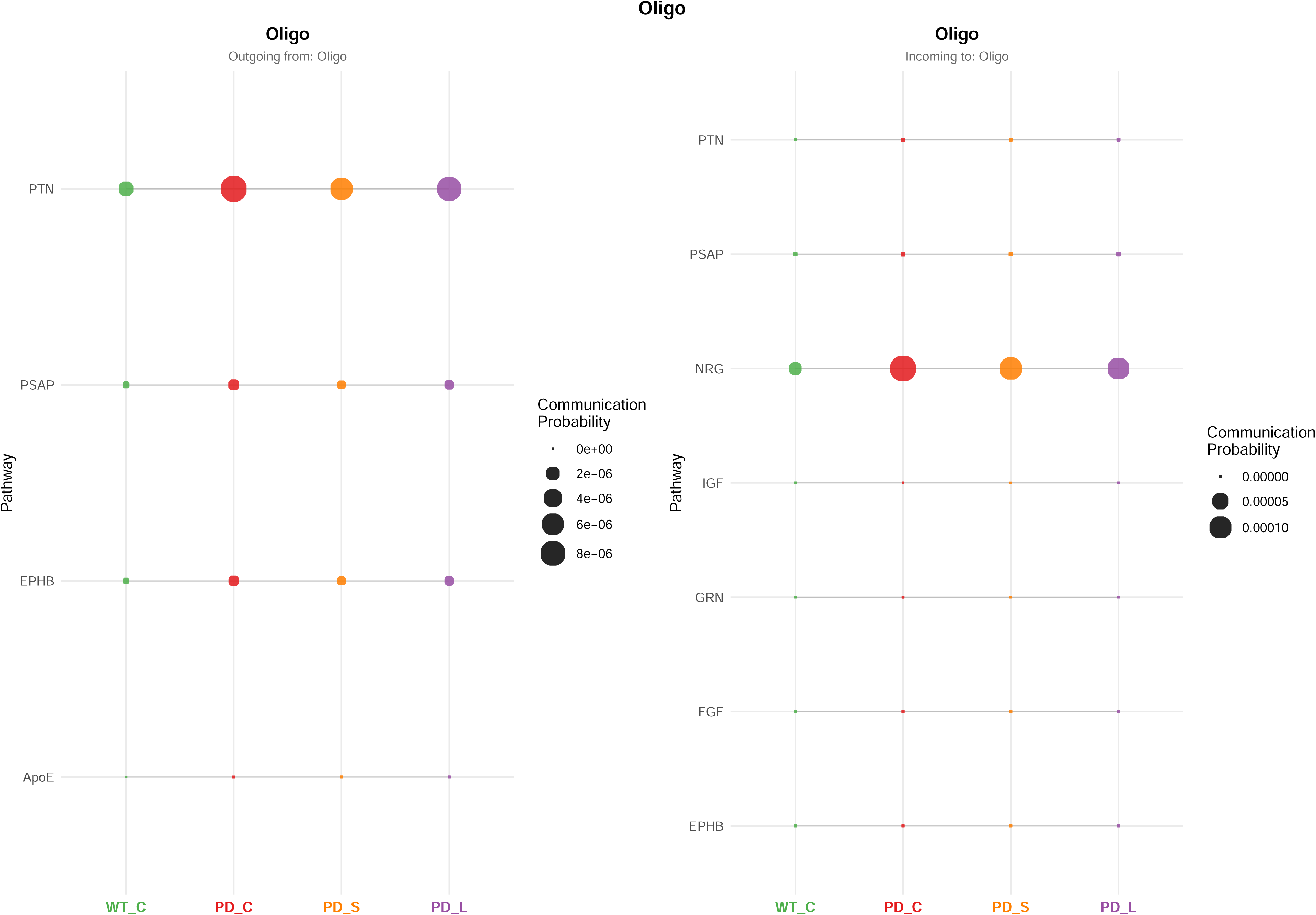

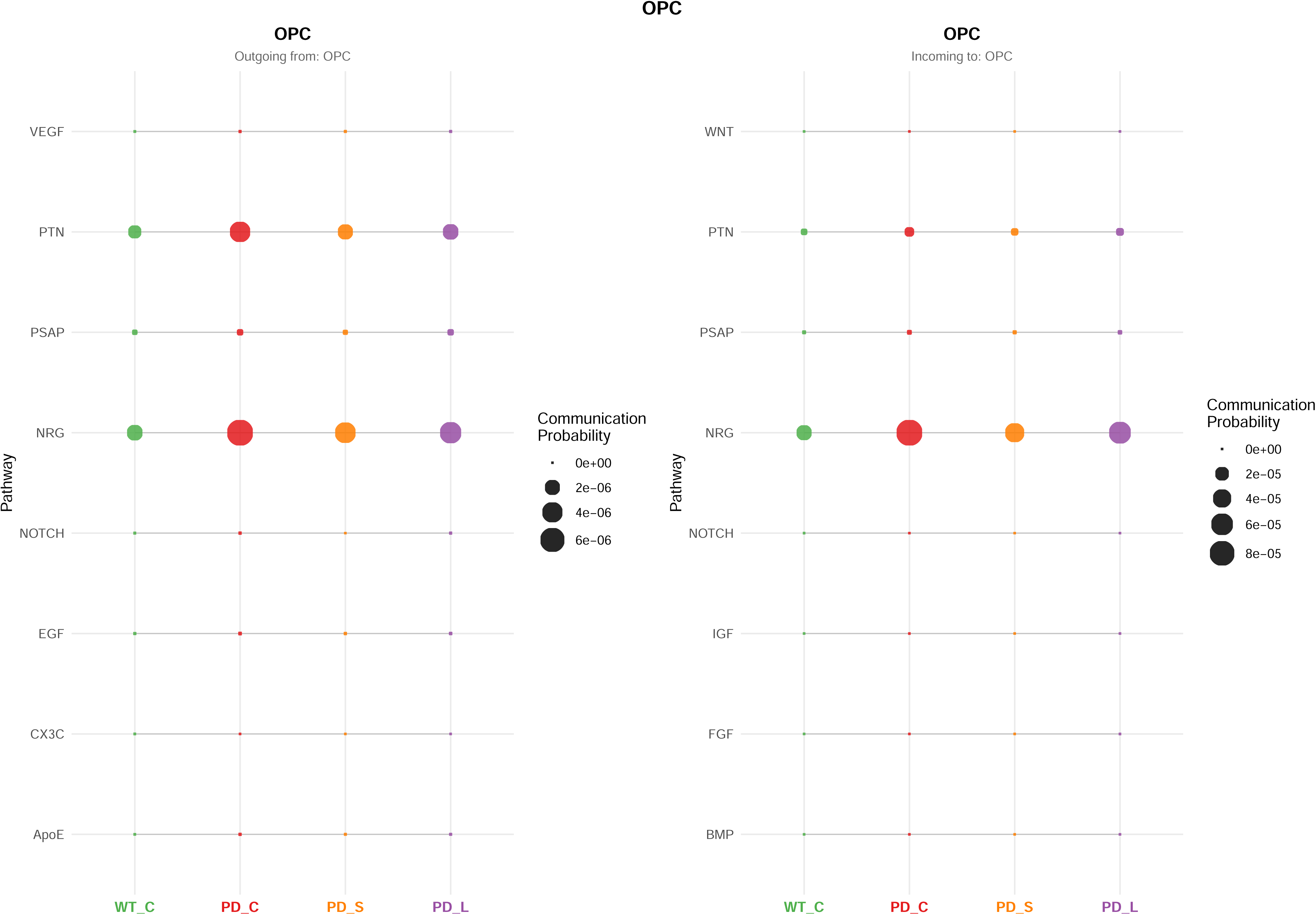

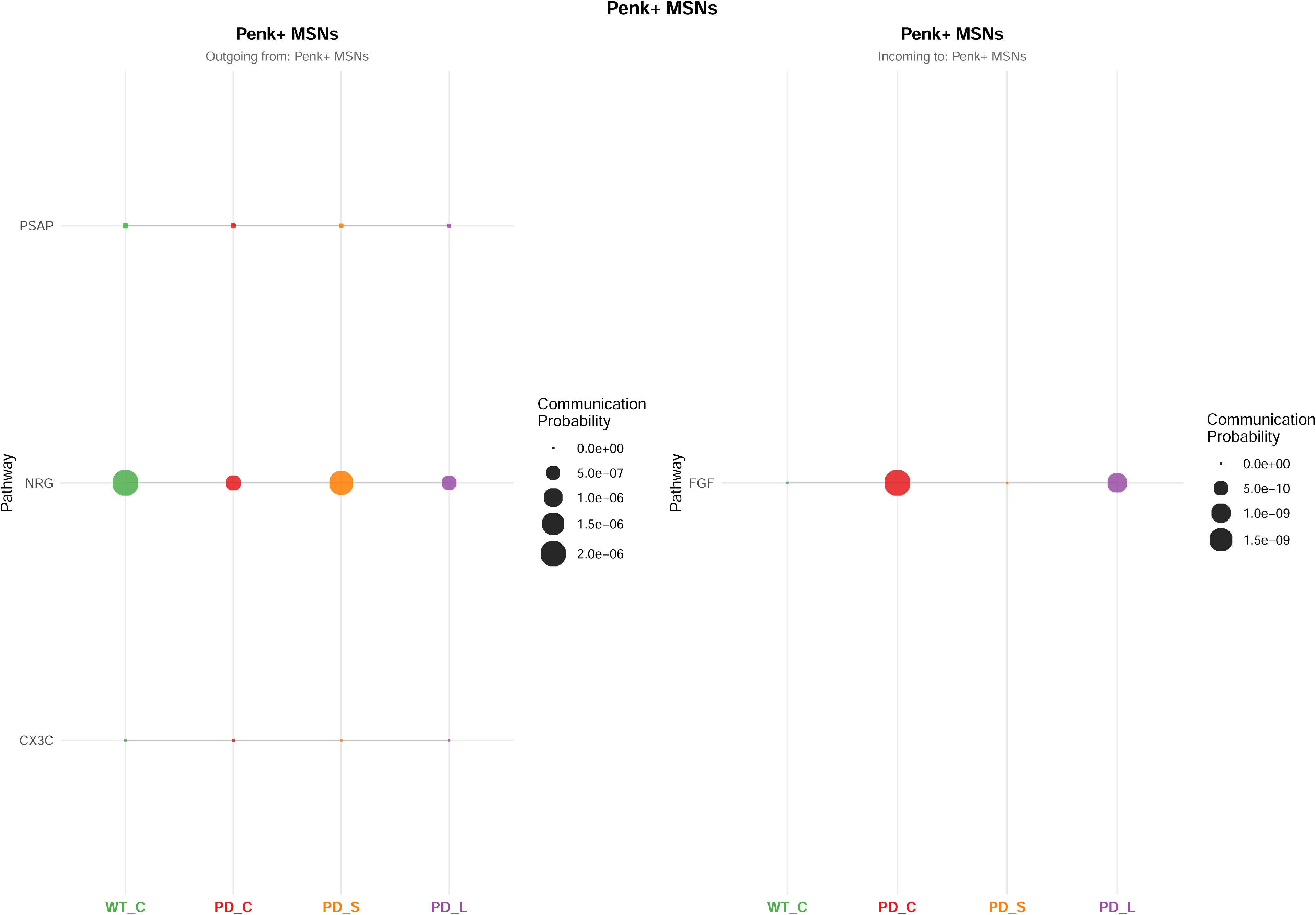

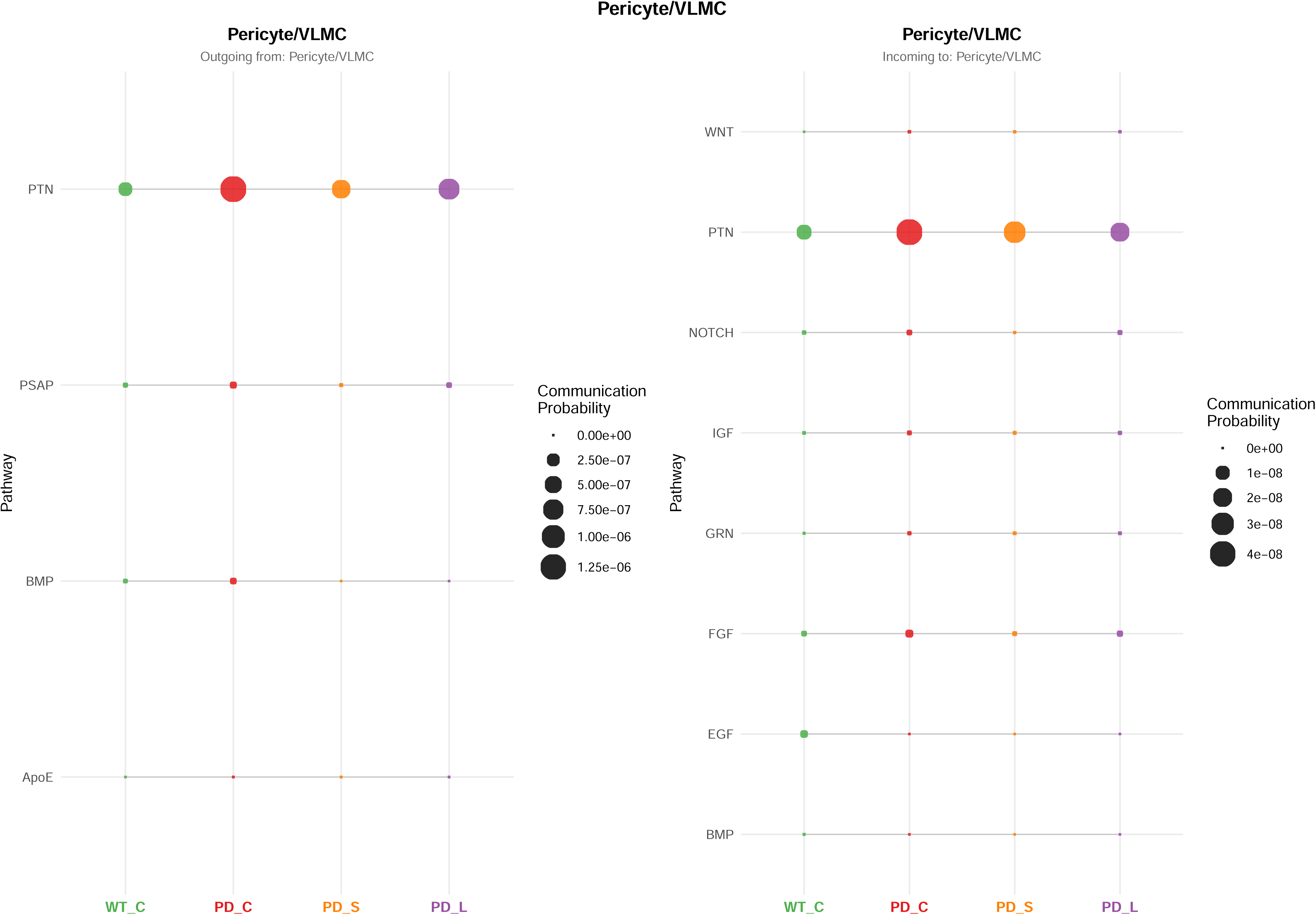

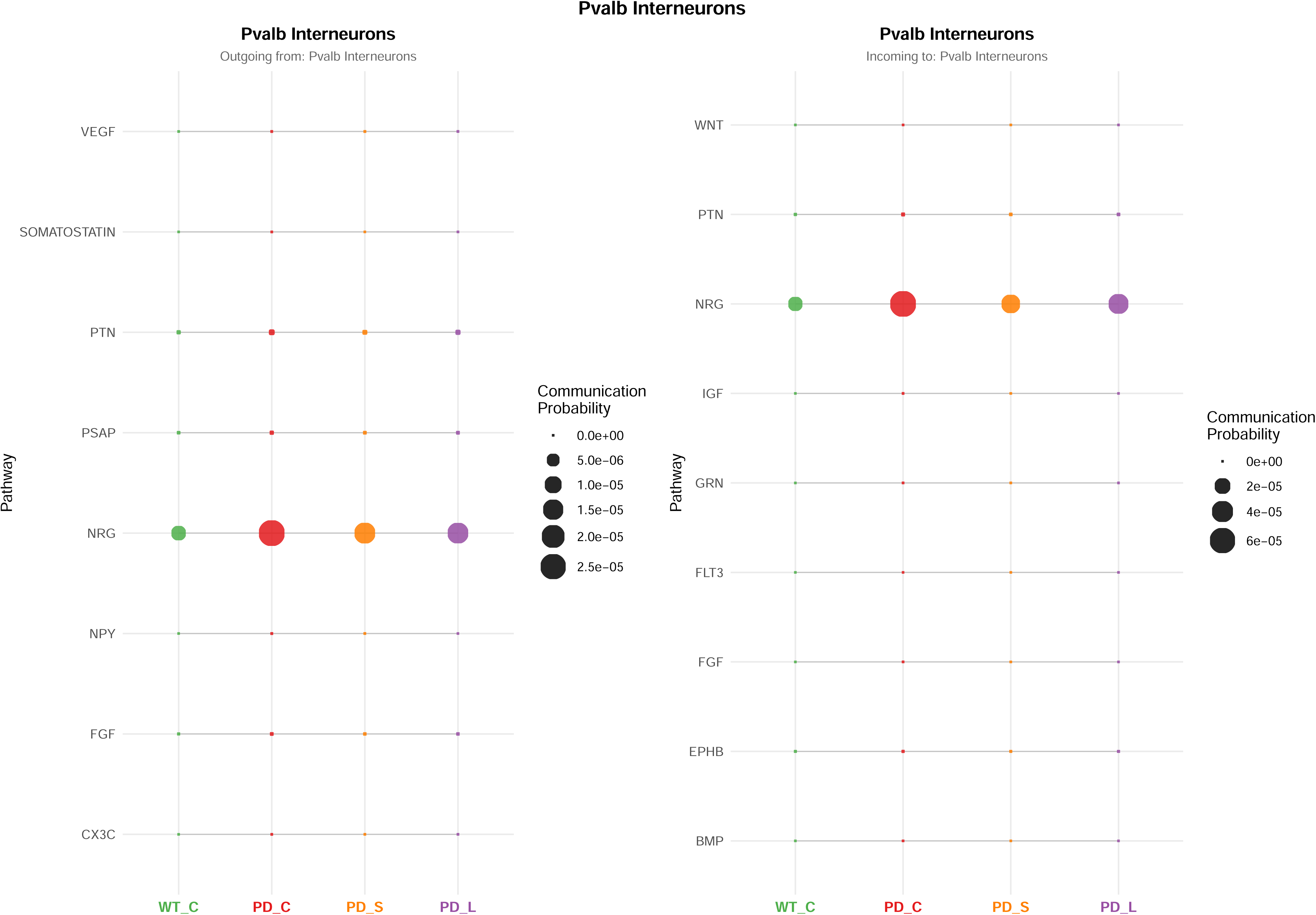

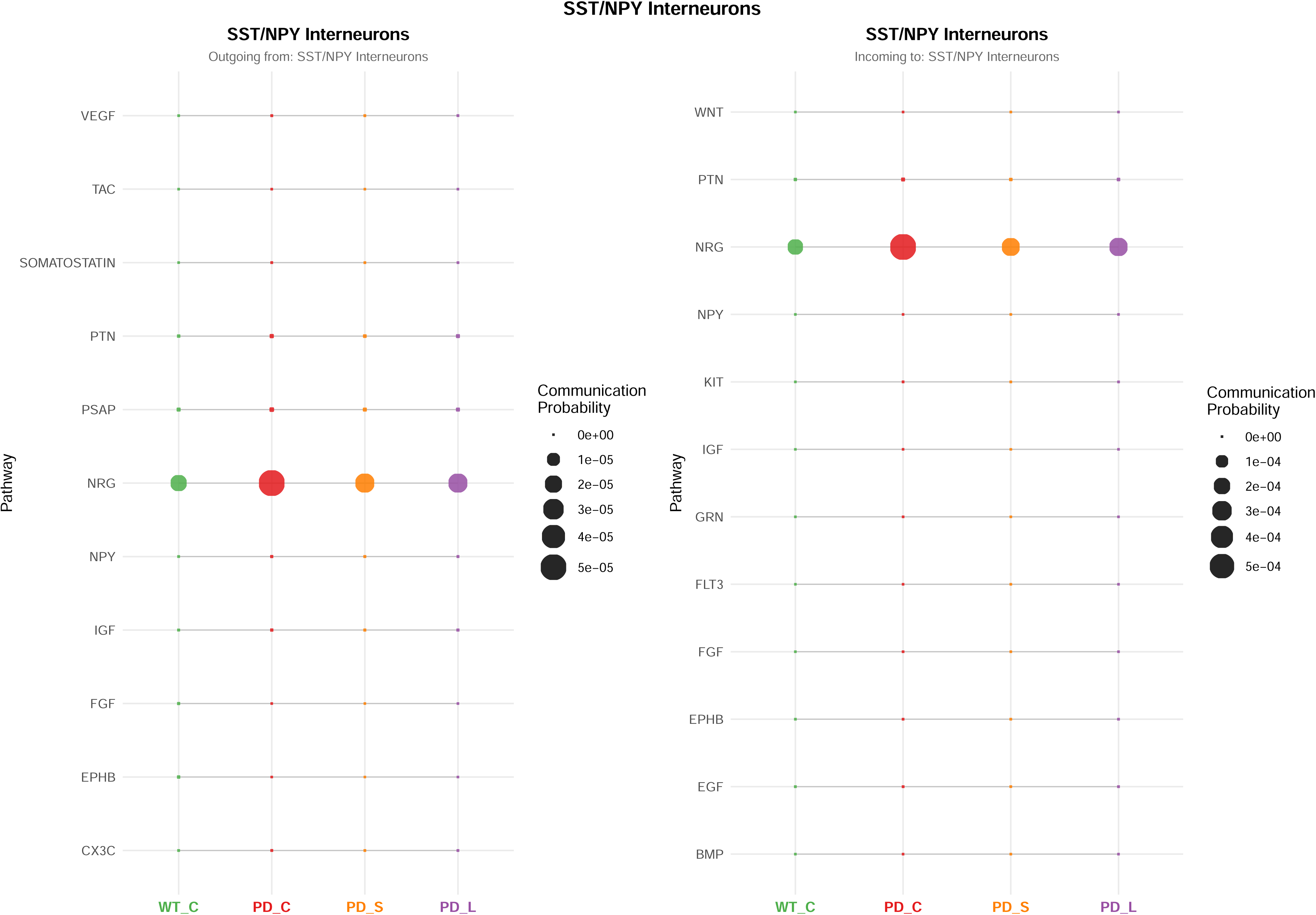

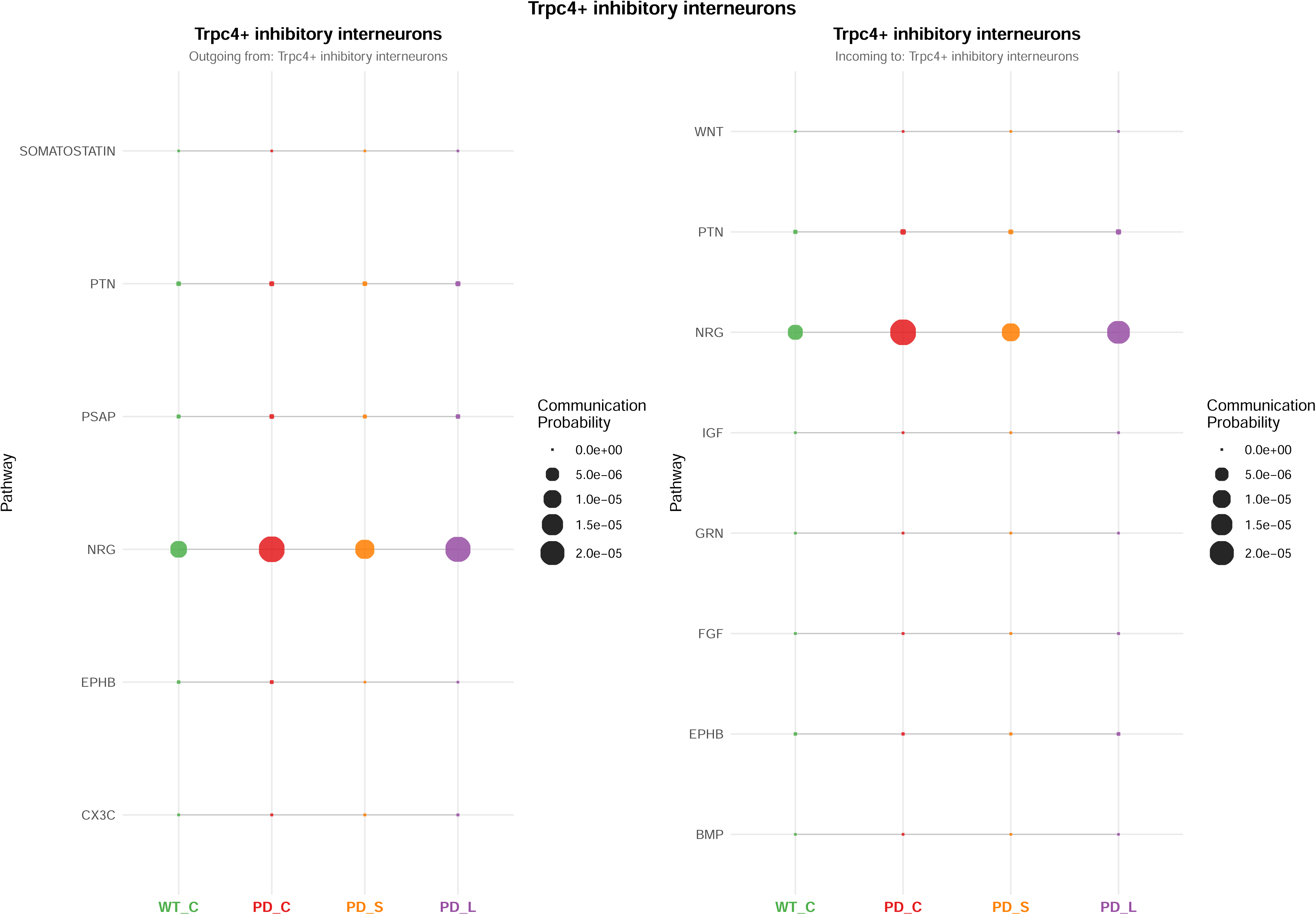

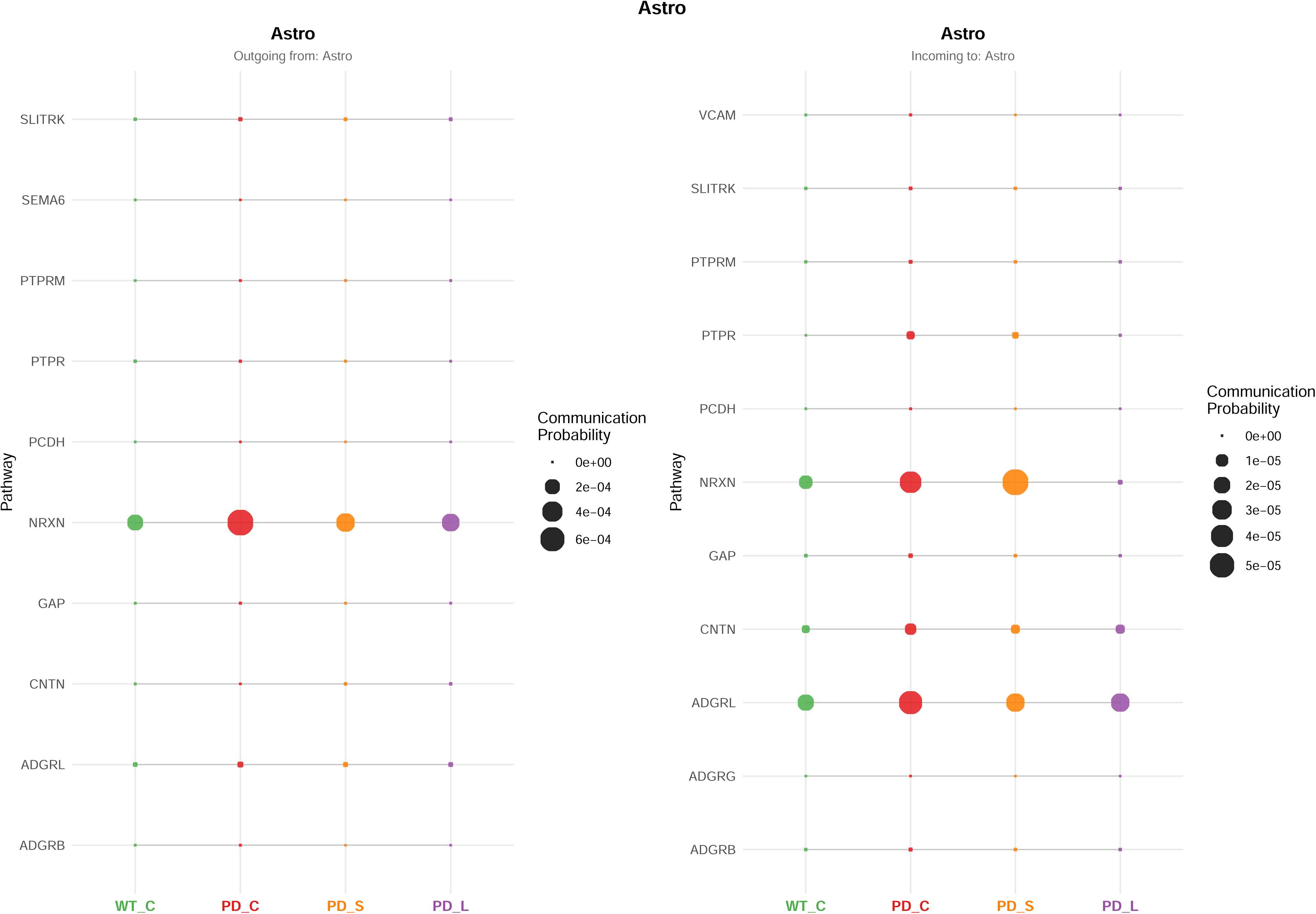

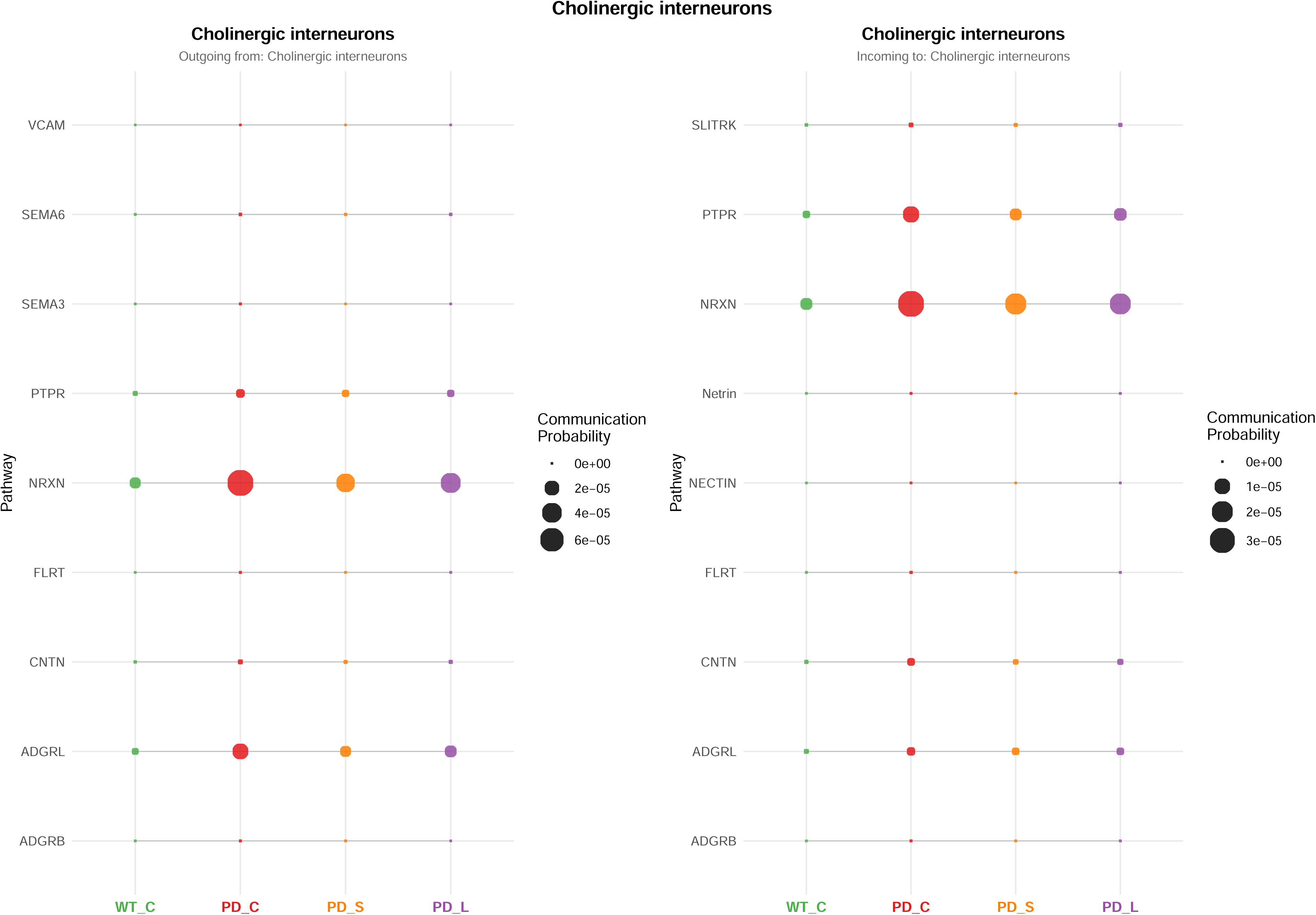

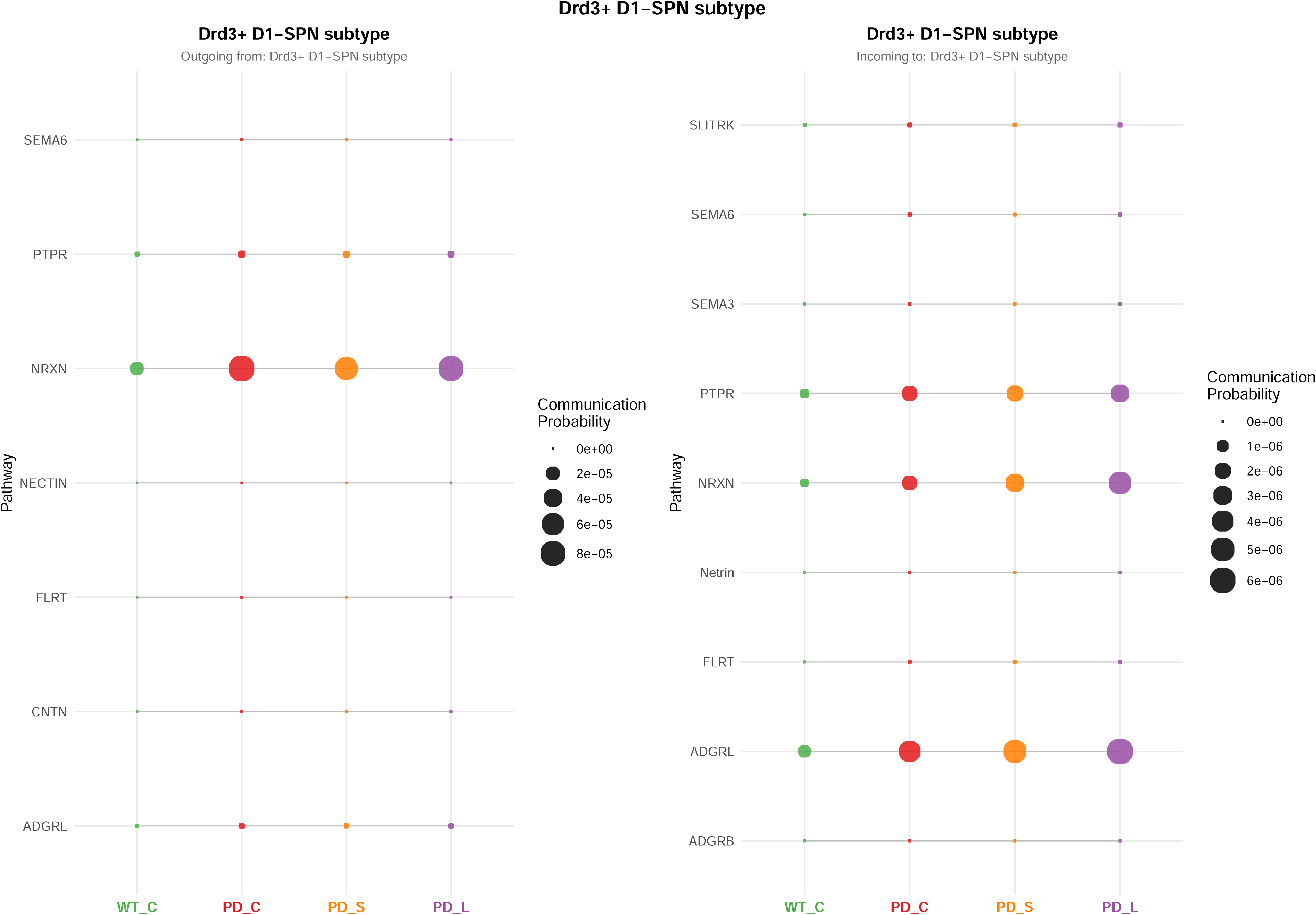

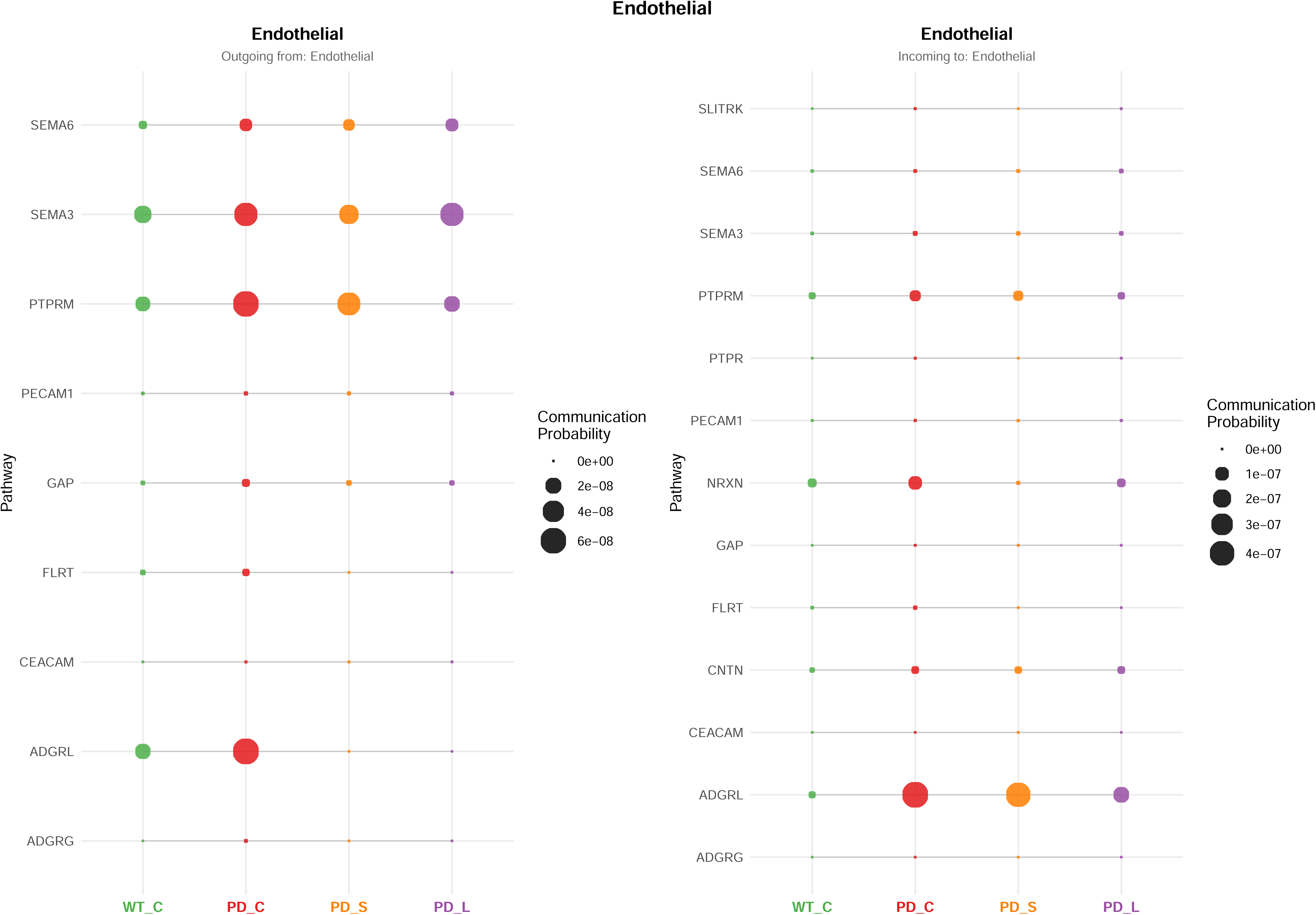

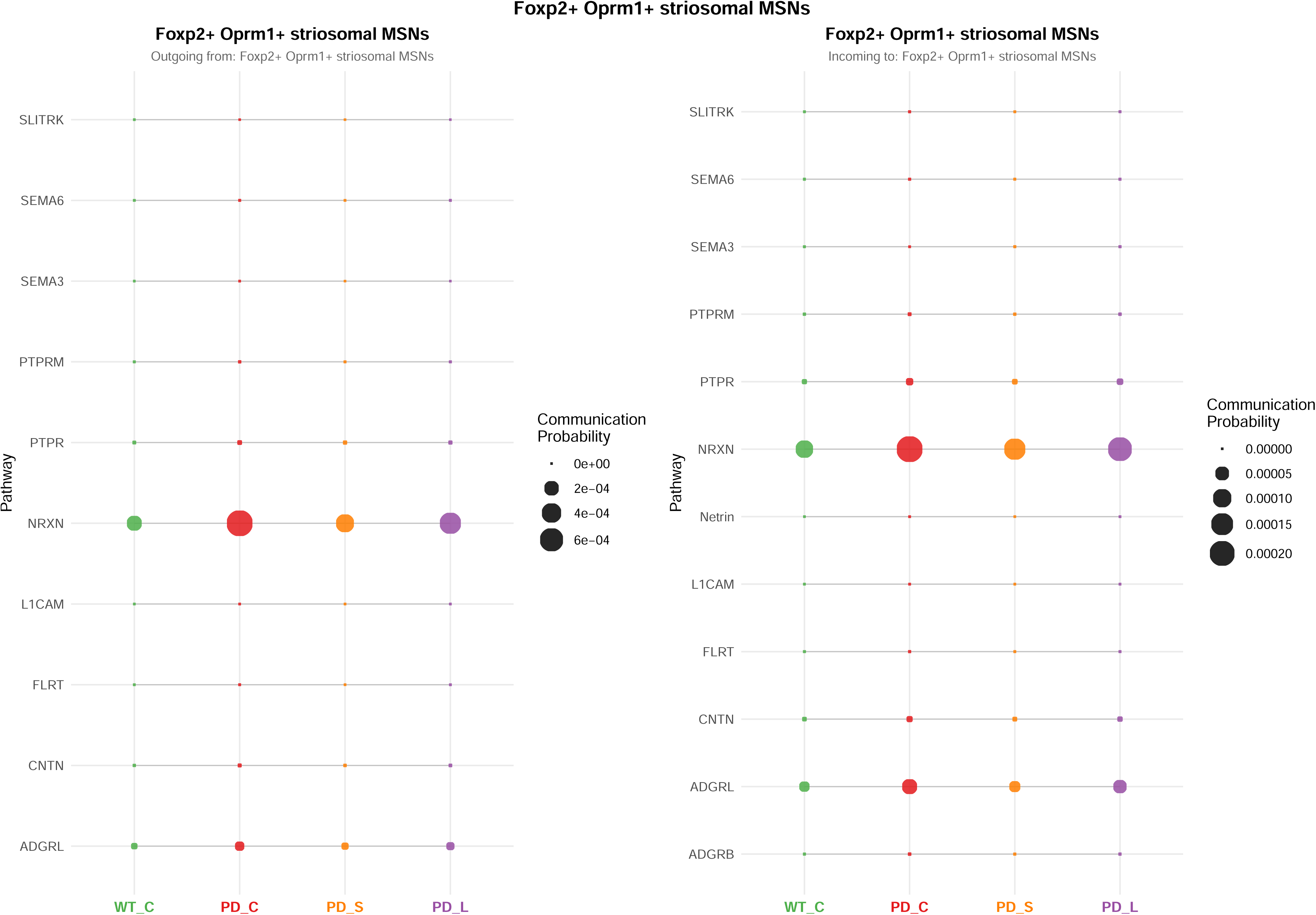

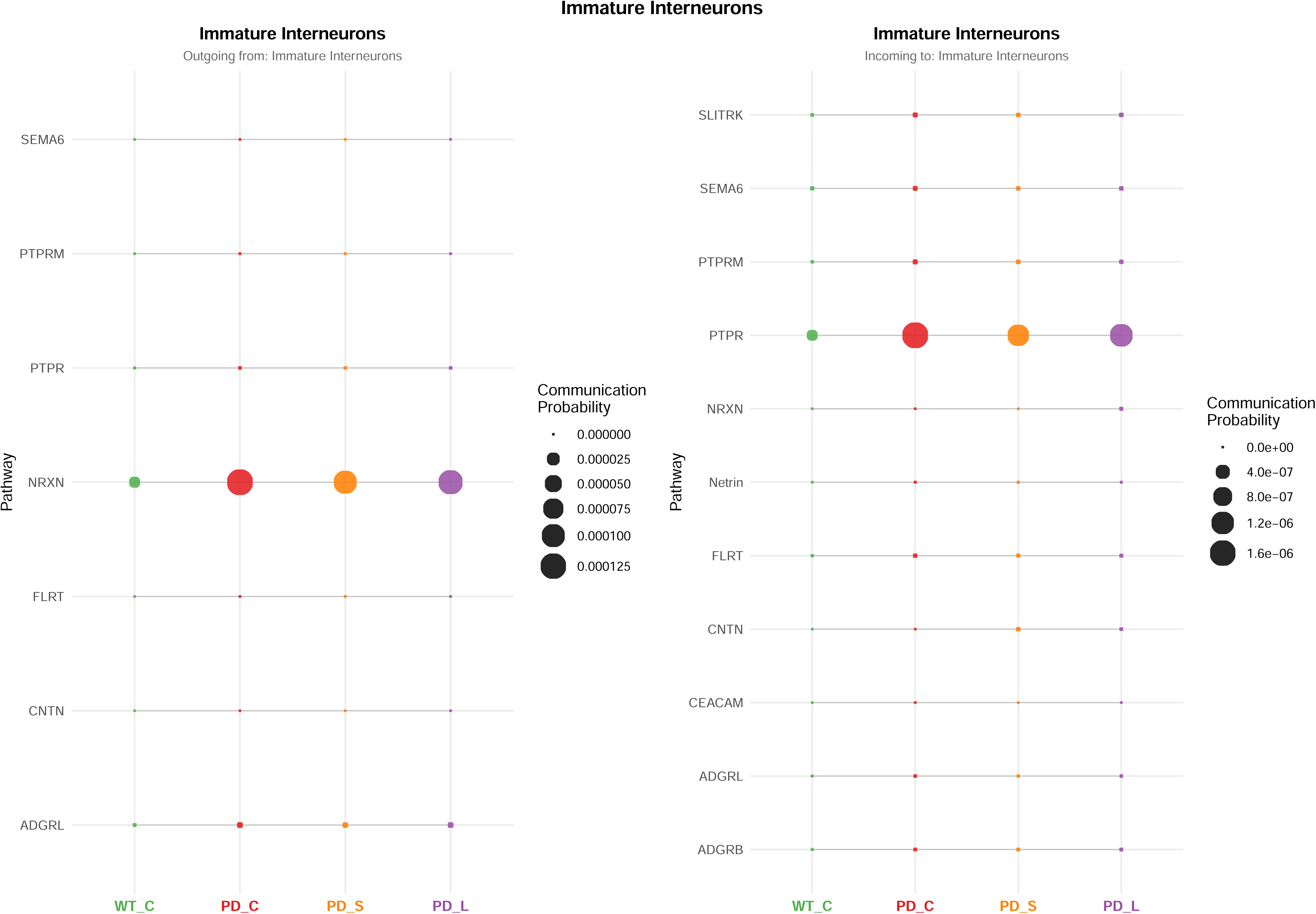

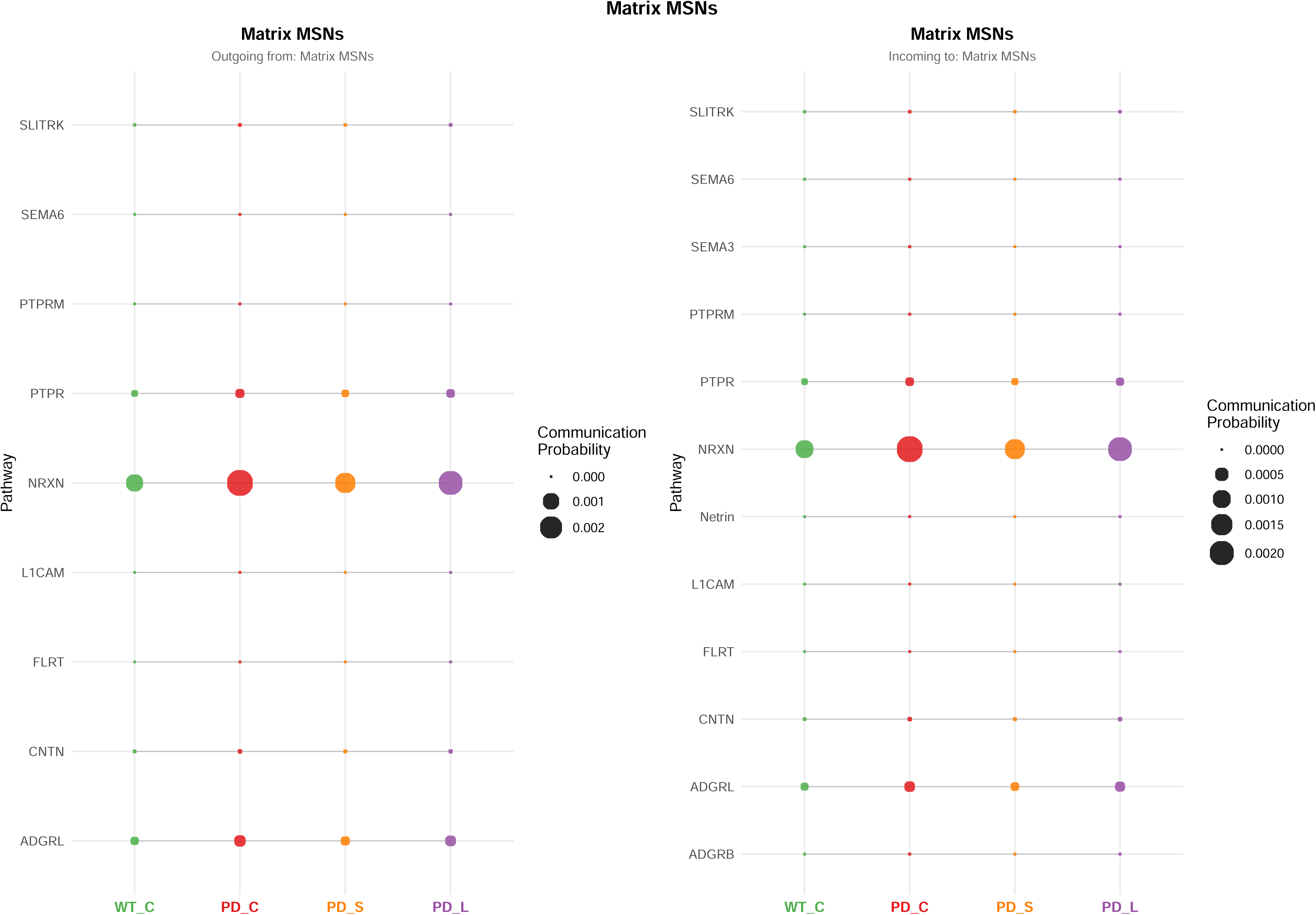

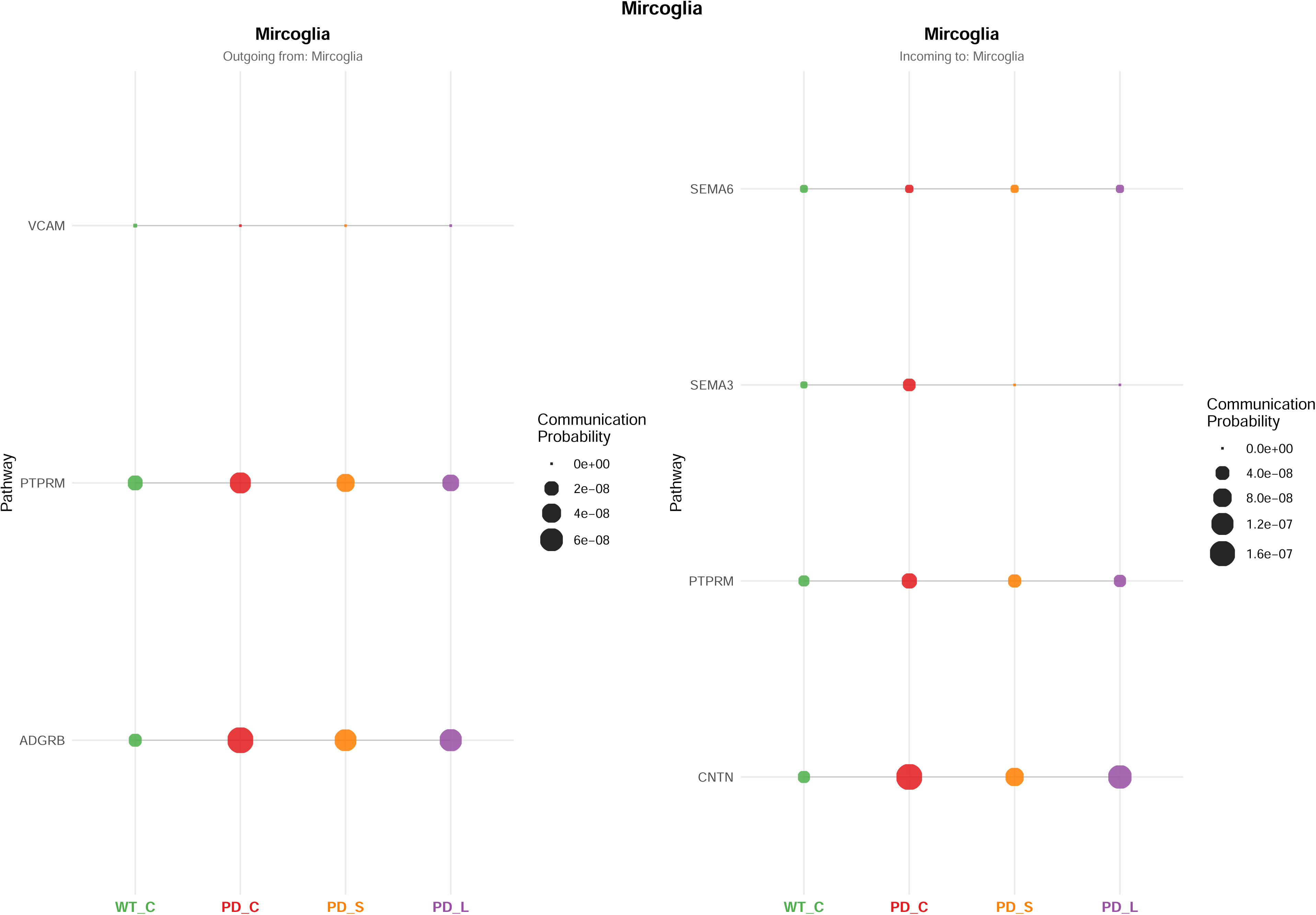

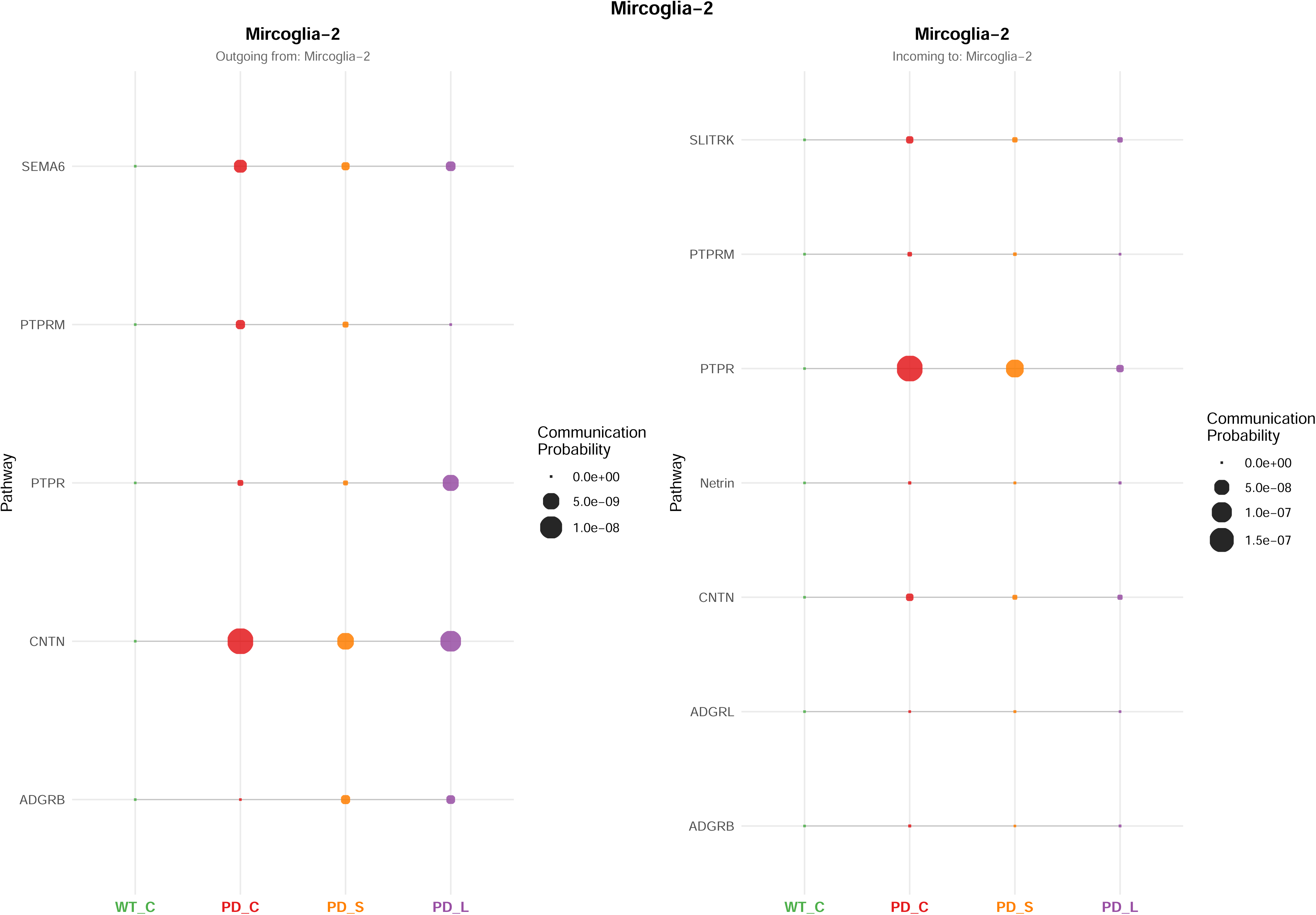

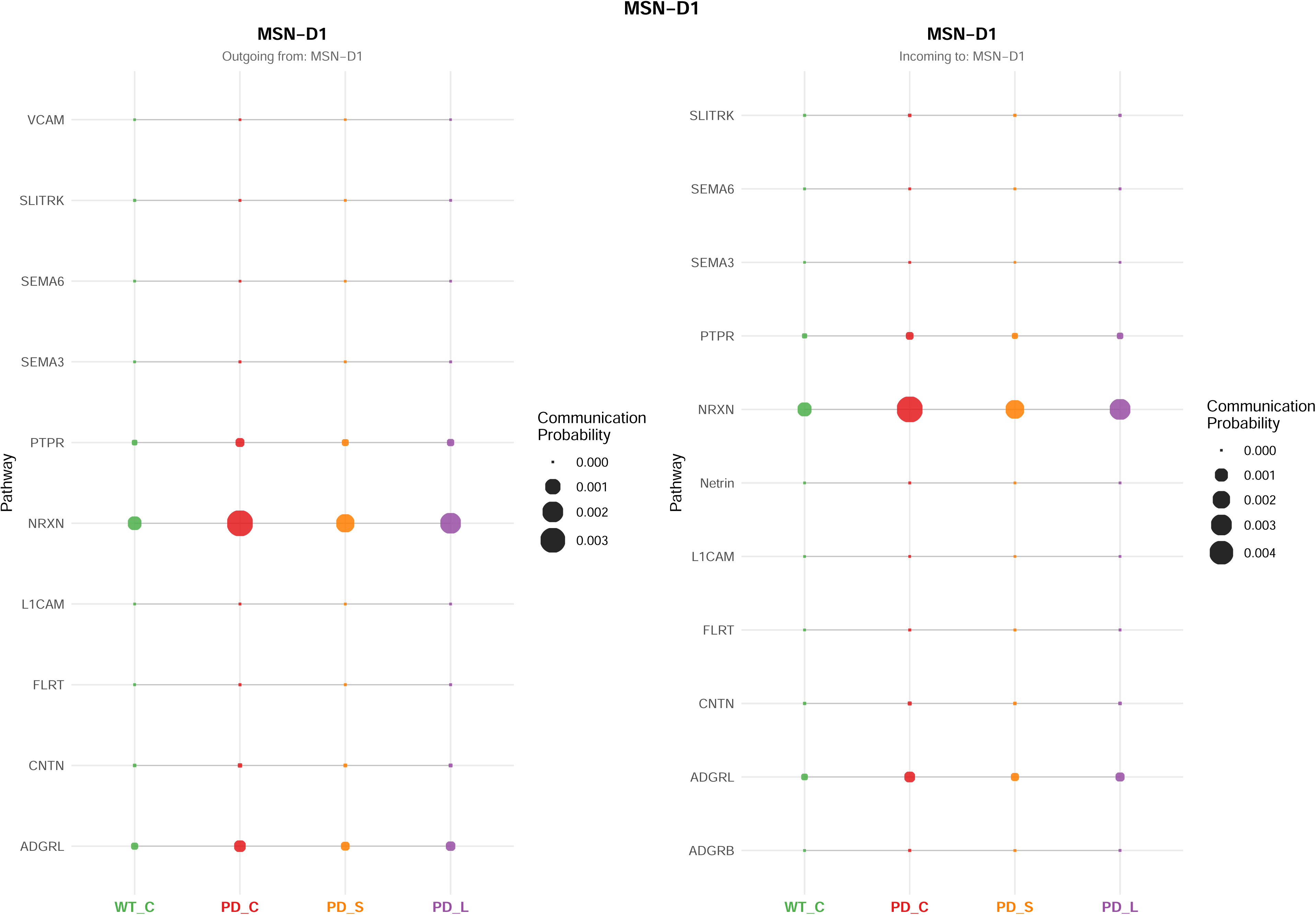

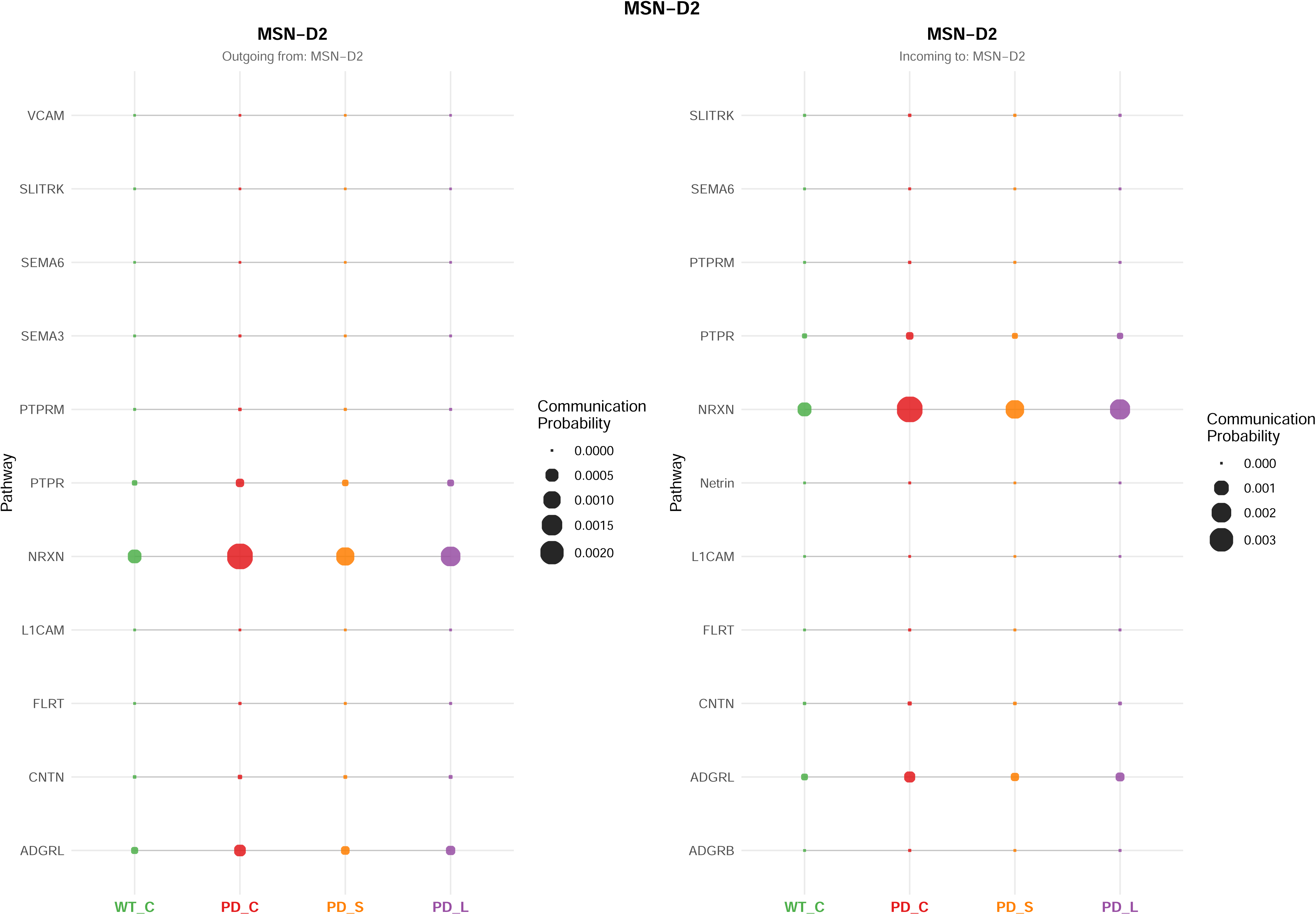

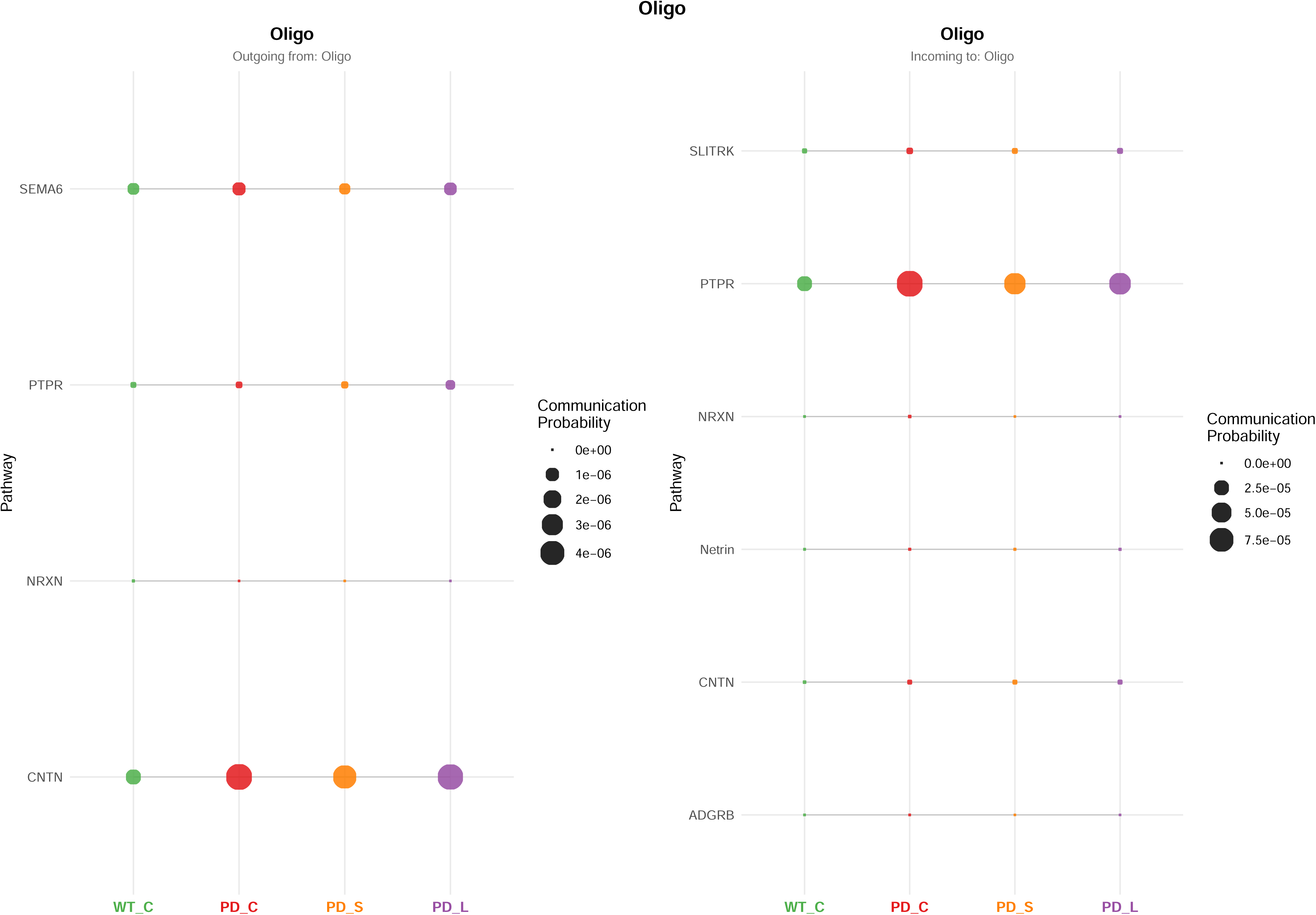

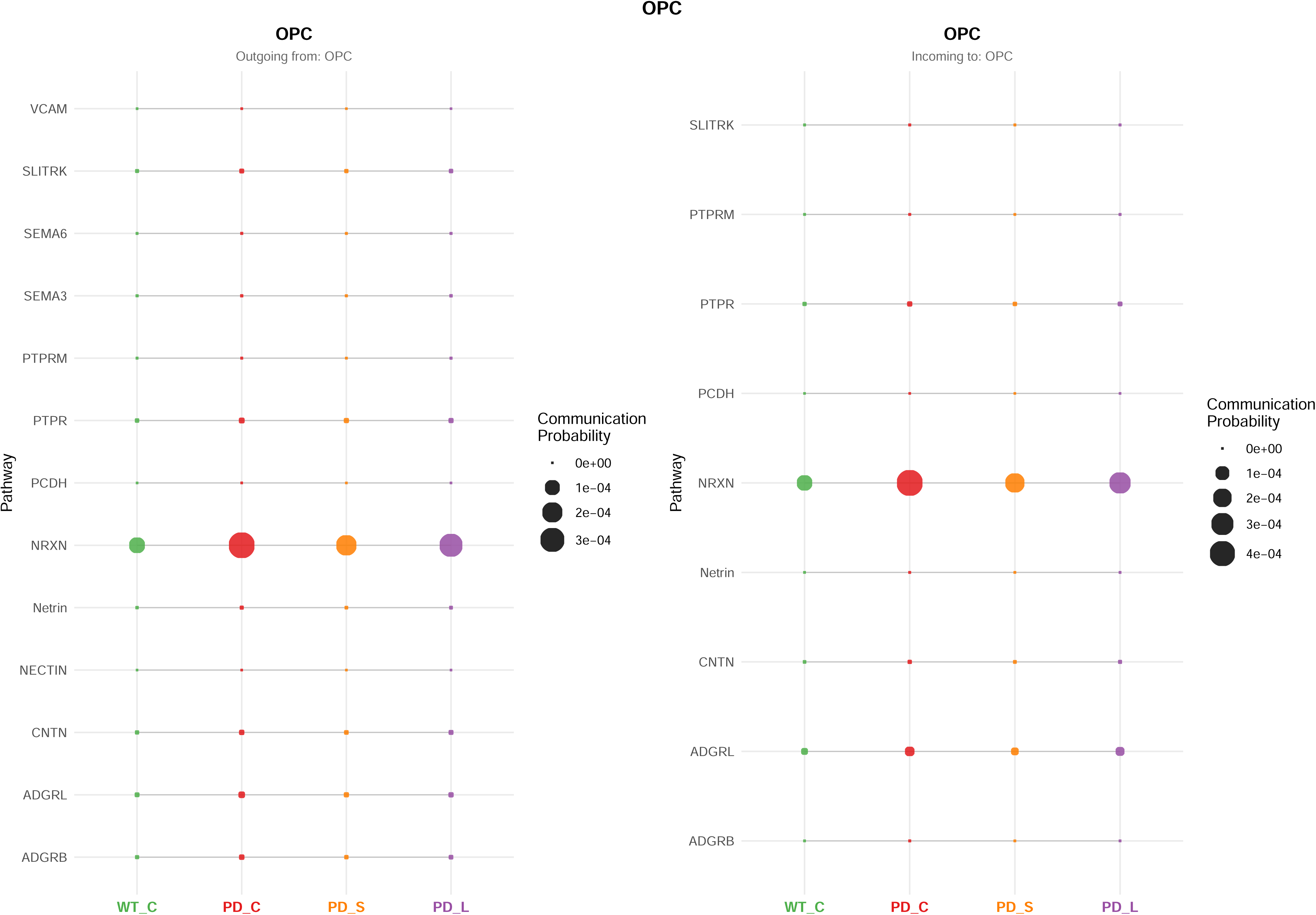

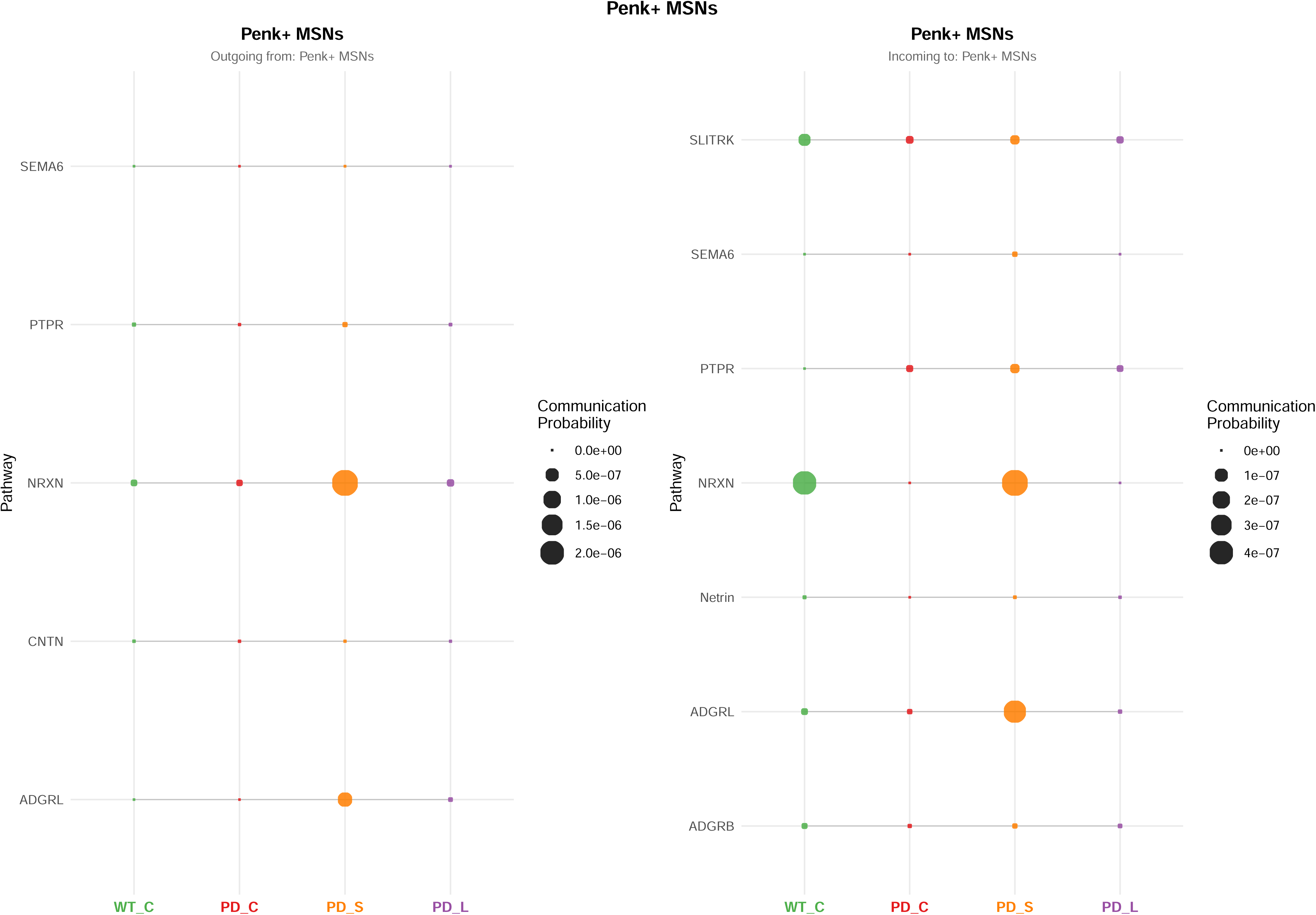

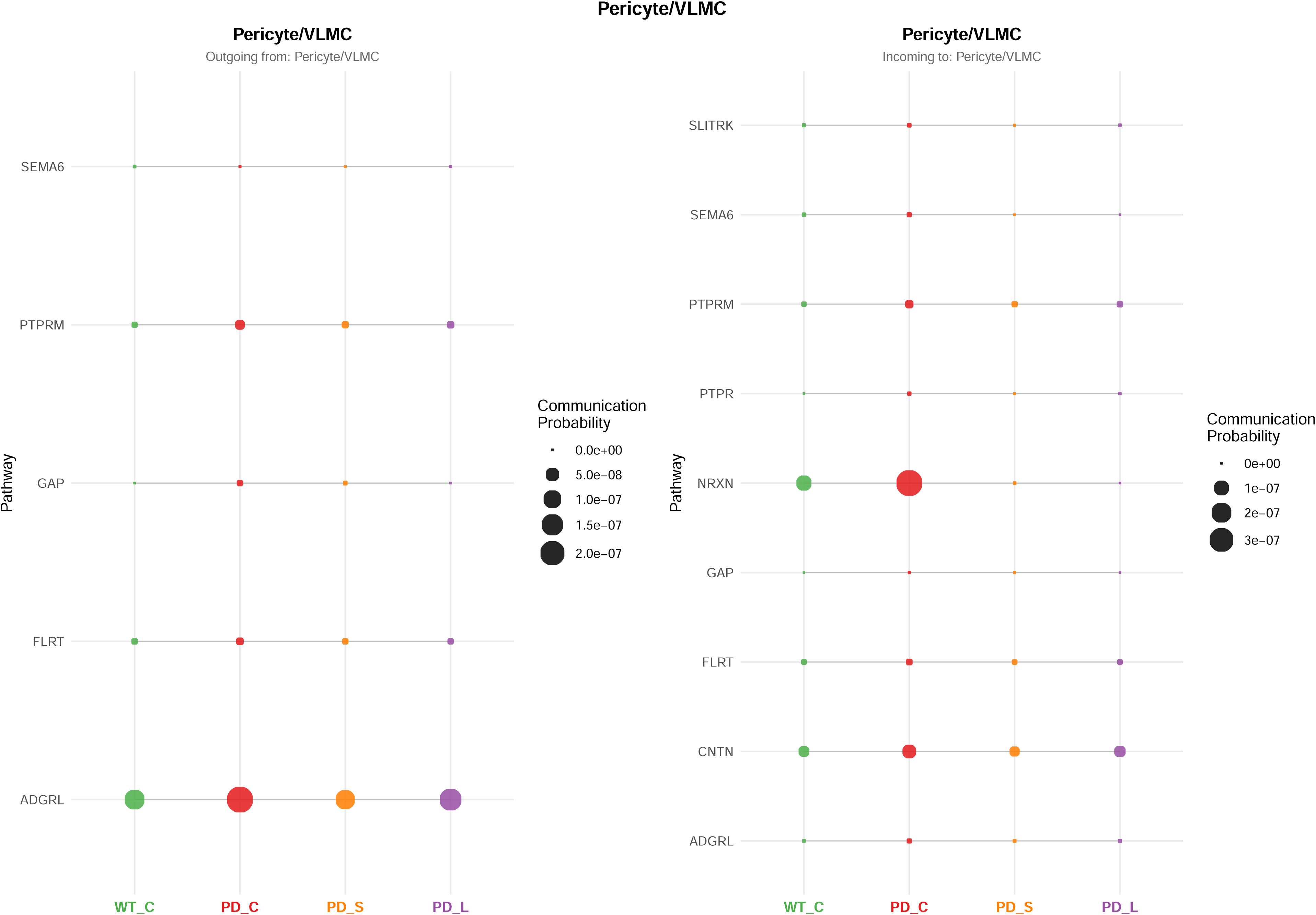

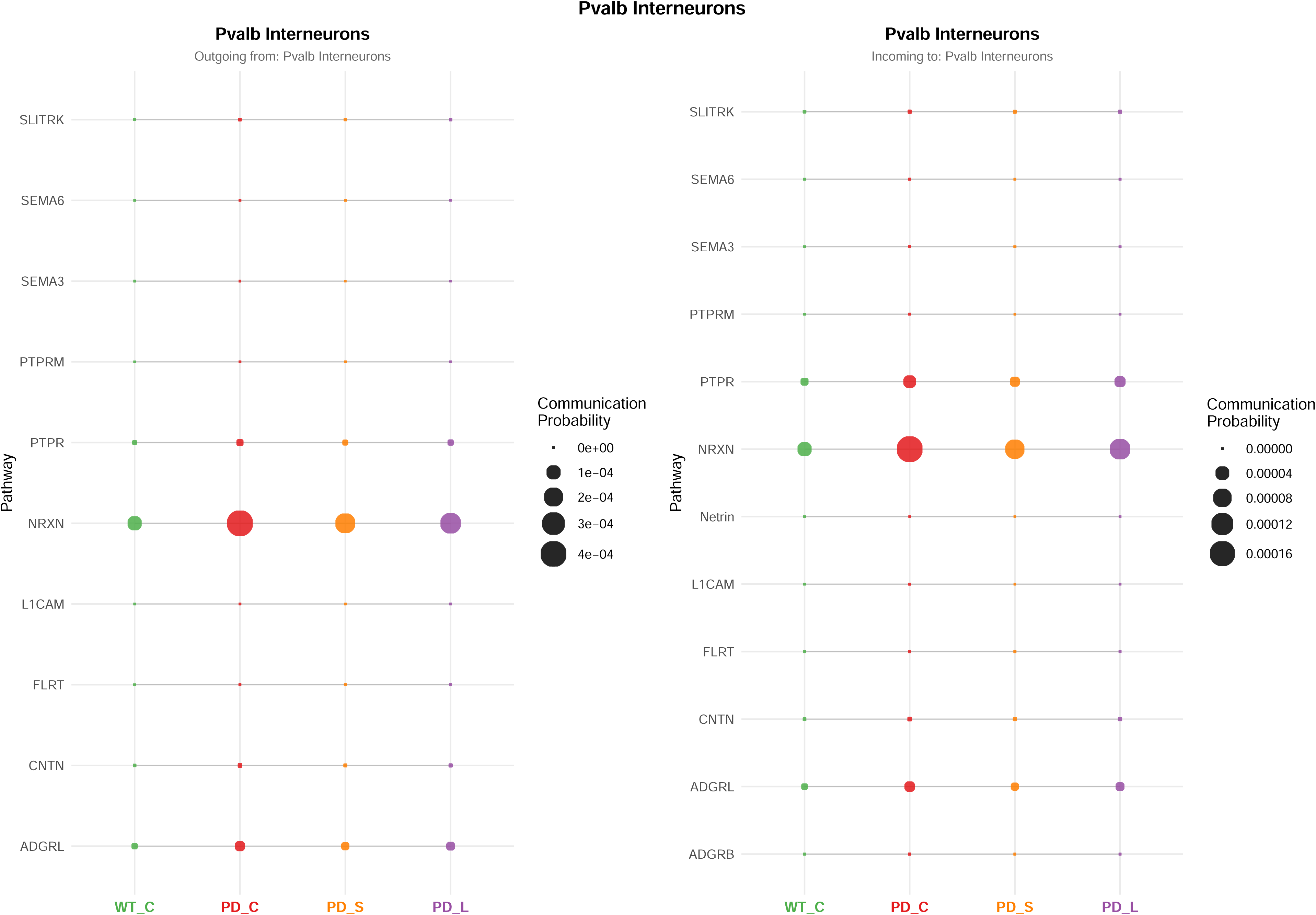

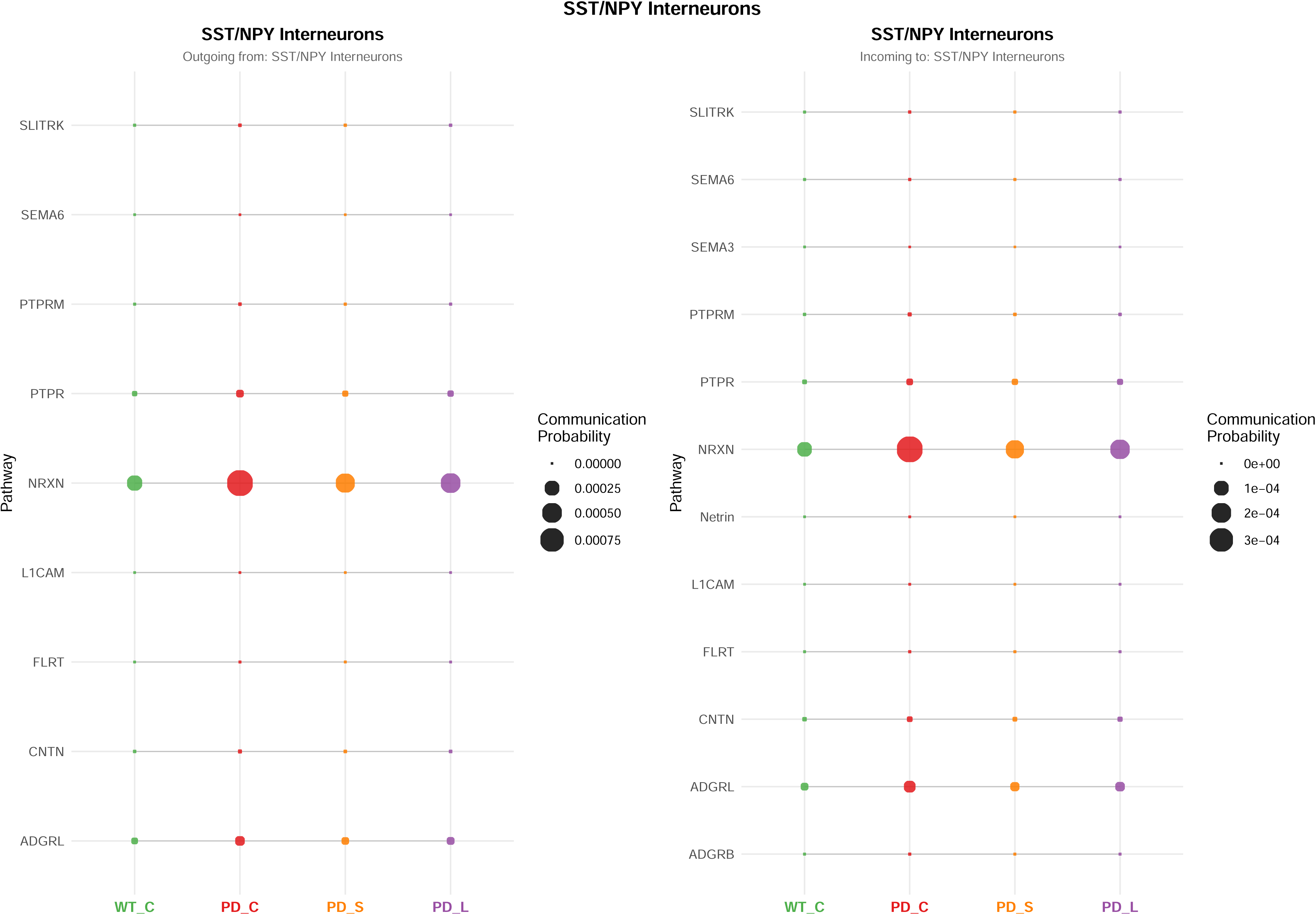

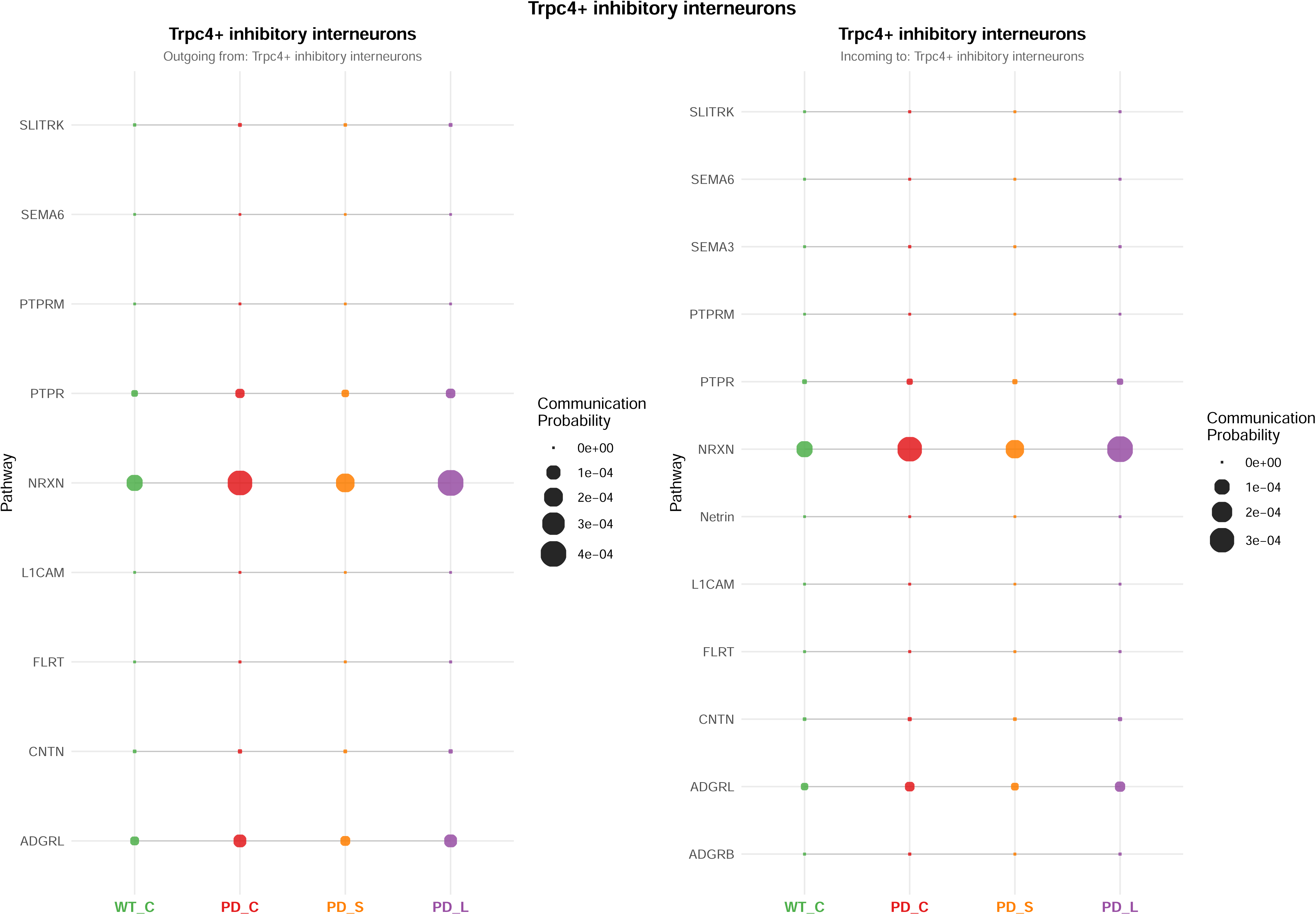

## References

1. D. Su, Y. Cui, C. He, P. Yin, R. Bai, J. Zhu, J. S. T. Lam, J. Zhang, R. Yan, X. Zheng, J. Wu, D. Zhao, A. Wang, M. Zhou, T. Feng, Projections for prevalence of Parkinson’s disease and its driving factors in 195 countries and territories to 2050: modelling study of Global Burden of Disease Study 2021. BMJ (Clinical research ed.) 388, e080952 (2025).

2. G. W. Ross, H. Petrovitch, R. D. Abbott, C. M. Tanner, J. Popper, K. Masaki, L. Launer, L. R. White, Association of olfactory dysfunction with risk for future Parkinson’s disease. Annals of neurology 63, 167–173 (2008).

3. D. Berg, R. B. Postuma, C. H. Adler, B. R. Bloem, P. Chan, B. Dubois, T. Gasser, C. G. Goetz, G. Halliday, L. Joseph, A. E. Lang, I. Liepelt-Scarfone, I. Litvan, K. Marek, J. Obeso, W. Oertel, C. W. Olanow, W. Poewe, M. Stern, G. Deuschl, MDS research criteria for prodromal Parkinson’s disease. Movement disorders : official journal of the Movement Disorder Society 30, 1600–1611 (2015).

4. W. Poewe, K. Seppi, C. M. Tanner, G. M. Halliday, P. Brundin, J. Volkmann, A. E. Schrag, A. E. Lang, Parkinson disease. Nature reviews. Disease primers 3, 17013 (2017).

5. W. G. Meissner, P. Remy, C. Giordana, D. Maltete, P. Derkinderen, J. L. Houeto, M. Anheim, I. Benatru, T. Boraud, C. Brefel-Courbon, N. Carriere, H. Catala, O. Colin, J. C. Corvol, P. Damier, E. Dellapina, D. Devos, S. Drapier, M. Fabbri, V. Ferrier, A. Foubert-Samier, S. Frismand-Kryloff, A. Georget, C. Germain, S. Grimaldi, C. Hardy, L. Hopes, P. Krystkowiak, B. Laurens, R. Lefaucheur, L. L. Mariani, A. Marques, C. Marse, F. Ory-Magne, V. Rigalleau, H. Salhi, A. Saubion, S. R. W. Stott, C. Thalamas, C. Thiriez, M. Tir, R. K. Wyse, A. Benard, O. Rascol, L. S. Group, Trial of Lixisenatide in Early Parkinson’s Disease. The New England journal of medicine 390, 1176–1185 (2024).

6. J. M. Friedman, The discovery and development of GLP-1 based drugs that have revolutionized the treatment of obesity. Proceedings of the National Academy of Sciences of the United States of America 121, e2415550121 (2024).

7. L. Zhang, L. Zhang, L. Li, C. Holscher, Semaglutide is Neuroprotective and Reduces alpha-Synuclein Levels in the Chronic MPTP Mouse Model of Parkinson’s Disease. Journal of Parkinson’s disease 9, 157–171 (2019).

8. J. Jalewa, M. K. Sharma, S. Gengler, C. Hölscher, A novel GLP-1/GIP dual receptor agonist protects from 6-OHDA lesion in a rat model of Parkinson’s disease. Neuropharmacology 117, 238–248 (2017).

9. N. Vijiaratnam, C. Girges, G. Auld, R. McComish, A. King, S. S. Skene, S. Hibbert, A. Wong, S. Melander, R. Gibson, H. Matthews, J. Dickson, C. Carroll, A. Patrick, J. Inches, M. Silverdale, B. Blackledge, J. Whiston, M. Hu, J. Welch, G. Duncan, K. Power, S. Gallen, J. Kerr, K. R. Chaudhuri, L. Batzu, S. Rota, E. Jabbari, H. Morris, P. Limousin, N. Greig, Y. Li, V. Libri, S. Gandhi, D. Athauda, K. Chowdhury, T. Foltynie, Exenatide once a week versus placebo as a potential disease-modifying treatment for people with Parkinson’s disease in the UK: a phase 3, multicentre, double-blind, parallel-group, randomised, placebo-controlled trial. Lancet (London, England) 405, 627–636 (2025).

10. Y. Kimura, T. Koda, H. Kurakami, S. Sakamoto, K. Iwasaki, K. Asai, L. Ge, H. Kato, T. Tsuboi, N. Matsukawa, O. Kano, D. Matsuse, M. Tomiyama, M. Yokoe, Y. Nagai, H. Mochizuki, Disease-modifying effect, safety and optimal dose of oral semaglutide tablets for patients with Parkinson’s disease (MOST-ABLE study): protocol for a randomised, double-blind, placebo-controlled study. BMJ open 15, e112318 (2025).

11. K. Brockmann, C. Quadalti, S. Lerche, M. Rossi, I. Wurster, S. Baiardi, B. Roeben, A. Mammana, M. Zimmermann, A. K. Hauser, C. Deuschle, C. Schulte, K. Waniek, I. Lachmann, S. Sjödin, A. Brinkmalm, K. Blennow, H. Zetterberg, T. Gasser, P. Parchi, Association between CSF alpha-synuclein seeding activity and genetic status in Parkinson’s disease and dementia with Lewy bodies. Acta neuropathologica communications 9, 175 (2021).

12. Y. Kim, J. McInnes, J. Kim, Y. H. W. Liang, S. Veeraragavan, A. R. Garza, B. D. W. Belfort, B. Arenkiel, R. Samaco, H. Y. Zoghbi, Olfactory deficit and gastrointestinal dysfunction precede motor abnormalities in alpha-Synuclein G51D knock-in mice. Proceedings of the National Academy of Sciences of the United States of America 121, e2406479121 (2024).

13. A. Siderowf, D. Jennings, S. Eberly, D. Oakes, K. A. Hawkins, A. Ascherio, M. B. Stern, K. Marek, P. Investigators, Impaired olfaction and other prodromal features in the Parkinson At-Risk Syndrome Study. Movement disorders : official journal of the Movement Disorder Society 27, 406–412 (2012).

14. K. L. Adams-Carr, J. P. Bestwick, S. Shribman, A. Lees, A. Schrag, A. J. Noyce, Constipation preceding Parkinson’s disease: a systematic review and meta-analysis. Journal of neurology, neurosurgery, and psychiatry 87, 710–716 (2016).

15. J. Lu, H. Liu, Q. Zhou, M. W. Wang, Z. Li, A potentially serious adverse effect of GLP-1 receptor agonists. Acta pharmaceutica Sinica. B 13, 2291–2293 (2023).

16. T. Simuni, C. Caspell-Garcia, C. S. Coffey, D. Weintraub, B. Mollenhauer, S. Lasch, C. M. Tanner, D. Jennings, K. Kieburtz, L. M. Chahine, K. Marek, Baseline prevalence and longitudinal evolution of non-motor symptoms in early Parkinson’s disease: the PPMI cohort. Journal of neurology, neurosurgery, and psychiatry 89, 78–88 (2018).

17. I. Stankovic, I. Petrovic, T. Pekmezovic, V. Markovic, T. Stojkovic, N. Dragasevic-Miskovic, M. Svetel, V. Kostic, Longitudinal assessment of autonomic dysfunction in early Parkinson’s disease. Parkinsonism & related disorders 66, 74–79 (2019).

18. L. A. Martins, A. Schiavo, L. L. Xavier, R. G. Mestriner, The Foot Fault Scoring System to Assess Skilled Walking in Rodents: A Reliability Study. Frontiers in behavioral neuroscience 16, 892010 (2022).

19. H. Shiotsuki, K. Yoshimi, Y. Shimo, M. Funayama, Y. Takamatsu, K. Ikeda, R. Takahashi, S. Kitazawa, N. Hattori, A rotarod test for evaluation of motor skill learning. Journal of neuroscience methods 189, 180–185 (2010).

20. B. Vaidya, S. Biswas, I. Roy, S. S. Sharma, HC070, a Transient Receptor Potential Canonical 5 (TRPC5) Channels Inhibitor Ameliorated alpha-synuclein Preformed Fibrils-Induced Parkinson’s Disease: A Neurobehavioural and Mechanistic Study. Journal of biochemical and molecular toxicology 39, e70207 (2025).

21. B. Araujo, R. Caridade-Silva, C. Soares-Guedes, J. Martins-Macedo, E. D. Gomes, S. Monteiro, F. G. Teixeira, Neuroinflammation and Parkinson’s Disease-From Neurodegeneration to Therapeutic Opportunities. Cells 11, (2022).

22. A. S. Kulkarni, M. Del Mar Cortijo, E. R. Roberts, T. L. Suggs, H. B. Stover, J. I. Pena-Bravo, J. A. Steiner, K. C. Luk, P. Brundin, D. W. Wesson, Perturbation of in vivo Neural Activity Following alpha-Synuclein Seeding in the Olfactory Bulb. Journal of Parkinson’s disease 10, 1411–1427 (2020).

23. N. L. Rey, S. George, J. A. Steiner, Z. Madaj, K. C. Luk, J. Q. Trojanowski, V. M. Lee, P. Brundin, Spread of aggregates after olfactory bulb injection of alpha-synuclein fibrils is associated with early neuronal loss and is reduced long term. Acta neuropathologica 135, 65–83 (2018).

24. H. Bernheimer, W. Birkmayer, O. Hornykiewicz, K. Jellinger, F. Seitelberger, Brain dopamine and the syndromes of Parkinson and Huntington. Clinical, morphological and neurochemical correlations. Journal of the neurological sciences 20, 415–455 (1973).

25. C. M. Lim, M. Vendruscolo, The alpha-synuclein proteostasis network and its translational applications in Parkinson’s disease. Proceedings of the National Academy of Sciences of the United States of America 123, e2513317123 (2026).

26. S. Lesage, M. Anheim, F. Letournel, L. Bousset, A. Honore, N. Rozas, L. Pieri, K. Madiona, A. Durr, R. Melki, C. Verny, A. Brice,G. French Parkinson’s Disease Genetics Study, G51D alpha-synuclein mutation causes a novel parkinsonian-pyramidal syndrome. Annals of neurology 73, 459–471 (2013).

27. M. H. Patton, B. J. W. Teubner, K. T. Thomas, S. L. Freshour, A. J. Trevisan, A. B. Schild, C. A. Ramirez, J. B. Bikoff, S. S. Zakharenko, Schizophrenia-associated 22q11.2 deletion elevates striatal acetylcholine and disrupts thalamostriatal projections to produce amotivation in mice. bioRxiv : the preprint server for biology, (2025).

28. S. Jin, C. F. Guerrero-Juarez, L. Zhang, I. Chang, R. Ramos, C. H. Kuan, P. Myung, M. V. Plikus, Q. Nie, Inference and analysis of cell-cell communication using CellChat. Nature communications 12, 1088 (2021).

29. E. Armingol, A. Officer, O. Harismendy, N. E. Lewis, Deciphering cell-cell interactions and communication from gene expression. Nature reviews. Genetics 22, 71–88 (2021).

30. M. Huang, L. Xu, J. Liu, P. Huang, Y. Tan, S. Chen, Cell-Cell Communication Alterations via Intercellular Signaling Pathways in Substantia Nigra of Parkinson’s Disease. Frontiers in aging neuroscience 14, 828457 (2022).

31. W. Dauer, S. Przedborski, Parkinson’s disease: mechanisms and models. Neuron 39, 889–909 (2003).

32. D. Dominguez-Paredes, K. Kozielski, M. Ries, Y. Temel, A. Jahanshahi, Phenotypical variability in the 6-OHDA mouse model of Parkinson’s disease despite consistent and robust nigral lesioning. J Brain Mechanisms, 202523 (2025).

33. S. P. Yun, T. I. Kam, N. Panicker, S. Kim, Y. Oh, J. S. Park, S. H. Kwon, Y. J. Park, S. S. Karuppagounder, H. Park, S. Kim, N. Oh, N. A. Kim, S. Lee, S. Brahmachari, X. Mao, J. H. Lee, M. Kumar, D. An, S. U. Kang, Y. Lee, K. C. Lee, D. H. Na, D. Kim, S. H. Lee, V. V. Roschke, S. A. Liddelow, Z. Mari, B. A. Barres, V. L. Dawson, S. Lee, T. M. Dawson, H. S. Ko, Block of A1 astrocyte conversion by microglia is neuroprotective in models of Parkinson’s disease. Nature medicine 24, 931–938 (2018).

34. C. H. Gibbons, T. Levine, C. Adler, B. Bellaire, N. Wang, J. Stohl, P. Agarwal, G. M. Aldridge, A. Barboi, V. G. H. Evidente, D. Galasko, M. D. Geschwind, A. Gonzalez-Duarte, R. Gil, M. Gudesblatt, S. H. Isaacson, H. Kaufmann, P. Khemani, R. Kumar, G. Lamotte, A. J. Liu, N. R. McFarland, M. Miglis, A. Reynolds, G. A. Sahagian, M. H. Saint-Hillaire, J. B. Schwartzbard, W. Singer, M. J. Soileau, S. Vernino, O. Yerstein, R. Freeman, Skin Biopsy Detection of Phosphorylated α-Synuclein in Patients With Synucleinopathies. Jama 331, 1298–1306 (2024).

35. R. D. Abbott, H. Petrovitch, L. R. White, K. H. Masaki, C. M. Tanner, J. D. Curb, A. Grandinetti, P. L. Blanchette, J. S. Popper, G. W. Ross, Frequency of bowel movements and the future risk of Parkinson’s disease. Neurology 57, 456–462 (2001).

36. A. Ismaiel, G. G. M. Scarlata, I. Boitos, D. C. Leucuta, S. L. Popa, N. Al Srouji, L. Abenavoli, D. L. Dumitrascu, Gastrointestinal adverse events associated with GLP-1 RA in non-diabetic patients with overweight or obesity: a systematic review and network meta-analysis. International journal of obesity (2005) 49, 1946–1957 (2025).

37. P. Gonzalez-Rodriguez, E. Zampese, K. A. Stout, J. N. Guzman, E. Ilijic, B. Yang, T. Tkatch, M. A. Stavarache, D. L. Wokosin, L. Gao, M. G. Kaplitt, J. Lopez-Barneo, P. T. Schumacker, D. J. Surmeier, Disruption of mitochondrial complex I induces progressive parkinsonism. Nature 599, 650–656 (2021).

38. P. M. Keeney, J. Xie, R. A. Capaldi, J. P. Bennett, Jr., Parkinson’s disease brain mitochondrial complex I has oxidatively damaged subunits and is functionally impaired and misassembled. The Journal of neuroscience : the official journal of the Society for Neuroscience 26, 5256–5264 (2006).

39. D. Narendra, J. E. Walker, R. Youle, Mitochondrial quality control mediated by PINK1 and Parkin: links to parkinsonism. Cold Spring Harbor perspectives in biology 4, (2012).

40. P. Seibler, J. Graziotto, H. Jeong, F. Simunovic, C. Klein, D. Krainc, Mitochondrial Parkin recruitment is impaired in neurons derived from mutant PINK1 induced pluripotent stem cells. The Journal of neuroscience : the official journal of the Society for Neuroscience 31, 5970–5976 (2011).

41. S. Du, Q. Long, Y. Zhou, J. Fu, H. Wu, L. Yang, Y. Xie, Y. Ding, M. Zhang, J. Guo, M. Wang, J. Lin, M. Hu, J. Zhang, D. Yao, W. Li, F. Bao, G. Xiang, Y. Wu, Y. Huang, H. Liang, R. Wang, H. Li, B. Chen, C. Li, J. Wang, J. Zhang, D. Qin, J. Sun, Y. Zhu, F. Sun, W. Wang, G. Lu, W. Y. Chan, H. Zhao, C. Liu, X. Liu, Transplantation of encapsulated mitochondria alleviates dysfunction in mitochondrial and Parkinson’s disease models. Cell, (2026).

42. M. Kobara, H. Toba, T. Nakata, A Glucagon-like Peptide 1 Analog Protects Mitochondria and Attenuates Hypoxia-Reoxygenation Injury in Cultured Cardiomyocytes. Journal of cardiovascular pharmacology 79, 568–576 (2022).

43. N. Nuamnaichati, S. Mangmool, N. Chattipakorn, W. Parichatikanond, Stimulation of GLP-1 Receptor Inhibits Methylglyoxal-Induced Mitochondrial Dysfunctions in H9c2 Cardiomyoblasts: Potential Role of Epac/PI3K/Akt Pathway. Frontiers in pharmacology 11, 805 (2020).

44. C. Luna-Marco, A. M. de Maranon, A. Hermo-Argibay, Y. Rodriguez-Hernandez, J. Hermenejildo, M. Fernandez-Reyes, N. Apostolova, J. Vila, E. Sola, C. Morillas, S. Rovira-Llopis, M. Rocha, V. M. Victor, Effects of GLP-1 receptor agonists on mitochondrial function, inflammatory markers and leukocyte-endothelium interactions in type 2 diabetes. Redox biology 66, 102849 (2023).

45. C. Cebrian, J. D. Loike, D. Sulzer, Neuroinflammation in Parkinson’s disease animal models: a cell stress response or a step in neurodegeneration? Current topics in behavioral neurosciences 22, 237–270 (2015).

46. H. M. Gao, F. Zhang, H. Zhou, W. Kam, B. Wilson, J. S. Hong, Neuroinflammation and alpha-synuclein dysfunction potentiate each other, driving chronic progression of neurodegeneration in a mouse model of Parkinson’s disease. Environmental health perspectives 119, 807–814 (2011).

47. R. Sharon, M. S. Goldberg, I. Bar-Josef, R. A. Betensky, J. Shen, D. J. Selkoe, alpha-Synuclein occurs in lipid-rich high molecular weight complexes, binds fatty acids, and shows homology to the fatty acid-binding proteins. Proceedings of the National Academy of Sciences of the United States of America 98, 9110–9115 (2001).

48. L. Merino-Galan, H. Jimenez-Urbieta, M. Zamarbide, T. Rodriguez-Chinchilla, A. Belloso-Iguerategui, E. Santamaria, J. Fernandez-Irigoyen, A. Aiastui, E. Doudnikoff, E. Bezard, A. Ouro, S. Knafo, B. Gago, A. Quiroga-Varela, M. C. Rodriguez-Oroz, Striatal synaptic bioenergetic and autophagic decline in premotor experimental parkinsonism. Brain : a journal of neurology 145, 2092–2107 (2022).

49. B. G. Brinkmann, A. Agarwal, M. W. Sereda, A. N. Garratt, T. Muller, H. Wende, R. M. Stassart, S. Nawaz, C. Humml, V. Velanac, K. Radyushkin, S. Goebbels, T. M. Fischer, R. J. Franklin, C. Lai, H. Ehrenreich, C. Birchmeier, M. H. Schwab, K. A. Nave, Neuregulin-1/ErbB signaling serves distinct functions in myelination of the peripheral and central nervous system. Neuron 59, 581–595 (2008).

50. T. C. Sudhof, Synaptic Neurexin Complexes: A Molecular Code for the Logic of Neural Circuits. Cell 171, 745–769 (2017).

51. K. Cuttler, D. de Swardt, L. Engelbrecht, J. Kriel, R. Cloete, S. Bardien, Neurexin 2 p.G849D variant, implicated in Parkinson’s disease, increases reactive oxygen species, and reduces cell viability and mitochondrial membrane potential in SH-SY5Y cells. Journal of neural transmission (Vienna, Austria : 1996) 129, 1435–1446 (2022).

52. S. Zhang, C. Zhang, Y. Zhang, Y. Feng, Unraveling the role of neuregulin-mediated astrocytes-OPCs axis in the pathogenesis of age-related macular degeneration and Parkinson’s disease. Scientific reports 15, 7352 (2025).

53. G. Ennequin, F. Capel, K. Caillaud, V. Chavanelle, M. Etienne, A. Teixeira, X. Li, N. Boisseau, P. Sirvent, Neuregulin 1 improves complex 2-mediated mitochondrial respiration in skeletal muscle of healthy and diabetic mice. Scientific reports 7, 1742 (2017).

54. F. Diaz-Saez, C. Balcells, L. Rossello, I. Lopez-Soldado, M. Romero, D. Sebastian, F. J. Lopez-Soriano, S. Busquets, M. Cascante, W. Ricart, J. M. Fernandez-Real, J. M. Moreno-Navarrete, J. Aragones, X. Testar, M. Camps, A. Zorzano, A. Guma, Neuregulin 4 Downregulation Alters Mitochondrial Morphology and Induces Oxidative Stress in 3T3-L1 Adipocytes. International journal of molecular sciences 25, (2024).

55. A. Guma, F. Diaz-Saez, M. Camps, A. Zorzano, Neuregulin, an Effector on Mitochondria Metabolism That Preserves Insulin Sensitivity. Frontiers in physiology 11, 696 (2020).

56. K. A. Levy, E. D. Weisz, T. A. Jongens, Loss of neurexin-1 in Drosophila melanogaster results in altered energy metabolism and increased seizure susceptibility. Human molecular genetics 31, 3422–3438 (2022).

57. L. Xu, Q. Feng, H. Deng, X. Zhang, H. Ni, M. Yao, Neurexin-2 is a potential regulator of inflammatory pain in the spinal dorsal horn of rats. Journal of cellular and molecular medicine 24, 13623–13633 (2020).

58. L. J. Simmons, M. C. Surles-Zeigler, Y. Li, G. D. Ford, G. D. Newman, B. D. Ford, Regulation of inflammatory responses by neuregulin-1 in brain ischemia and microglial cells in vitro involves the NF-kappa B pathway. Journal of neuroinflammation 13, 237 (2016).

59. X. Mao, M. T. Ou, S. S. Karuppagounder, T. I. Kam, X. Yin, Y. Xiong, P. Ge, G. E. Umanah, S. Brahmachari, J. H. Shin, H. C. Kang, J. Zhang, J. Xu, R. Chen, H. Park, S. A. Andrabi, S. U. Kang, R. A. Gonçalves, Y. Liang, S. Zhang, C. Qi, S. Lam, J. A. Keiler, J. Tyson, D. Kim, N. Panicker, S. P. Yun, C. J. Workman, D. A. Vignali, V. L. Dawson, H. S. Ko, T. M. Dawson, Pathological α-synuclein transmission initiated by binding lymphocyte-activation gene 3. Science (New York, N.Y.) 353, (2016).

60. D. Athauda, K. Maclagan, N. Budnik, L. Zampedri, S. Hibbert, I. Aviles-Olmos, K. Chowdhury, S. S. Skene, P. Limousin, T. Foltynie, Post hoc analysis of the Exenatide-PD trial-Factors that predict response. The European journal of neuroscience 49, 410–421 (2019).

61. R. Brauer, L. Wei, T. Ma, D. Athauda, C. Girges, N. Vijiaratnam, G. Auld, C. Whittlesea, I. Wong, T. Foltynie, Diabetes medications and risk of Parkinson’s disease: a cohort study of patients with diabetes. Brain : a journal of neurology 143, 3067–3076 (2020).

62. J. Kim, B. Tadros, Y. H. Liang, Y. Kim, C. Lasagna-Reeves, J. Y. Sonn, D. C. Chung, B. Hyman, D. M. Holtzman, H. Y. Zoghbi, TYK2 regulates tau levels, phosphorylation and aggregation in a tauopathy mouse model. Nature neuroscience 27, 2417–2429 (2024).

63. B. Vaidya, H. Kaur, P. Thapak, S. S. Sharma, J. N. Singh, Pharmacological Modulation of TRPM2 Channels via PARP Pathway Leads to Neuroprotection in MPTP-induced Parkinson’s Disease in Sprague Dawley Rats. Molecular neurobiology 59, 1528–1542 (2022).

64. Y. Kim, B. Vaidya, J. McInnes, H. Y. Zoghbi, Alpha-Synuclein Phosphomimetic Y39E and S129D Knock-In Mice Show Cytosolic Alpha-Synuclein Localization without Developing Neurodegeneration or Motor Deficits. eNeuro 12, (2025).

65. S. Kaye, A. Gold, D. Lin, M. Chen, J. Zhu, J. Gao, Hypercholesterolemia drives microglial dysfunction and weakens response to amyloid plaques. Experimental neurology 390, 115272 (2025).

66. Y. Li, A. G. Anderson, G. Qi, S. R. Wu, J. P. Revelli, Z. Liu, H. Y. Zoghbi, Early transcriptional signatures of MeCP2 positive and negative cells in Rett syndrome. bioRxiv : the preprint server for biology, (2025).

67. Chromium GEM-X Single Cell 5’ Reagent Kits v3. https://www.10xgenomics.com/support/universal-five-prime-gene-expression/documentation/steps/library-prep/chromium-gem-x-single-cell-5-v3-gene-expression-user-guide

